# Protective variant in PLCγ2 mitigates Alzheimer’s disease-associated pathologies via enhancing beneficial microglia functions

**DOI:** 10.1101/2024.09.12.612207

**Authors:** Mari Takalo, Heli Jeskanen, Taisia Rolova, Inka Kervinen, Marianna Hellén, Sami Heikkinen, Henna Koivisto, Kimmo Jokivarsi, Stephan A. Müller, Esa-Mikko Koivumäki, Petra Mäkinen, Sini-Pauliina Juopperi, Roosa-Maria Willman, Rosa Sinisalo, Dorit Hoffman, Henna Jäntti, Michael Peitz, Klaus Fließbach, Teemu Kuulasmaa, Teemu Natunen, Susanna Kemppainen, Pekka Poutiainen, Ville Leinonen, Tarja Malm, Henna Martiskainen, Alfredo Ramirez, Annakaisa Haapasalo, Stefan F. Lichtenthaler, Heikki Tanila, Christian Haass, Juha Rinne, Jari Koistinaho, Mikko Hiltunen

## Abstract

**Background:** PLCγ2-P522R (phospholipase C gamma 2, proline 522 to arginine) is a protective variant that reduces the risk for late onset Alzheimer’s disease. Recently, it was shown to decrease β-amyloid pathology in 5XFAD mouse model of AD. In this study, our goal was to investigate the protective functions of PLCγ2-P522R variant in a less aggressive mouse model of AD as well as to assess the underlying mechanisms at the molecular and cellular level using mouse and human microglia models.

**Methods:** The effects of the protective PLCγ2-P522R variant on microglia activation, AD-related β-amyloid and neuronal pathologies, as well as behavioral changes were investigated in PLCγ2-P522R knock-in mice crossbred with APP/PS1 AD model mice. Transcriptomic, proteomic, and functional studies were carried out in cultured and acutely isolated adult PLCγ2-P522R mouse microglia to study molecular mechanisms. Finally, microglia-like cell models generated from blood and skin biopsy samples of the PLCγ2-P522R variant carriers were employed to translate the key findings to human cells.

**Results:** Our results demonstrate that the PLCγ2-P522R variant reduced brain β-amyloid plaque burden of APP/PS1 mice. Simultaneously, PLCγ2-P522R variant increased non-proinflammatory microglia activation and microglia clustering around β-amyloid plaques, leading to reduced β-amyloid plaque-associated neuronal dystrophy. In cultured mouse primary microglia, PLCγ2-P522R variant decreased accumulation of large lipid droplets, reduced cell stress, and increased acute response to strong inflammatory stimuli. Transcriptomic and proteomic analyses in acutely isolated adult mouse microglia as well as in human monocyte-derived microglial cells showed that PLCγ2-P522R upregulates mitochondrial fatty acid oxidation and downregulates inflammatory/interferon signaling pathways. Accordingly, PLCγ2-P522R increased mitochondrial respiration in iPSC-derived microglial cells.

**Conclusions:** Together, these findings suggest that PLCγ2-P522R variant exerts protection against AD-associated β-amyloid and neuronal pathologies via enhancing microglial barrier formation around β-amyloid plaques and suppressing pro-inflammatory activation. Observed changes in fatty acid metabolism and mitochondrial flexibility as well as the downregulation of genes involved in inflammatory signaling pathways suggest that these protective effects of the PLCγ2-P522R variant are mediated through an anti-ageing mechanism.

## BACKGROUND

Phospholipase C gamma 2 (PLCγ2) is a hub enzyme regulating multiple cellular signaling pathways. PLCγ2 plays a crucial role in various cellular processes, including signal transduction, cell differentiation, cell proliferation, survival, and immune responses. Upon activation, PLCγ2 specifically converts phosphatidylinositol 4,5-bisphosphate (PIP2) into second messengers, inositol trisphosphate (IP3) and diacylglycerol (DAG). In turn, these trigger intracellular calcium release, activate downstream signaling pathways, and lead to various cellular responses. PLCγ2 is particularly important in immune cells, where it is activated by antigen binding to cell surface immune receptors (Bill & Vines, 2020).

In the brain, PLCγ2 is primarily expressed in microglia, the resident immune cells of the brain (Sims et al., 2017; Magno et al., 2019). Microglia play a key role in surveillance, tissue homeostasis, as well as response to injury, infection, and accumulation of toxic metabolites, including β-amyloid (Hansen et al., 2018). In Alzheimer’s disease (AD), activated microglia limit disease pathology via clearing and/or compacting β-amyloid as well as protecting vulnerable neurons from its toxic effects (Keren-Shaul et al., 2017). Conversely, extensive microglia activation promotes chronic neuroinflammation, which causes neuronal damage, and thus, is a major player in AD pathogenesis alongside with β-amyloid aggregation and hyperphosphorylated tau deposition (Hansen et al., 2018).

In microglia, PLCγ2 operates as a key downstream effector for Triggering receptor expressed on myeloid cells 2 (TREM2) and Toll-like receptors (TLR) (Andreone et al., 2020; Takalo et al., 2020). TREM2 is a microglia-specific lipid sensor, which upon ligand binding initiates an intracellular signaling cascade leading to changes in gene transcription, cell proliferation, cell metabolism, phagocytosis, and survival. Genetic depletion of either TREM2 or PLCγ2 similarly impair lipid processing, leading to compromised mitochondrial energy capacity in human iPSC-derived microglia (iMGL) (Andreone et al., 2020; Van Lengerich et al., 2023). Furthermore, buildup of lipids into specific primary storage units, lipid droplets (LD), in aging and during β-amyloid accumulation drives the dysfunctional and neurotoxic phenotype in microglia (Haney et al., 2024; Marschallinger et al., 2020).

Recent genetic studies support a strong link between late onset AD (LOAD) and *PLCG2,* the gene encoding PLCγ2. A rare gain-of-function variant (rs72824905, C>G, p.P522R) was shown to reduce the risk of AD (Sims et al., 2017). The same variant was suggested to have beneficial effects on brain health also beyond AD. The PLCγ2-P522R variant was shown to reduce the risk of dementia with Lewy bodies and frontotemporal dementia, and the carriers of the protective variant were enriched among the oldest (+100 years) individuals with well-preserved cognitive functions (Van Der Lee et al., 2019). Furthermore, PLCγ2-P522R variant slowed the rate of cognitive decline in patients with mild cognitive impairment, suggesting that the PLCγ2-P522R variant has universally beneficial effects on cognition and brain health upon aging (Kleineidam et al., 2020). In addition, other gain– and loss-of-function variants in *PLCγ2* have recently been linked to the decreased or increased risk of AD, respectively, which further supports the key role for PLCγ2 in AD and beyond (Kunkle, et al., 2019; Bellenguez et al., 2017; Coulon et al., 2024, preprint; Bellenquez et al., 2022).

PLCγ2-P522R is a functional hypermorph, which mildly increases PLCγ2 enzyme activity (Andreone et al., 2020; Magno et al., 2019; Takalo et al., 2020). We have previously shown that PLCγ2-P522R variant increases microglia activation in adult C57BL/6J mice and improves acute inflammatory response and survival in mouse bone marrow-derived macrophages upon treatment with lipopolysaccharide (LPS) (Takalo et al., 2020). Recently, PLCγ2-P522R was shown to enhance microglia-mediated phagocytosis of β-amyloid, leading to more compacted plaque morphology, reduced neuronal damage, and enhanced working memory in 5XFAD mouse model of AD (Tsai et al., 2023). Importantly, very similar effects have been reported for agonistic TREM2 antibodies, which are designed to boost positive microglia functions in AD patients (Schlepckow et al., 2020; Van Lengerich et al., 2023). Together these studies emphasize the importance of TREM2-PLCγ2-driven microglia functions in restricting AD-related pathologies and highlight PLCγ2 as a plausible disease-modifying therapeutic target in AD.

Despite promising results, detailed cellular mechanisms underlying the activation of PLCγ2 are still very limited, and data gathered from different models are needed. This study aimed to investigate the effects of the protective PLCγ2-P522R variant on microglial responses and AD-related β-amyloid and neuronal pathologies as well as behavioral changes for the first time in a less aggressive APP/PS1 AD mouse model. Importantly, mouse primary microglia as well as human microglial models were utilized for comprehensive assessment of molecular and cellular mechanisms underlying the protective effects of the PLCγ2-P522R variant.

## MATERIALS AND METHODS

### Study design

In this study APPswe/PS1dE9 transgenic mouse line (later simply APP/PS1) (Jankowsky et al., 2004) was crossbred with PLCγ2-P522R knock in (KI) mouse line (Takalo et al., 2020) to investigate the effects of the PLCγ2-P522R variant on AD-related β-amyloid and associated neuronal pathologies, on microglia activation, and behavioral changes. Mice at the age of 7 and 13-months were subjected to TSPO-PET imaging with [18F]FEPPA ligand to monitor microglia activation (Takalo et al., 2020). Behavioral testing was conducted for mice at the age of 12 months. These mice were then sacrificed, and half of the brain was cut into coronal sections and used for immunohistochemical analyses while the other half was dissected into regional blocks and utilized for biochemical assays. The whole brain of a subset of mice was used for CD11b+ microglia isolation and utilized for RNA sequencing studies. Unless otherwise informed, female mice were used for all experiments conducted in APP/PS1.

In addition, primary microglia isolated from 0-3 days old PLCγ2-P522R KI mouse pups were used for mechanistic studies. Primary microglia isolated from TREM2 knock-out (KO) and wildtype (WT) mice were used as controls. Lipid accumulation within microglia was analyzed upon LPS and myelin treatments. To assess functional changes associated with PLCγ2-P522R, mitochondrial respiration and glycolysis were analyzed in LPS and myelin-treated cells. Cellular response to acute inflammatory stimuli was assessed at 3 h, 24 h and 48 h time points after LPS treatment and cell stress was evaluated by measuring production of lactate dehydrogenase (LDH) and reactive oxygen species (ROS). Molecular targets and pathways related to the PLCγ2-P522R-driven functional phenotype were analyzed using global transcriptomic and proteomic analyses in CD11b+ microglia acutely isolated from 13-month-old PLCG2-P522R KI and WT mice.

Finally, transcriptomic analyses were conducted in blood monocyte-derived microglial cells (MDMi) from PLCγ2-P522R variant carriers and matched controls to study how well the findings are translated to human cells. Induced pluripotent stem cell-derived microglia (iMGL) homozygous for PLCγ2-P522R variant having either *APOE33* or *APOE44* genetic background and isogenic control lines were used to functionally assess the key findings.

### Mouse lines

The APP/PS1 line in C57Bl/6J background was used as a model for AD (Jankowsky et al., 2004). This mouse line co-expresses in a single transgene a chimeric mouse/human APP695 harboring the Swedish K670M/N671L mutations (Mo/Hu APPswe) and human PS1 with the exon-9 deletion (PS1dE9) under mouse prion protein promoter. This line was crossbred with a PLCG2-P522R KI mouse line generated using CRISPR/Cas9-assisted gene editing in C57Bl/6J background as we have reported before (Takalo et al., 2020). WT (C57Bl/6J) and TREM2 KO (The Jackson Laboratory, stock no. 027197) mice were used as controls in *in vitro* experiments. The mice were kept in an environment with constant temperature, 22 ± 1 °C, humidity 50 – 60 %, lights on 07:00-19:00, with food and water available *ad libitum*. All mouse studies were carried out in accordance with the guidelines of the European Community Council Directives 86/609/EEC and approved by the Animal Experiment Board of Finland/Regional State Administrative Agency and Project Authorisation Board (ESAVI 21203–2019, EKS-004-2019).

### Mouse genotyping

Genomic DNA was purified from ear biopsies by incubating the samples with 50 mM NaOH for 1 h at 95 °C and neutralizing with 1 M Tris, pH 8.0. Genotyping of PLCγ2-P522R locus was carried out with SNP genotyping. Ten ng DNA was mixed with TaqMan Genotyping master mix (Applied Biosystems) and TaqMan SNP genotyping assay (Applied biosystems) targeted to trinucleotide variation leading to substitution of proline 522 with arginine (Takalo et al., 2020). Reaction mixture was amplified for 10 s at 95 °C, 15 s at 95 °C, and 60 s at 60 °C for 40 cycles and measured with LightCycler® 480 II (Roche).

For confirming the expression of *APP* and *PSEN1* genes, locus harboring human *PSEN1* and mouse prion protein promoter were PCR amplified with 5’-CCTCTTTGTGACTATGTGGACTGATGTCGG-3’, 5’-GTGGATACCCCCTCCCCCAGCCTAGACC-3’, and 5’-CAGGTGGTGGAGCAAGATG-3’ primers using touchdown PCR with the following program: 3 min at 94 °C; 30 s at 94 °C, 1 min at 62-58 °C, and 2 min at 72 °C, for two cycles at each temperature; 30 s at 94 °C, 1 min at 57 °C, and 2 min at 72 °C for 25 cycles; 5 min at 72 °C. 1300 bp (*PSEN1*) and 750 bp (internal control) bands were detected with agarose gel electrophoresis.

TREM2 KO mouse genotypes were confirmed using touchdown PCR with primers 5’-TCAGGGAGTCAGTCATTAACCA-3’, 5’-AGTGCTTCAAGGCGTCATAAGT-3’, and 5’-CAATAAGACCTGGCACAAGGA-3’. 396 bp (KO), 254 bp and 396 bp (heterozygote) and 254 bp (WT) bands were detected with agarose gel electrophoresis.

### Behavioral studies

Mice at the age of 12 months were subjected to a behavioral test battery. Mice were weighed before starting the testing.

Spontaneous explorative activity was assessed by an automated activity monitor (TruScan, Coulbourn Instruments, Whitehall,PA, USA) based on infrared photobeam detection. The system consisted of an observation cage with white plastic walls (26 cm × 26 cm × 39 cm) and two frames of photo detectors for monitoring horizontal and vertical activity. The test cage was cleaned with 70% ethanol before each mouse to avoid odor traces. The following parameters were measured during a 10-min session: ambulatory distance (gross horizontal locomotion), rearing time, and time in the area center vs. periphery.

A light-dark box with an electric grid floor in the dark compartment was used to test spontaneous fear toward a lit compartment and for foot-shock reinforced passive avoidance. On day 1, the mouse was first placed in the lit compartment with free access to the dark compartment through a narrow opening, and the time spent in the light vs. dark compartment was recorded for 5 min. Thereafter, the mouse was placed again in the lit compartment for 30 s with the entrance to the dark compartment closed. Then the slide door separating the compartments was opened. As soon as the mouse entered the dark side (cut-off 3 min), the slide door was closed and a mild foot-shock delivered (2 x 2 s at 0.30 mA). The mouse was then returned to its home cage. On day 3 (48 h later), the mouse was again placed in the lit compartment with the slide door open, and the time to enter the dark side was recorded as an index for fear memory. Finally, the mouse was confined to the dark compartment, and the pain threshold to electric foot shock was determined by gradually increasing the current on the grid floor until the mouse reacted.

Motor coordination and balance was tested in an automated Rotarod device (Uno Bacile, Italy). Each mouse was first familiarized with the rotating rod twice for 2 min by letting it rotate at a constant speed of 4 rpm. On a separate day, the mouse was tested twice with the speed of the rod increasing stepwise from 4 to 40 rpm. The cut-off time was 6 min. The trial with longer latency to fall was recorded.

Pain threshold was tested on a hotplate with a surface temperature of 47 and 51°C. The time to lick a paw or to jump was recorded until a cutoff time of 60 s. Each mouse was tested three times at both temperatures (lower first), with 1 h between the test sessions. The average reaction time at each temperature was recorded.

Spatial learning and memory were assessed in the Morris swim navigation task. The test was conducted in a white circular wading pool (diameter 120 cm) with a transparent submerged escape platform (14 cm × 14 cm). The pool was open to landmarks in the room. The water temperature was 20 ± 0.5 °C. The testing was preceded by two practice days with a guiding alley to the platform. During the acquisition phase (days 1– 5), the location of the hidden platform was kept constant and the start position varied between four different locations at the pool edge. Each mouse was placed in the water with its nose pointing towards the pool wall. If the mouse failed to find the escape platform within 60 s, it was placed on the platform for 10 s by the experimenter. The acquisition phase consisted of five daily trials with a 10 min inter-trial-interval. On day 5, the search bias was tested in a 60-s probe trial (the 5th trial) without the platform. The mouse was video tracked, and the video analysis program (Ethovision, Noldus, The Netherlands) calculated the escape latency, swim speed, path length and time in the pool periphery (10 cm from the wall) and in the platform zone (diameter 30 cm).

### PET-imaging

[F18]FEPPA-PET-imaging was conducted on female and male mice at the age of 7 and 13 months. [18F]FEPPA is a specific positron emission tomography (PET) ligand for the mitochondrial translocator protein 18-kDa (TSPO) that is highly expressed in activated microglia (Wilson et al., 2008). Mice were anesthetized with isoflurane (1.5% with N2/O2 70%/30% through nose cone) and placed on a heated animal holder (Équipement Vétérinaire Minerve, Esternay, France) on the scanner bed in a prone position and secured with tape to prevent movement during scanning. For the 7-month timepoint the mice were imaged using a dedicated PET scanner (Inveon DPET, Siemens Healthcare) and immediately afterwards with CT (Flex SPECT/CT, Gamma Medica, Inc.) for anatomical reference images using the same animal holder. For the 13-month timepoint imaging was done using a preclinical PET/MRI scanner (MR Solutions Ltd, Surrey, UK). Change of scanners was due to unexpected technical problems with the dedicated PET scanner. Dynamic imaging of 70 min was started at the time of the administration of the activity. 18F-FEPPA (8.0 ± 2.0 MBq) was injected as a slow bolus over a period of 30 s through the tail vein. PET data were gathered in list-mode form, and corrected for deadtime, randoms, scatter and attenuation. Data were reconstructed using the scanner manufacturers’ software with 2D-(7 mo) or 3D-(13 mo) ordered-subsets expectation maximization (OSEM) algorithm. Regions of interest (ROIs) were drawn for whole brain (excluding cerebellum and olfactory bulb), frontal cortex, pons, hippocampus and cerebellum. The ROIs were drawn using Carimas 2.10 software (Turku PET Centre, Finland). Final uptake values (percent of injected dose per ml of tissue) were the average values from 20 to 70 minutes from the ligand injection.

### Tissue collection and sample preparation

At the age of 13 months, the mice were terminally anesthetized with 60 mg/ml pentobarbital, after which they were transcardially perfused with ice-cold phosphate-buffered saline (DPBS, #17-512F, BioNordica) for 5 min to rinse blood from the brain. The brain was quickly removed and placed on ice. One brain hemisphere was immersion fixed in 4 % paraformaldehyde (PFA), followed by incubation in 30 % sucrose overnight. Fixed tissue was stored in antifreeze at – 20 °C until it was cut into 35 µm thick coronal sections using a freezing sliding microtome.

The other hemisphere was dissected on ice into temporo-occipital cortex and hippocampus and snap-frozen in liquid nitrogen. Tissue pieces were mechanically homogenized in 300 µl of ice-cold DPBS. Fifty µl of the lysate was mixed with 500 µl of TRI-reagent (#T9424, Sigma-Aldrich) and used for RNA extraction (below). The remaining lysate was supplemented with 1:100 HALT™ EDTA-free protease (#87785, Thermo Scientific) and HALT™ phosphatase (#78420, Thermo Scientific) inhibitor cocktails. Proteins were extracted by incubating the lysate on ice for 30 min with Tissue Protein Extraction Reagent (T-PER, #78510, Thermo Scientific) and collecting supernatant after centrifuging for 10 min at 10,000 x g at + 4 °C. Fraction of PBS lysate was ultracentrifuged at 100,000 × g for 2 h at + 4 °C and supernatant containing the soluble fraction was collected to clean tubes. Pellets containing the insoluble fraction were solubilized in 5 M Guanidine-HCl by vortexing for 3 h at room temperature. Samples were centrifuged at 10,000 × g for 10 s. Protein concentrations were measured using BCA protein assay kit (#23225, Pierce).

### Western blotting

Total protein lysates (15–20 μg) were supplemented with NuPAGE LDS Sample Buffer (#NP0007, Invitrogen) including β-mercaptoethanol and separated with SDS-PAGE using NuPAGE 4–12 % Bis-Tris Midi Protein Gels (#WG1402BOX, Invitrogen). Proteins were subsequently transferred to polyvinylidene difluoride (PVDF) membranes (#IB24001, Invitrogen) using the iBlot 2 Dry Blotting System (Invitrogen). Unspecific antibody binding was blocked by incubating the blots in 5 % non-fat milk or 5 % bovine serum albumin (BSA, #A9647, Sigma-Aldrich) in 1x Tris-buffered saline with 0.1 % Tween 20 (TBST) for 1 h at room temperature. Proteins were detected from the blots using the following primary antibodies diluted in 1x TBST and incubated overnight at + 4 °C: mouse anti-β-amyloid 1:1000 (6E10, #SIG-39320, BioLegend), rabbit anti-sAPPβ 1:100 (#18957, IBL America), mouse anti-APP N-terminal 1:1000 (22C11, #MAB348, Merck), rabbit anti-APP C-terminal 1:1000 (#A8717, Merck), and mouse anti-GAPDH 1:10000 (#ab8226, Abcam). Subsequently, blots were incubated with sheep anti-mouse 1:5000 (#NA931V, Cytiva) or donkey anti-rabbit 1:5000 (#NA934V, Cytiva) horseradish peroxidase (HRP)-conjugated secondary antibodies in 1x TBST and for 1 h at room temperature. Enhanced chemiluminescence Prime (#RPN2232, Cytiva) and Select (#RPN2235, Cytiva) Western blotting detection reagents were used to detect the protein bands. SeeBlue Plus2 Prestained Standard (#LC5925, Invitrogen) was used to estimate the size of the detected bands. Blots were imaged with the Chemidoc MP system (Bio-Rad) and images were quantified using the ImageLab (Bio-Rad) software. Results are shown as percent of A+/P^wt/wt^ group.

### Aβ40, Aβ42, sTREM2, and APOE ELISA

Aβ40 and Aβ42 levels in the soluble and insoluble lysates were determined with Amyloid β (40) Human/Rat (#294-64701, FUJIFILM, Wako) and High Sensitive Amyloid β (42) Human/Rat (#292-64501, FUJIFILM, Wako) ELISA Kits following manufacturer’s instructions. Aβ levels were normalized to the total protein concentration within each sample.

Soluble and insoluble APOE were measured from the respective lysates using Apolipoprotein E (APOE) Mouse ELISA Kit (#ELK2007, Gentaur) according to kit instructions. APOE levels were normalized to the total protein concentration within the same sample.

For the sTREM2 measurement from the respective lysates, 96-well Costar Assay Plate (9018, Corning) was coated with 500 ng/ml Human/Mouse TREM2 antibody (#MAB17291-100, R&D Systems) in 0.5 M Carbonated-Bicarbonate buffer (pH 9.6, #C3041, Sigma-Aldrich) and incubated at

+ 4 °C overnight. Wells were washed 3 times with 0.05 % Tween20 (#93773-250G, Sigma-Aldrich) in DPBS and blocked by 1 % Block Ace (#BUF029, Bio-Rad) in DPBS for 4 h at room temperature. Wells were washed once before adding Recombinant Mouse TREM2 Protein in concentrations 12.21-781.25 pg/ml (#50149-M08H, Sino Biological Inc.) and samples diluted in Assay buffer (1 % BSA (#A9647, Sigma-Aldrich), 0.05 % Tween20 in DPBS). Standards and samples were incubated at + 4 °C overnight. Wells were washed five times following incubation of 66.67 pg/ml Mouse TREM2 biotinylated antibody (#BAF1729, R&D Systems) for 1 h at room temperature after which wells were washed 3 times. Streptavidin Poly-HRP40 (#20102011, Fitzgerald) diluted 1:3000 in assay buffer was added and incubated for 1 h at room temperature covered from light after which wells were washed 5 times. 3,3′,5,5′-Tetramethylbenzidine Liquid Substrate (TMB, Super Slow, #T5569, Sigma-Aldrich) was added and incubated for 15 min at room temperature. The reaction was stopped with 1 M H_3_PO_4_ and absorbances were read with plate reader (Tecan Infinite, 450 nm).

### Mesoscale inflammatory marker assay

Inflammatory cytokines and transforming growth factor beta (TGF-β) 1-3 in the cortical and hippocampal lysates were determined with MSD mouse V-plex Proinflammatory panel 1 (#K15048, Mesoscale) and U-plex TGFβ combo panel (#K15242K, Mesoscale), respectively, according to kits instructions. The results were normalized to the total protein concentration in each sample.

### Immunohistochemistry of mouse brain sections

Three to four brain sections between bregma − 3.1 mm and − 3.5 mm coronal planes according to the Paxinos and Franklin’s the Mouse Brain in Stereotaxic Coordinates (2001) were selected from each mouse for every staining. Before staining, the slices were kept in sodium phosphate buffer overnight at room temperature. Next day, the slices were pre-treated in 0.05 M citrate solution (pH 6.0) for 30 min at 80 °C and cooled down in sodium phosphate buffer for 20 min. To visualize β-amyloid plaques, the sections were incubated with 0.1 mM X-34 (#SML1954, Sigma-Aldrich) for 1 h at room temperature in a solution containing 60 % DPBS and 40 % EtOH. Slices were then rinsed three times with 60 % DPBS + 40 % EtOH followed by washing with TBST three times for 5 min. Non-specific antibody binding was blocked with 3 % BSA in TBST, after which the sections were incubated with following primary antibodies at 4 °C overnight: rabbit anti-β-amyloid 1:1000 (6E10, #SIG-39320, BioLegend) to detect diffuse β-amyloid, mouse anti-APP N-terminal 1:2000 (22C11, #MAB348, Merck) to visualize dystrophic neurites, rabbit anti-IBA1 1:1000 (#019–19,741, FUJIFILM Wako) to detect microglia, rabbit anti-GFAP 1:500 (#Z0334, Dako) to detect astrocytes, and goat anti-APOE 1:1000 (#178479, Merck). After washing three times 5 min with TBST at room temperature, the sections were incubated with appropriate fluorescent secondary antibodies Alexa Fluor™ 568 Donkey anti-Mouse (#A-10037, Thermo Scientific), Alexa Fluor™ 488 Goat anti-rabbit (#A-11008, Invitrogen), Alexa Fluor™ 488 Donkey anti-rabbit (#A-21206, Thermo Scientific), Alexa Fluor™ 568 Goat anti-rabbit (#A-11036, Invitrogen), and Alexa Fluor™ 568 Donkey anti-goat (#A-11057, Thermo Scientific) for 1 h at room temperature. The sections were again washed three times 5 min and then mounted with Vectashiled HardSet Antifade Mounting Medium (#H-1400, Vector Laboratories) on gelatin-coated slides. Control sections without the primary antibodies were processed simultaneously.

### Fluorescence microscopy and image analysis

For X-34 positive β-amyloid plaque coverage analysis, the slices were imaged with a Leica Thunder Imager 3D Tissue slide scanner equipped with K5-14400955 camera and HC PL FLUOTAR 20x (NA 0.5) DRY objective (Leica Microsystems). Z-stack images were taken with 19 layers with a total distance of 27 μm. The whole entorhinal cortex and hippocampus were scanned, stitched, and subjected to automated Thunder image processing computational clearance. All stainings including more than one fluorophore were imaged with a Zeiss Axio Observer inverted microscope with a Zeiss LSM 800 confocal module (Carl Zeiss Microimaging GmbH) using 20 × (NA 0.5) objective. Plan-Apochromat 63 × (NA 1.4) objective was used for imaging 6E10 and IBA1 for Imaris-based colocalization analysis. Three 1024 x 1024-pixel fields from both brain areas were taken using Z-stack imaging method including 15-19 layers with a total distance of 22.5-27 μm. All laser, light, and detector settings were kept constant for all samples within each immunostaining. Images were taken by investigators blinded to the genotypes.

Quantitative image analysis was carried out with Fiji (Image J, version 1.53c) software. Maximum intensity projections were generated for each channel and rolling ball algorithm (radius 50) and Gaussian blurring (sigma 2) were applied to subtract background and remove noise, respectively. The following automatic thresholding methods were used for creating a mask for each staining: Otsu for X-34, Moments for 22C11 and IBA1, and Li for GFAP and APOE. For Leica Thunder Imager 3D Tissue slide scanner images, entorhinal cortex and hippocampal ROIs were manually drawn for each image. Subsequently, X-34 positive β-amyloid plaque area and count in the entorhinal cortex and hippocampus were normalized to the area of respective ROI. For the analysis of IBA1, GFAP, 22C11, and APOE within and surrounding β-amyloid plaque area, the outlines of β-amyloid plaques were first determined from the X-34 staining, then IBA1, GFAP, 22C11, and APOE+ areas were analyzed within 0-40 µm, 0-40 µm, 0-14 µm, and 0-10 µm from the plaque outline, respectively (**Figure S1A-D**). The smallest area covering most of the IBA1, GFAP, 22C11 and APOE signal surrounding β-amyloid plaque was used for choosing the width of the analyzed area. Image pre-processing and thresholding methods were kept constant between the samples within each staining. No signal in the negative control samples was detected. The ZEN 2012 software Blue edition (Carl Zeiss Microimaging GmbH) was used for visualization purposes.

Imaris (10.1.1) software was used in quantitative image analysis for 6E10 and IBA1 colocalization. A surface for 6E10 and IBA1 was created using the surface tool. A Gaussian filter was applied using smoothing function and the surface detail value was set to 0.198 µm. Absolute Intensity Thresholding method was used for surface creation with set treshold of 3.8 for 6E10 and 12 for IBA1. Surfaces below 600 voxels in IBA1 channel and 70.7 voxels in 6E10 channel were filtered out from the analysis to exclude small particles. A mask was created from the 6E10 channel using the created IBA1 surface. Voxel intensity outside IBA1 surface was set as 0 for this mask. A new surface was created from the mask using the same settings as creating the 6E10 surfaces previously. The new surface represented the overlapping signal of 6E10 IBA1. All image analysis parameters were kept the same for all the samples. The data are presented as 6E10 volume inside IBA1 surface as percent of total 6E10 volume. All image analyses were done by an investigator blinded to the genetic background of the mice.

### Acute CD11b+ microglia isolation

Mice at 13 months of age were anesthetized with 60 mg/ml pentobarbital and briefly perfused with ice cold DPBS to get rid of blood cells. Mice were then decapitated, and brains were placed in ice cold Hank’s Balanced Salt Solution without Ca^2+^ or Mg^2+^ (HBSS, #14175129, Gibco). Brains were mechanically dissociated into small pieces and further into single cell suspension using enzymatic digestion. Cell isolation was done according to instructions of Adult Brain Dissociation Kit (#130-107-677, Miltenyi) and separation of CD11b+ microglia following CD11b MicroBeads, human and mouse (#130-093-634, Miltenyi) protocol. Magnetic separation was done using QuadroMACS Separator (#130-090-976, Miltenyi) with LS Columns (#130-042-401, Miltenyi). Cell pellet was washed twice with DPBS and immediately used for RNA extraction (below) or stored as a dry pellet at –80 °C for proteomic analysis (below).

### Mouse primary microglia cultures

Homozygous PLCγ2-P522R KI and WT mouse pups (P0-P3) were decapitated and brains taken out on 6-cm petri dishes in ice-cold HBSS (#24020091, Thermo Fisher). Meninges were removed under the microscope and the brains were washed three times with ice-cold HBSS. For tissue lysis, 0.25 % Trypsin (#15090046, Gibco) was added, and the brains were incubated for 10 minutes at 37 °C in cell culture incubator. Reaction was stopped by adding culture medium including DMEM with 4.5 g/l glucose, without L-glutamine (#ECB7501L, BioNordica), 10 % (v/v) heat-inactivated fetal bovine serum (FBS, #10500-064, Gibco), 2 mM L-glutamine (#BE17-605E, Lonza Bioscience), and 1 % (v/v) penicillin-streptomycin (#DE17-602E, Lonza Bioscience). Next, 400 µl of 5 mg/ml Deoxyribonuclease I (#D4527-40KU, Sigma) was added, and cells were separated by pipetting carefully. Cells were then centrifuged for 10 min at 800 x g at room temperature and resuspended in culture medium. Cells were cultured on T-75 cell culture flasks coated with 0.02 % Poly-L-lysine hydrobromide (PLL, #P6407, Sigma), until microglia were shaken off a week later and then every 3-4 days.

### Myelin preparation

Myelin was isolated from the brains of adult male C57BL/6J mice by a discontinuous sucrose gradient according to a protocol described as in (Larocca & Norton, 2006; Safaiyan et al., 2016). Briefly, the brain tissue was homogenized in a solution containing 10 mM HEPES, 5 mM EDTA, 0.32 M sucrose and protease inhibitor. The homogenate was layered on top of 0.85 M sucrose solution prepared in the HEPES-EDTA buffer and centrifuged at 75,000 x g for 30 min with low acceleration and deceleration using Beckman Ultracentrifuge with Ti 50.2 rotor. The white myelin layer was collected from the interface, suspended in sterile water, and centrifuged at 75,000 g for 15 min. The pellet was purified twice by a hypo-osmotic shock using ice-cold water and centrifugation at 12,000 g for 10 min. After that, the sucrose gradient and the hypo-osmotic shocks were repeated to purify the myelin fraction. At the end, the myelin pellet was resuspended in HEPES-EDTA buffer and stored at −80 °C. The concentration of the myelin was determined using BCA Protein Assay Kit (#23225, Pierce).

Extracted myelin was labeled with pHrodo™ iFL Red Microscale Protein Labeling Kit (Thermo Fisher Scientific, P36014) by adding 25 µl dye per 1 mg of myelin and incubating for 60 min at room temperature. The myelin was then pelleted by centrifuging 10 min, 15 000 x g and washed twice with 1x DPBS to remove the excess dye. pHrodo-labeled myelin was resuspended in 1x DPBS in concentration of 1 mg/ml and stored as aliquots at –80 °C.

### Study subjects, MDMi preparation and analysis

Cognitively healthy PLCγ2-P522R monoallelic (GC, *APOE33,* age 68 ± 6.7, n=7) carriers and matched controls (CC, *APOE33,* age 70 ± 5.5, n=7) were recruited during 2022-2024 through Auria Biobank based on pre-existing genome data returned to biobank from FinnGen (Kurki et al., 2023). Blood samples for monocyte and DNA isolation were collected from the study subjects following written informed consent from each participant.

To extract peripheral blood mononuclear cells (PBMCs), 60-100 ml peripheral venous blood was collected from the participants and processed within 24 h of sample collection. PBMCs were extracted by gradient centrifugation over Ficoll-Paque PLUS (#17-1440-02, Cytiva) in SepMate-50 tubes (#85450, Stemcell Technologies). Positive selection of monocytes from PBMCs was done with human CD14 MicroBeads (#130-050-201, Miltenyi Biotec) and magnet-activated cell sorting.

Monocytes were plated at the density of 1 million cells per well onto 12-well plates. Monocyte differentiation into MDMis was done according to a previously published protocol (K. J. Ryan et al., 2017) with modifications described in (Martiskainen et al., 2024, preprint). After MDMi differentiation for 12 days in vitro, conditioned medium was replaced with fresh MDMi culture medium or with culture medium containing myelin (25 µg/ml) or LPS (200 ng/ml, O26:B6, L5543, Sigma Aldrich), and cells were cultured for 24 h prior to sample collection. For RNA extraction, cells were washed three times with ice-cold PBS and stored in TRI Reagent (#T9424 Sigma-Aldrich). RNA extraction protocol is described below.

### iMGL preparation

The original iPSC line used in this study carrying APOE44 and control PLCG2 variants CC/APOE44 (UKBi011-A, https://hpscreg.eu/cell-line/UKBi011-A) was described previously (Peitz et al., 2018). The iPSC line CC/APOE33 (UKBi011-A3, https://hpscreg.eu/cell-line/UKBi011-A-3) was generated from the original line using CRISPR/Cas9 as described (Schmid et al., 2021). For the generation of homozygous PLCγ2-P522R lines on APOE33 (GG/APOE33) and APOE44 (GG/APOE44) backgrounds, the gRNA CCCCAAAATGTAGTTCTGTA was used. The sequence of the ssODN used as HDR donor was GTGAGACAGAAGGACCTGTCTAGTGATGCTGGGGTTTGGTCCAAGGCTTTCAGAAACCCCTCCTCTCTTTGC GGCCCAGGATATACG<u>CCCGACGGAGCTACATTTTGGGG</u>AGAAATGGTTCCACAAG.

Human iPSC lines were cultured in Essential 8 (E8) medium (#A1517001, Gibco) on Matrigel (#356231, Corning)-coated 3.5 cm dishes (#83.3900.500, Sarstedt) at 37 °C 5% CO_2_ and splitted with 0.5 mM EDTA every 4–5 days. For iMGL differentiation from iPSCs, a protocol described previously was used (Abud et al., 2017; Koskuvi et al., 2024; McQuade et al., 2018). Briefly, iPSC colonies were detached using ReLeSR reagent (#5872, STEMCELL Technologies) and plated at density 3-6 colonies per cm^2^ on Matrigel-coated 6-well plates (#356231, Corning) in E8 medium with 5 µM ROCK inhibitor (#Y-27632, Merck). On the following day, the differentiation into hematopoietic stem cells was started using the commercial STEMdiff Hematopoietic kit (#05310, STEMCELL Technologies). After 11-13 days of differentiation the floating hematopoietic progenitors were collected and plated at density 7000-8000 cells per cm^2^ on new Matrigel coated 6-well plates in microglial differentiation medium consisting of DMEM/F12 (#21331020, Gibco), 2× insulin-transferrin-selenite (#41400045, Invitrogen), 2× B27 (#17504044, Invitrogen), 0.5× N2, 1× glutamax (#35050-038, Invitrogen), 1× non-essential amino acids (#11140050, Invitrogen), 400 μM monothioglycerol (#M1753, Sigma), 5 μg/mL human insulin (#19278, Sigma), 100 ng/mL human IL-34 (#200-34, PeproTech), 50 ng/mL human TGF-β1 (#100-21, PeproTech), and 25 ng/mL human M-CSF (#300-25, PeproTech). The cells were grown for 27 days, and fresh medium was added every other day. During the last 4 days of culture the microglial maturation was promoted by adding 100 ng/mL human CD200 (#77002, Biolegend) and 100 ng/mL human CX3CL1 (#300-31, PeproTech) to the cells.

### Lipid droplet analysis

For analyzing lipid accumulation in mouse primary microglia, 50 000 cells were plated onto a 48-well plate on top of glass cover slips pre-coated with 0.02 % PLL in microglia culture medium (above) with reduced (5 %) FBS. Next day, cells were treated with 5 µg/ml LPS (#L5543, Sigma-Aldrich) for 24 h or with 30 µg/ml myelin for 48 h. Control wells were left untreated. Cells were then washed three times with warm DPBS and stained with 2 µM BODIPY 493/503 (#D3922, Invitrogen) for 15 min in cell culture incubator. Cells were rinsed several times with DPBS, fixed with 4 % PFA for 20 min at room temperature and mounted on microscope glasses with Vectashield Hard Set containing Phalloidin (#H-1699, Vector Laboratories).

Images of primary microglia stained with BODIPY 493/503 to visualize neutral lipid-containing lipid droplets (LD) and phalloidin to visualize actin cytoskeleton (cell border) were acquired with Zeiss Axio Observer inverted microscope with a Zeiss LSM 700 confocal module using 20 × (NA 0.5) objective. Five to ten 1024 x 1024-pixel fields per cover slip were obtained. All laser and detector settings were kept constant for all samples. Imaging was done by investigator blinded to the genetic background and treatment of the cells.

LDs were analyzed using (Image J, version 1.53c). LDs were segmented with Intermodes automatic threshold method. Outliers with radius 3 and threshold 50 were removed from the mask, after which particles bigger than 100 pixels were included in the particle analysis. number of LD-positive cells were manually counted. Analysis was done by an investigator blinded to the genetic background and treatment of the cells. Image pre-processing and thresholding methods were kept constant throughout the analysis. The ZEN 2012 software Blue edition (Carl Zeiss Microimaging GmbH) was used for visualization purposes. The % of LD-positive cells of all cells and average LD size (µm^2^) per imaged field are shown. Number of analyzed images are WT untreated=11, LPS=22, myelin=42 and KI PLCγ2-P522R untreated=6, LPS=12, myelin=35.

### Seahorse assay for mitochondrial and glycolytic energy metabolism

Mouse primary microglia were plated (25000 cells/well) on a Seahorse XF96 Cell Culture Microplate (#101085-004, Agilent) as 4-6 technical replicates per genotype and treatment in each experiment. On the next day, cells were treated with 200 ng/ml LPS (#L5543, Sigma-Aldrich), 25 µg/ml myelin, or left untreated. After 24 h, The Cell Mito Stress test or Cell Glycolysis Stress test was performed using assay parameters provided by Agilent. On the day of the experiment, medium was changed to Seahorse XF DMEM medium (#103575-100, Agilent) supplemented with 10 mM Seahorse XF glucose solution (#103577-100, Agilent), 2 mM Seahorse XF L-glutamine solution (#103578-100, Agilent), and 1 mM sodium pyruvate (#P5280-25G, Sigma). For the Cell Glycolysis Stress test, only glutamine was added to the Seahorse medium. Cells were kept in CO_2_-free incubator for 1 h prior to starting the experiment. For the Cell Mito Stress test, final concentration of 2 µM carbonyl cyanide-4-(trifluoromethoxy)phenylhydrazone (FCCP, #C2920, Sigma-Aldrich), 1 µM oligomycin (#75351, Sigma-Aldrich), and a mixture of 1 µM antimycin A and 1 µM rotenone (#A8674, Sigma-Aldrich) were used. Changes in oxygen consumption rate (OCR) in response to injections were measured with Seahorse XFe96 analyzer (Agilent). For the Cell Glycolysis stress test, final concentrations of 10 mM Seahorse XF glucose solution (#103577-100, Agilent), 1 µM oligomycin, and 50 mM 2-Deoxy-D-glucose (2-DG, #D6134-5G, Sigma-Aldrich) were used and extracellular acidification rate (ECAR) measured in response to the injections. Following the assay, cells were stained using 5 µM Vybrant DyeCycle Green Stain (#V35004, Thermo Fisher) and images were taken with 4x objective using brightfield and green fluorescence channel with IncuCyte S3 (Essen BioScience). Cell number per well was counted using IncuCyte software (v2022B) and used for normalization of the data. Mitochondrional and glycolytic parameters were calculated using Wave 2.6.3 software (Agilent). Results are shown from five independent experiments, each having 3-6 technical replicates.

To run Cell Mito Stress test in the iMGLs, 40 000 cells per well were seeded in 200 μl maturation medium one week before the experiment. Every other day half of the medium was replaced with fresh medium. On the day of the experiment Seahorse XF Assay medium was prepared by adding 2 mM Glutamax (#35050-038, Invitrogen) to Seahorse DMEM medium. First, the cells were rinsed with 180 μl of Seahorse XF medium following Seahorse XF medium addition to the cells to the final volume of 180 μl. The cells were incubated in a non-CO_2_ incubator for 1h at +37 °C before running on XFe96 Analyzer. At the beginning of the assay, the XFe96 Analyzer added 10mM glucose and 1mM sodium pyruvate (#11360039, Gibco) to the cells. Next, modulators of the electron transport chain were injected at 1 μM concentration to the cells in a following order: oligomycin (#11342, Cayman Chemical), carbonyl cyanide-4 (trifluoromethoxy) phenylhydrazone (FCCP) (#15218, Cayman Chemical) and a mixture of rotenone (#13995, Cayman Chemical) and antimycin A (#A8674, Sigma-Aldrich). During the assay, oxygen consumption rate (OCR) was directly measured by the XFe96 Analyzer. The results were normalized by the cell confluence measured by IncuCyte S3 (Sartorius) before the beginning of the assay. Results are shown from three independent Seahorse XF96 Cell Culture plates, each having 3-4 technical replicates per line.

### Tumor necrosis factor alpha (TNF-α) and interleukin 6 (IL-6) ELISA, LDH-assay

For analyzing acute inflammatory response in mouse primary microglia, 100 000 cells were plated on 48-well plate pre-coated with 0.2 % PLL in culture medium with 5 % FBS. On the following day, the cells were treated with 5 µg/ml LPS or left untreated. After 3 h, 24 h and 48 h, conditioned medium was collected and centrifuged at 10 000 x g, for 10 min at 4 °C. TNFα and IL-6 levels in the conditioned medium were measured using TNF alpha Mouse Uncoated ELISA kit (#88-7324-88, Thermo Fisher) and IL-6 Mouse Uncoated ELISA kit (#88-7064-88, Thermo Fisher).

Cytotoxicity based on the release of lactate dehydrogenase (LDH) by damaged cells was assayed from conditioned medium using Cytotoxicity Detection Kit (#11644793001, Roche). Detection of LDH release was done according to the instructions from manufacturer. Results are shown as a percentage of the untreated WT group. TNF-α, IL-6, and LDH data are from two independent experiments each having three to four technical replicates.

### Phagocytosis assay

For the phagocytosis assay, 20 000 PLCγ2-P522R KI, WT, and TREM2 KO cells well were seeded one day before the experiments to PLL-coated 96-well plates. On the day of the experiment, pHrodo-red-labeled Zymosan (10 µg/ml, #P35364, Invitrogen) and pHrodo-labeled myelin (48 µg/ml) either without or in combination with Cytochalasin D (40 µM, #C2618, Sigma-Aldrich), which inhibits phagocytosis, was added to the wells. Fluorescent images (4 images per well, one image per hour) were taken with IncuCyte S3 Live Cell imaging system using 20X objective for 4 hours. One µM Vybrant DyeCycle Green stain (#V35004, Life Technologies) was then added for 30 minutes to detect nuclei and count cells. All images were analyzed using IncuCyte analysis software. pHrodo integrated density per well was normalized to the cell count within the same well and acquired intensity values are presented as % of pHrodo-Zymosan or pHrodo-myelin-treated WT group at the last time point. Data from four independent experiments each having three to four technical replicates per experiment are shown.

### Measurement of ROS production

To conduct an assay determining intracellular production of ROS, 20 000 PLCγ2-P522R KI (n = 4), WT (n = 3−4), and TREM2 KO (n = 4) cells were seeded on PLL-coated 96-well plate one day before the assay. On the day of the experiment, cells were treated with LPS (1 µg/ml), myelin (30 µg/ml), or menadione (50 µM). CellROX Deep Red reagent (5 µM, #C10422, Invitrogen), which reacts with ROS, was then added to all wells. Fluorescent images (4 images per well) were taken with IncuCyte S3 Live Cell imaging system using 20X objective every four hours for total of 24 hours. After this, 1 µM Vybrant DyeCycle Green stain (#V35004, Life Technologies) was added for 30 minutes to stain nuclei for counting the cells. All images were analyzed using IncuCyte analysis software. CellROX area per well was normalized to the cell count in the same well. Data are shown for 3-4 technical replicates and from two independent experiments. All groups within each experiment were normalized to the untreated WT group at the 24 h time point.

### RNA extraction

Mouse temporo-occipital cortex was homogenized in TRI Reagent (#T9424, Sigma-Aldrich) after which chloroform was added (one fifth of the volume of TRI Reagent) and the tubes were shaken vigorously. Mixtures were incubated at room temperature for 5 min and the phase separation was performed by centrifugation at 12000 × g for 20 min at +4 °C. RNA was precipitated from the aqueous phase with 2-propanol by mixing and incubating at room temperature for 30 min. Samples were centrifuged at 12000 × g for 25 min at +4 °C and the pellets were washed twice with 75 % ethanol. RNA pellets were air-dried and dissolved in RNAse-free H_2_O.

Isolated mouse CD11b+ microglia were resuspended in DPBS and immediately mixed with lysis buffer of the High Pure RNA Isolation kit (#11828665001, Roche). RNA was extracted following the kit instructions and sent for sequencing to Novogene Europe (below).

MDMi cells were washed once with 1 x DPBS, and cells were collected to TRI Reagent where one fifth of the TRI Reagent volume of chloroform was added. Samples were mixed and incubated at room temperature for 2 min followed by centrifugation at 12000 x g for 15 min at +4 °C. The aqueous phase was collected, and room-temperature 2-propanol was added. Samples were vortexed and incubated at room temperature for 10 min. RNA was washed twice with 75 % ethanol and centrifuged 13000 x g for 5 min at +4 °C. Supernatant was removed and the rest of the ethanol was let to evaporate at room temperature for approximately 10 min. RNA pellet was resuspended in RNase free H_2_O and sent for sequencing to Novogene Europe (below).

### RNA sequencing and data analysis

Library preparation and RNA sequencing were conducted by Novogene (Cambridge, UK) and executed with an Illumina high-throughput sequencing platform. In brief, mRNA enrichment was performed with oligo(dT) bead pulldown, from where pulldown material was subjected to fragmentation, followed by reverse transcription, second strand synthesis, A-tailing, and sequencing adaptor ligation. The final amplified and size selected library comprised of 250–300 bp insert cDNA and paired-end 150 bp. Sequencing yielded reads for 20.2–31.34, 29.4–49.9, and 19.3–31.8 million library fragments per CD11b+ mouse microglia, mouse cortex, and human MDMi cells, respectively.

The 150-nucleotide pair-end RNA-seq reads were quality-controlled using FastQC (version 0.11.7) (https://www.bioinformatics.babraham.ac.uk/projects/fastqc/). Reads were then trimmed with Trimmomatic (version 0.39) (Bolger et al., 2014) to remove Illumina sequencing adapters and poor quality read ends, using as essential settings: ILLUMINACLIP:2:30:10:2:true, SLIDINGWINDOW:4:10, LEADING:3, TRAILING:3, MINLEN:50. Trimmed reads were aligned to the Gencode mouse transcriptome version M25 (for genome version GRCm38), or human transcriptome version 38 (for genome version GRCh38) using STAR (version 2.7.9a) (Dobin et al., 2013) with essential non-default settings: –-seedSearchStartLmax 12, –-alignSJoverhangMin 15, –-outFilterMultimapNmax 100, –-outFilterMismatchNmax 33, –-outFilterMatchNminOverLread 0, –-outFilterScoreMinOverLread 0.3, and –-outFilterType BySJout. The unstranded, uniquely mapping, gene-wise counts for primary alignments produced by STAR were collected in R (version 4.2.2 or 4.3.2) using Rsubread::featureCounts (version 2.12.0, 2.12.3 or 2.16.1)(Liao et al., 2019), ranging from 16.2 to 19.7, 25.1 to 43.1, or 14.5 to 25.1 million per CD11b+ mouse microglia, mouse cortex, and human MDMi cells, respectively. Differentially expressed genes (DEGs) between experimental groups were identified in R (version 4.3.2) using DESeq2 (version 1.38.3) (Love et al., 2014) by employing Wald statistic and lfcShrink for FC shrinkage (type=“apeglm”) (Zhu et al., 2019). For the human MDMi samples, the comparisons between PLCγ2-P522R variant carriers (GC) and matched controls (CC) were performed adjusting for the sample delivery batch. Pathway enrichment analysis was performed on the gene lists ranked by the pairwise DEG test log2FCs in R (version 4.2.3) using clusterProfiler::GSEA (version 4.6.2) (Subramanian et al., 2005) with Molecular Signatures Database gene sets (MSigDB, version 2022.1 for mouse CD11b+ microglia, and version 2023.1 for mouse cortex and human MDMi)(Liberzon et al., 2015).

### Proteomic analysis

Sample preparation for proteomics analyses of acutely isolated CD11b+ microglia from PLCγ2-P522R mice was carried out as we have described previously (Martiskainen et al., 2024, preprint). An amount of 350 ng of peptides were separated on a on an in-house packed C18 analytical column (15 cm × 75 µm ID, ReproSil-Pur 120 C18-AQ, 1.9 µm, Dr. Maisch GmbH) using a 120 min long binary gradient of water and acetonitrile (B) containing 0.1% formic acid at flow rate of 300 nL/min and a column temperature of 50°C for CD11b+ microglia. A Data Independent Acquisition Parallel Accumulation–Serial Fragmentation (DIA-PASEF) method with a cycle time of 1.9 s was used for spectrum acquisition. Briefly, ion accumulation and separation using Trapped Ion Mobility Spectrometry (TIMS) was set to a ramp time of 100 ms. One scan cycle included one TIMS full MS scan followed by 17 DIA PASEF scans with two 26 m/z windows covering the m/z range from 350-1200 m/z.

The raw data were analyzed using the software DIA-NN version 1.8 (Demichev et al., 2020) for protein label-free quantification (LFQ). A one protein per gene canonical fasta database of Mus Musculus (download date: January 12th 2023, 21957 entries) from UniProt and a fasta database with 125 potential human contaminations from Maxquant (Cox et al., 2014). Trypsin was defined as protease. DIA-NN was used to generate a spectral library based on the individual data sets and to perform label-free quantification. Two missed cleavages were allowed, and peptide charge states were set to 2-4. Carbamidomethylation of cysteine was defined as static modification. Acetylation of the protein N-term as well as oxidation of methionine were set as variable modifications. The false discovery rate for both peptides and proteins was adjusted to less than 1 %. Data normalization was disabled.

Identification of differentially expressed proteins between genotypes, non-normalized intensities as the starting point, and the subsequent pathway enrichment were done as for the corresponding RNA-seq data, except using MSigDB version 2023.1 (Liberzon et al., 2015).

### Statistical analyses

GraphPad Prism 10.1.2 software was used for all statistical analyses and for the visualization of the data. Shapiro-Wilk’s test was used to test whether the data had normal distribution. Accordingly, the statistical significance was tested using unpaired samples T-test or Mann Whitney test. In case of multiple dependent variables, Two-way ANOVA together with Šídák’s or Tukey’s multiple comparisons test was used. Correlation analyses were done using Pearson correlation coefficients. P-values smaller than 0.05 were considered statistically significant. All data are presented as mean ± SEM.

## RESULTS

### PLCγ2-P522R variant decreases β-amyloid burden in APP/PS1 mice

PLCγ2-P522R variant protects against AD and it was previously shown to reduce β-amyloid pathology in 5XFAD mice (Sims et al., 2017; Tsai et al., 2023). To study if the PLCγ2-P522R variant affects brain β-amyloid pathology in APP/PS1 mice, we stained β-amyloid plaques using a fluorescent amyloid stain, X-34 (Styren, et al., 2000). Image analysis revealed that β-amyloid plaque number (normalized to the size of the analyzed area, p=0.005) and β-amyloid plaque coverage (total plaque area % of the size of the analyzed area, p=0.006) were both significantly decreased in the entorhinal cortex of the 13-month-old APP/PS1 *x* PLCγ2-P522R KI (hereafter shortly A+/P^ki/ki^) mice as compared to the APP/PS1 (hereafter shortly A+/P^wt/wt^) mice (**Figure 1A**). However, a similar reduction was not detected in the hippocampus of the same mice (**Figure S2A**). The average size of individual X-34+ β-amyloid plaques did not differ between the genotypes, neither in the entorhinal cortex (**Figure 1A**) nor hippocampus (**Figure S2A**).

**Figure 1.**
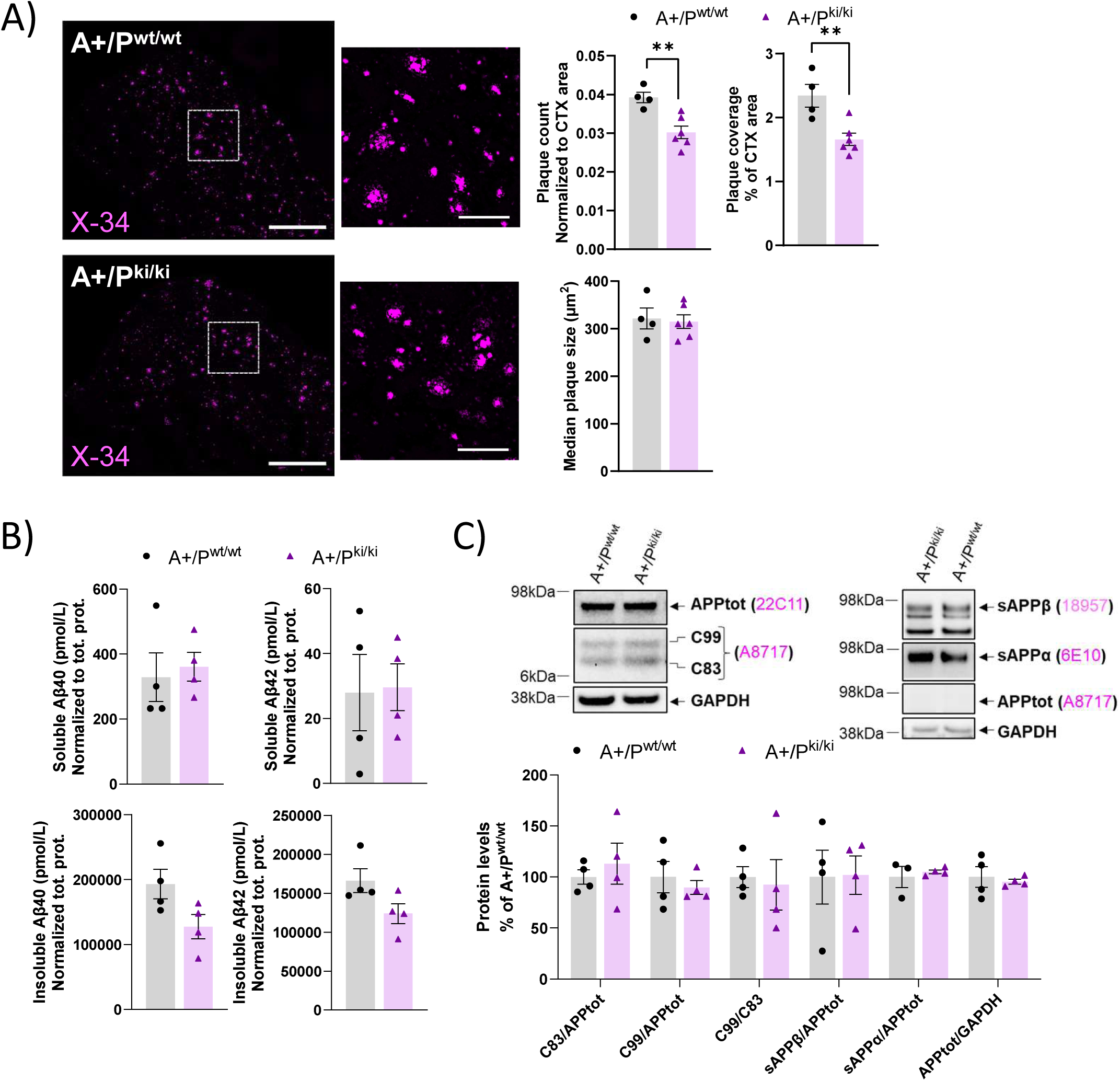
PLCγ2-P522R variant decreases β-amyloid burden in the APP/PS1 mouse brain. A) X-34-positive β-amyloid plaque count (**p=0.005) and coverage (**p=0.006, total plaque area, % of whole analyzed area) are decreased in the entorhinal cortex of the APP/PS1xPLCγ2-P522R (A+/P^ki/ki^) mice as compared to the APP/PS1 (A+/P^wt/wt^) mice. Size (µm^2^) of the individual plaques does not differ between the genotypes. n(A+/P^wt/wt^)=4, n(A+/P^ki/ki^)=6. B) Insoluble, but not soluble Aβ40 and –42 levels are slightly, but not significantly lower in the temporo-occipital cortex of the A+/P^ki/ki^ mice as compared to the A+/P^wt/wt^ mice. Aβ40 and –42 levels are normalized to the total protein concentration in the same sample. n(A+/P^wt/wt^)=4, n(A+/P^ki/ki^)=4. C) Representative Western blots and corresponding quantification show no differences in the levels of full-length APP (APPtot, normalized to GAPDH), APP C-terminal fragments (C99 and C83, normalized to APPtot), or soluble APPα and –β (sAPPα, sAPPβ, normalized to APPtot) species in the temporo-occipital cortex of the A+/P^ki/ki^ and the A+/P^wt/wt^ mice. n(A+/P^wt/wt^)=4, n(A+/P^ki/ki^)=4. Scale bars in the representative immunofluorescent images are 657µm for the whole area and 164µm for the zoomed view. Unpaired samples T-test. All data are presented as mean ± SEM. Each datapoint represents an individual mouse.

In line with this, insoluble, but not the soluble levels of Aβ40 and Aβ42 in the ELISA assay were lower the temporo-occipital cortex of A+/P^ki/ki^ female mice than in A+/P^wt/wt^ mice (**Figure 1B**). A similar trend was observed in the hippocampus, with A+/P^ki/ki^ mice showing a slight, but nonsignificant decrease in insoluble Aβ40 and Aβ 42 levels as compared to the A+/P^wt/wt^ mice (**Figure S2B**). Importantly, there were no differences in the levels of full-length APP (APPtot), APP C-terminal fragments (C83 and C99, **Figure 1C and S2C,** left panel), or soluble APPα and –β (sAPPα and –β, **Figure 1C and S2C** right panel) in the temporo-occipital cortex or hippocampus in Western blot analyses, suggesting that the histochemically determined decrease in the β-amyloid burden in the entorhinal cortex of A+/P^ki/ki^ mice is not due to alterations in APP stability or processing.

### PLCγ2-P522R variant increases microglia clustering around β-amyloid plaques

*PLCG2* expression in the brain is enriched in the microglia (Magno et al., 2019). We have previously shown that the protective PLCγ2-P522R variant increases microglia activation in WT mice without amyloid pathology (Takalo et al., 2020). We now analyzed whether the PLCγ2-P522R variant promotes microglia activation in APP/PS1 mice using PET imaging with TSPO-ligand, [18F]FEPPA, that is highly expressed in activated microglia (Wilson et al., 2008). To increase statistical power, female and male mice were pooled for the analysis. Given the baseline difference between the sexes, the data were normalized to the average of A+/P^wt/wt^ within each sex. There was no difference in the [18F]FEPPA ligand uptake between the genotypes in 7-month-old mice (**Figure 2A**). However, in 13-month-old animals, increased [18F]FEPPA ligand uptake was observed in the whole brain (p=0.032) of A+/P^ki/ki^ mice as compared to the A+/P^wt/wt^ mice. When examining the brain regions separately, [18F]FEPPA ligand uptake was significantly increased in pons (p=0.013) and hippocampus (p=0.012) (**Figure 2B**). This suggests that the protective variant moderately increases microglia activation in APP/PS1 mouse brain.

**Figure 2.**
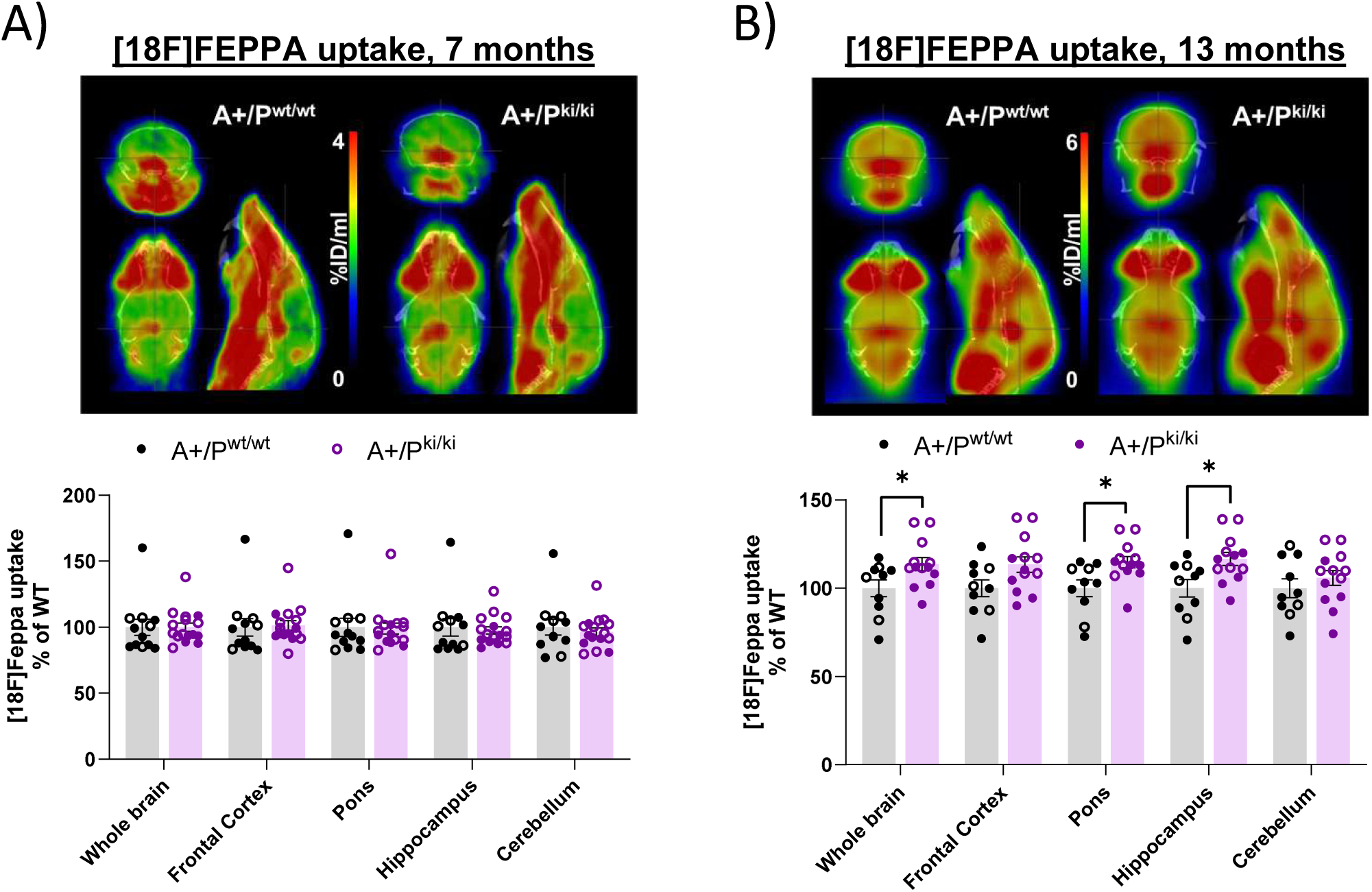
PLCγ2-P522R variant increases microglia activation in the brain of 13-month-old APP/PS1 mice. Example positron emission (PET) images of the uptake (percent of the injected dose per ml of tissue) of [18F]FEPPA for both APP/PS1 (A+/P^wt/wt^) and the APP/PS1xPLCγ2-P522R (A+/P^ki/ki^) mice at (A) 7 months and (B) 13 months of age with an anatomical CT reference image in the background. [18F]FEPPA uptake remains unaltered between the genotypes at the 7-month timepoint. n(A+/P^wt/wt^)=12, n(A+/P^ki/ki^)=17. [18F]FEPPA uptake is significantly increased in the whole brain (*p=0.032), pons (*p=0.013), and hippocampus (*p=0.012) of the 13-month old A+/P^ki/ki^ as compared to A+/P^wt/wt^ mice. n(A+/P^wt/wt^)=10, n(A+/P^ki/ki^)=13. Colored circles indicate data obtained from female mice and hollow circles data obtained from male mice. Data have been normalized to the A+/P^wt/wt^ mice within each sex.

To assess whether the PLCγ2-P522R affects the microglial response to β-amyloid deposition, we quantified the IBA1+ area overlapping and within 0-40 µm distances from the X-34-positive β-amyloid plaque in 13-month-old APP/PS1 mouse brain (**Figure S1A**). IBA1+ area was increased within the 0-40 µm (p=0.03) radius from the β-amyloid plaque outline in the entorhinal cortex of A+/P^ki/ki^ mice as compared to the A+/P^wt/wt^ mice (**Figure 3A**. A similar increase in the IBA1+ area within the 0-40 µm (p=0.037) radius from the β-amyloid plaque outline was observed in the hippocampus of A+/P^ki/ki^ mice as compared to the A+/P^wt/wt^ mice (**Figure S3A**). In contrast, no difference was found in the IBA1+ signal overlapping the β-amyloid plaque area itself, either in the entorhinal cortex or hippocampus. Total IBA1+ area in the entorhinal cortex was marginally increased, while in the hippocampus, total IBA1+ area was decreased (p=0.043). Together, these data suggest that the PLCγ2-P522R variant improves microglia clustering around β-amyloid plaques and therefore may resist β-amyloid deposition more efficiently.

**Figure 3.**
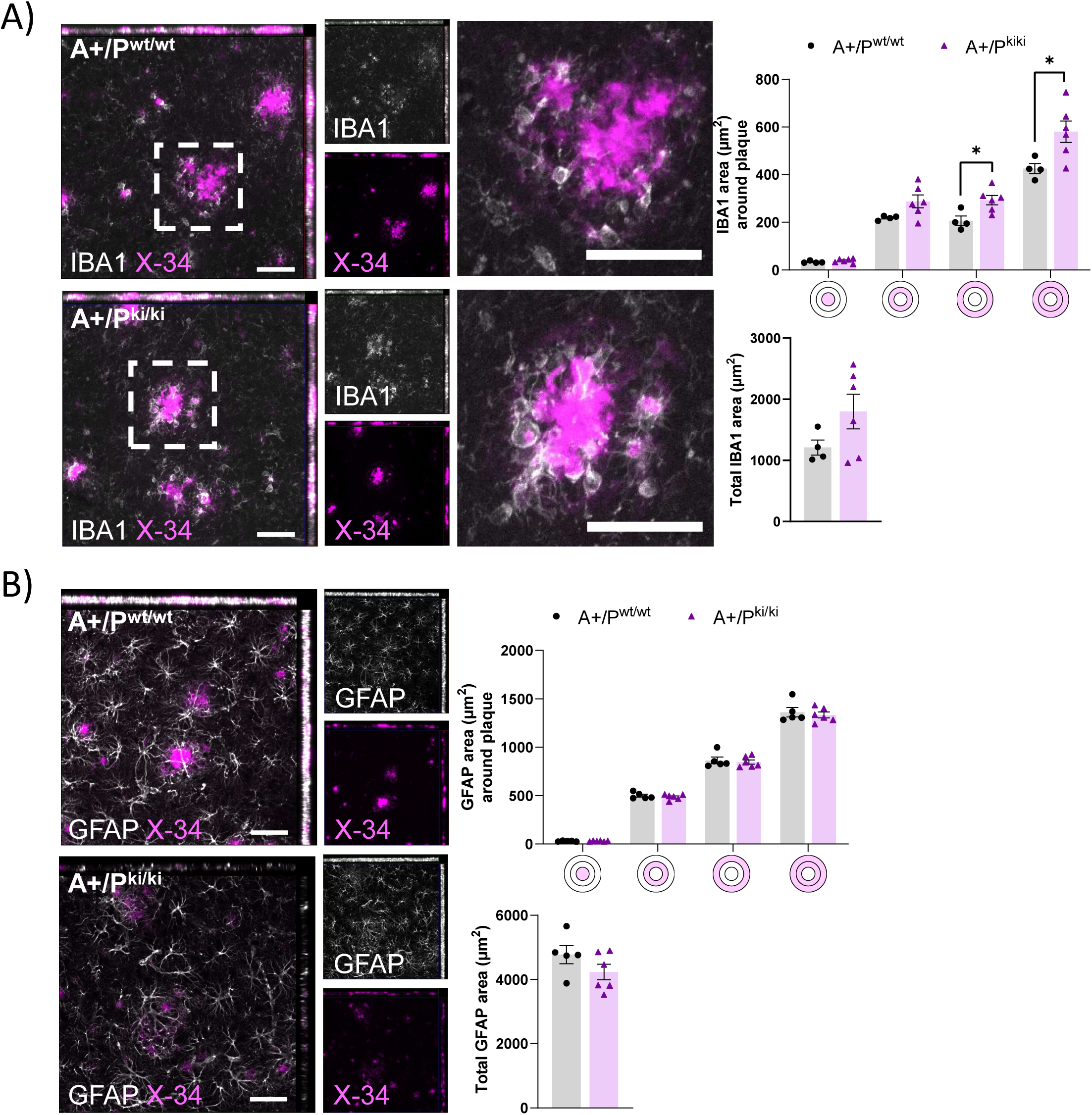
PLCγ2-P522R variant increases microglia clustering around β-amyloid plaques in APP/PS1 mouse brain. A) Area (µm^2^) of IBA1-positive microglia is increased at 20-40 µm (*p=0.017) and 0-40 µm (*p=0.03) distances from the plaque outline in the entorhinal cortex of the APP/PS1xPLCγ2-P522R (A+/P^ki/ki^) mice as compared to the APP/PS1 (A+/P^wt/wt^) mice. Simultaneously, a trend towards an increase in total IBA1 area is observed. n(A+/P^wt/wt^)=4, n(A+/P^ki/ki^)=6. B) GFAP-positive astrocyte area (µm^2^) around plaques (within 0-20 µm and 20-40 µm from the plaque outline) and total GFAP area remain unaltered between the genotypes. n(A+/P^wt/wt^)=5, n(A+/P^ki/ki^)=6. Colored disks on the x-axis of C and D indicate the analyzed area overlapping (middle circle) and surrounding β-amyloid plaque. The scale bar in the representative immunofluorescent images is 50 μm. Unpaired samples T-Test. All data are presented as mean ± SEM. Each datapoint represents an individual mouse.

Since activated microglia secrete chemokines and cytokines which promote astrocyte activation (Liddelow et al., 2017), we stained astrocytes with glial fibrillary acidic protein (GFAP) antibody to study potential changes in astrocytes in the entorhinal cortex and hippocampus of 13-month-old mice (Liddelow et al., 2017). We found no differences in the total GFAP+ area or in the GFAP+ area overlapping or surrounding (within 0-40 µm radius) X-34-positive β-amyloid plaques (**Figure S1B**) either in the entorhinal cortex (**Figure 3B**) or in the hippocampus (**Figure S3B**) of A+/P^ki/ki^ mice as compared to the A+/P^wt/wt^ mice, suggesting that PLCγ2-P522R variant-driven microglia activation does not promote astrocyte activation in these mice.

### PLCγ2-P522R variant decreases dystrophic neurites around β-amyloid plaques

Microglia clustered around β-amyloid plaques have been suggested to pack diffuse β-amyloid into a more compacted and less toxic form (Yuan et al., 2016). Hence, to investigate β-amyloid plaque morphology, we used X-34 to detect compact plaques and anti-β-amyloid antibody 6E10 to detect diffuse plaques. Despite of the evident improvement in microglia response towards β-amyloid plaques, we found no differences between the genotypes in the total 6E10+ area or in the percentage of X-34+ or 6E10+ area of total β-amyloid burden (X-34+6E10) either in the entorhinal cortex (**Figure 4A**) or in the hippocampus (**Figure S4A**). Furthermore, no differences in the colocalization of 6E10 within IBA1+ area was found between the genotypes (**Figure 4B and S4B**), suggesting that the PLCγ2-P522R variant does not promote microglial uptake of diffuse β-amyloid nor changes the ratio of diffuse and compact β-amyloid in the brain of APP/PS1 mice.

**Figure 4.**
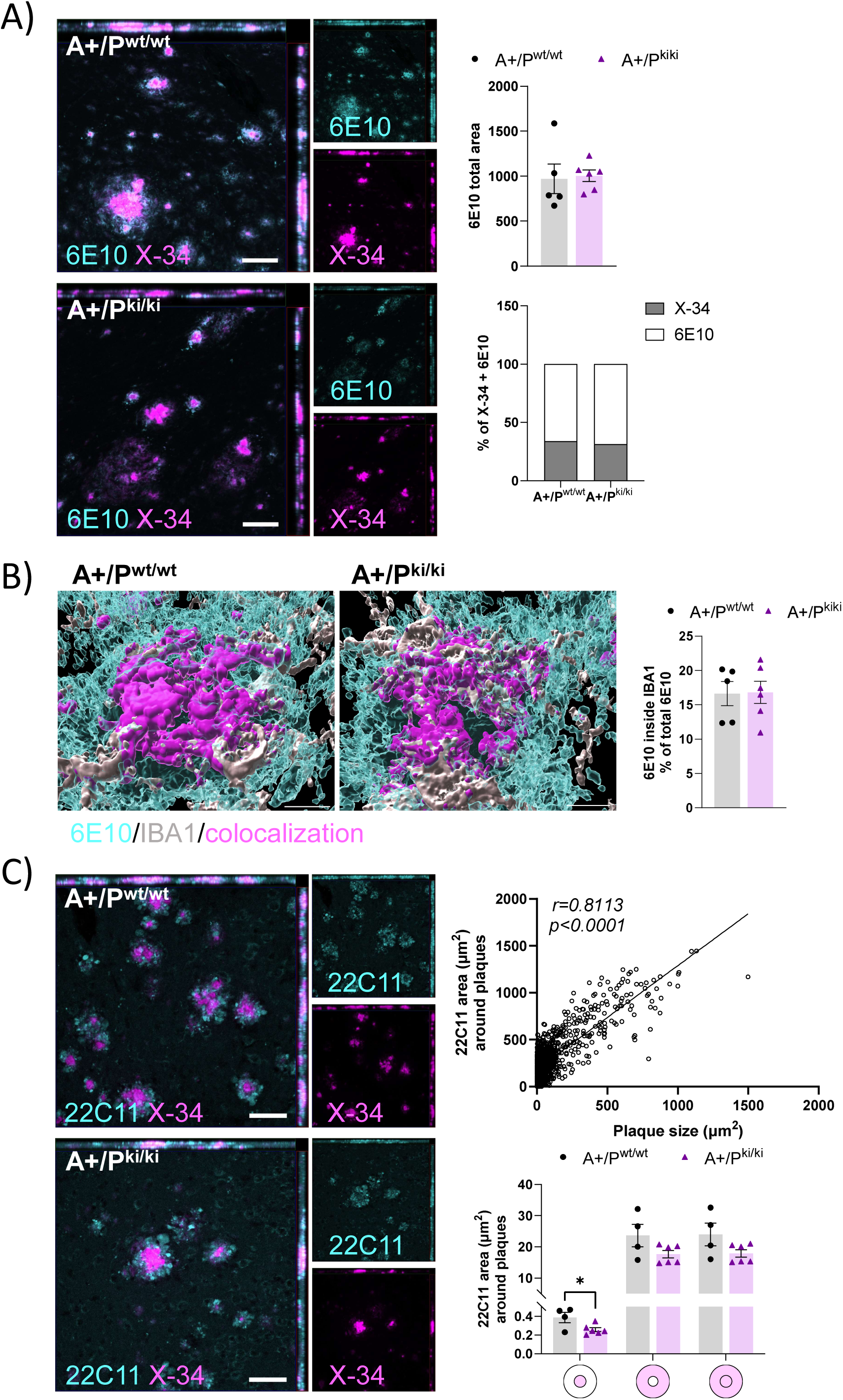
PLCγ2-P522R variant decreases β-amyloid plaque-associated dystrophic neurites in APP/PS1 mouse brain. A) Total area (µm^2^) of diffuse β-amyloid (6E10) and composition β-amyloid plaques, as indicated by the percentage of compact (X-34) and diffuse (6E10) β-amyloid, remain unaltered in the entorhinal cortex of the APP/PS1xPlγ2-P522R (A+/P^ki/ki^) mice as compared to the APP/PS1 (A+/P^wt/wt^) mice. n(A+/P^wt/wt^)=5, n(A+/P^ki/ki^)=6. B) A 3D-reconstruction showing 6E10 and IBA1 (microglia) signal and their co-localization. Respective quantification showing 6E10 within IBA1-positive area as % of all 6E10 signal in the entorhinal cortex of A+/P^ki/ki^ mice as compared to the A+/P^wt/wt^ mice. C) Area (µm^2^) of 22C11-labeled dystrophic neurites around β-amyloid plaques strongly correlates with plaque size (r=0.8113, ****p<0.0001). 22C11-positive area shows a decrease within β-amyloid plaque area (p=0.033) and a similar decreasing trend within the 0-14 µm radius from the β-amyloid plaque outline in the entorhinal cortex of the A+/P^ki/ki^ mice as compared to the A+/P^wt/wt^ mice when normalized to the plaque size. n(A+/P^wt/wt^)=4, n(A+/P^ki/ki^)=6. Colored disks on the x-axis indicate the analyzed area overlapping (middle circle) and surrounding β-amyloid plaque. Scale bars in the representative immunofluorescent images are 50 μm. Pearson correlation and unpaired samples T-test. All data are presented as mean ± SEM. Each datapoint represents an individual mouse.

To assess whether the PLCγ2-P522R variant influences the formation of neuronal dystrophies in the APP/PS1 mouse brain, brain sections were stained with the N-terminal APP antibody, 22C11 (Sadleir et al., 2016). Since the 22C11 signal was strongly dependent on the β-amyloid plaque size (Pearson correlation r=0.81, p<0.0001) (**Figure 4C**), we normalized the 22C11+ area around each plaque to the size of the analyzed plaque. The 22C11+ area overlapping the plaque area in the entorhinal cortex was significantly smaller (p=0.033) in A+P^ki/ki^ mice than A+/P^wt/wt^ mice, and a similar but nonsignificant trend was observed within the 0-14 µm radius from the plaque outline (**Figure S1C and 4C**). In the hippocampus, 22C11+ area was significantly decreased within the 0-14 µm radius from the plaque outline (p=0.010) (**Figure S4C**). Taken together, the PLCγ2-P522R variant decreases the formation of dystrophic neurites around β-amyloid plaques, suggesting that enhanced microglial barrier function promoted by the protective variant alleviates neurotoxic effects caused by β-amyloid deposits.

### PLCγ2-P522R variant alleviates inflammation in the brain of APP/PS1 mice

The well-established functions of microglia to compact β-amyloid deposits and protect surrounding neurons are considered beneficial. However, microglia activation also promotes neuroinflammation, which in the chronic state is harmful to neuronal health. Therefore, we next analyzed the levels of several inflammatory markers in 13-month-old mouse temporo-occipital cortex and hippocampal lysates using MSD mouse inflammatory panel. In general, A+/P^ki/ki^ mice appeared to have lower levels of several inflammatory markers in both temporo-occipital cortex and hippocampus than the A+/P^wt/wt^ mice (**Figure 5**). The levels of IL-12-p70 (p70 refers to an active heterodimer, p=0.049) and IL-6 (p=0.003) in A+/P^ki/ki^ mice were significantly lower in the cortex, while those of IL-1β (p=0.021) and IL-8 (p=0.026) were lower in the hippocampus. This suggests that despite increasing microglia activation, PLCγ2-P522R variant does not increase chronic β-amyloid-associated neuroinflammation in APP/PS1 mouse brain upon aging.

**Figure 5.**
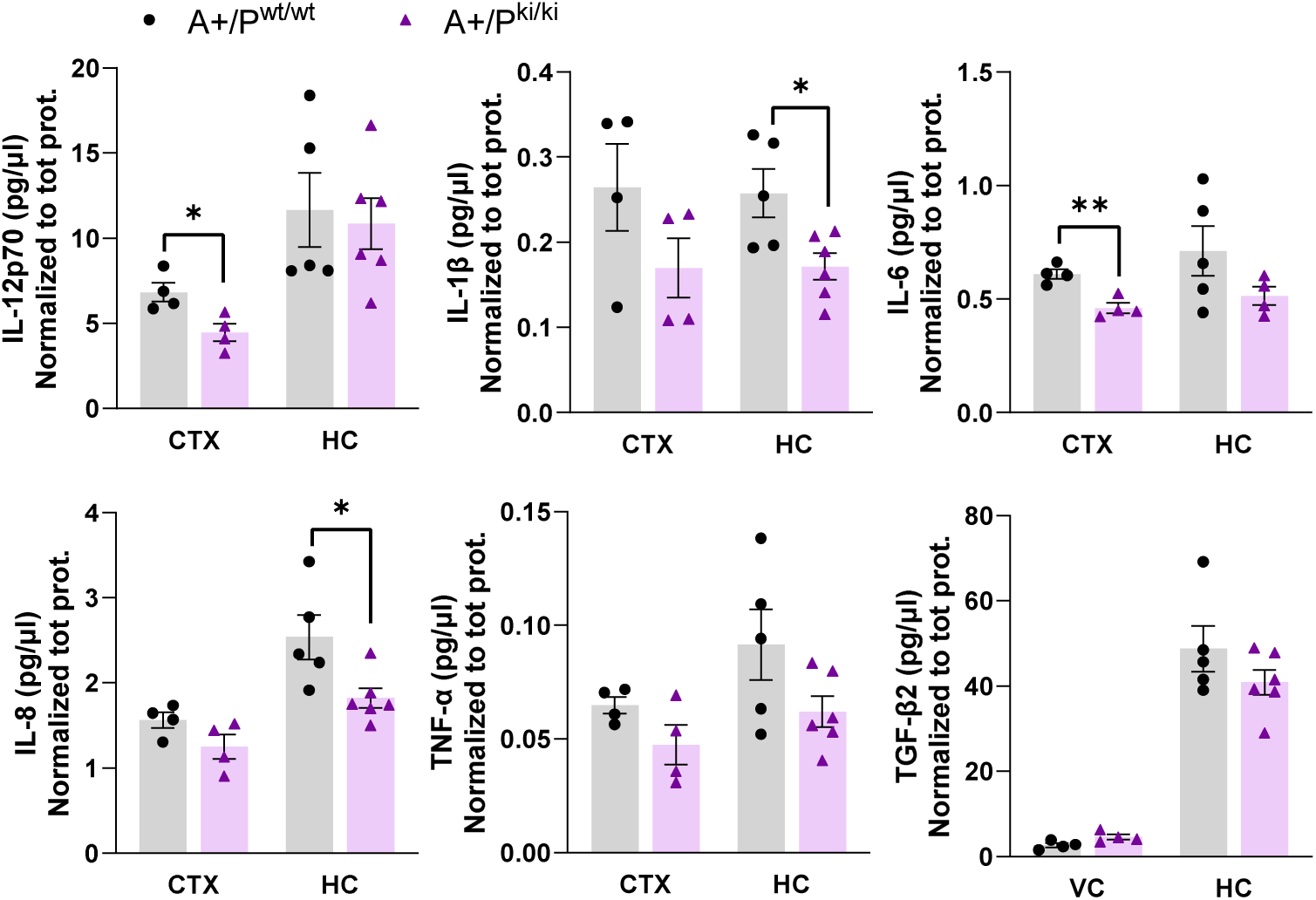
PLCγ2-P522R variant alleviates neuroinflammation in APP/PS1 mouse brain. A) IL-12p70 (p=0.049) and IL-6 (p=0.003) levels are significantly lower in the temporo-occipital cortex (CTX) while IL-1β (p=0.021) and IL-8 (p=0.026) levels are significantly lower in the hippocampus (HC) of the APP/PS1xPlγ2-P522R (A+/P^ki/ki^) mice as compared to the APP/PS1 (A+/P^wt/wt^) mice. In addition, IL-1β, IL-8, and TNF-α levels show a trend towards a decrease in the CTX while IL-6, TNF-α, and TGF-β2 levels show a trend towards a decrease in the HC of the A+/P^ki/ki^ as compared to the A+/P^wt/wt^ mice. CTX(A+/P^wt/wt^) n=4, CTX(A+/P^ki/ki^) n=4, HC(A+/P^wt/wt^) n=5, HC(A+/P^ki/ki^) n=6. Unpaired samples T-test. All data are presented as mean ± SEM. Each datapoint represents an individual mouse.

### PLCγ2-P522R variant does not increase β-amyloid plaque-associated APOE or promote DAM signature

β-amyloid plaque-associated APOE has been suggested to enhance plaque compaction and support microglia responses towards β-amyloid deposition (Parhizkar et al., 2019; Ulrich et al., 2018). To address if APOE plays a role in PLCγ2-P522R variant-mediated microglia response to β-amyloid, we analyzed APOE+ area within and surrounding plaques. We found no differences in the APOE+ area overlapping or within 10 µm radius from the plaque outline (**Figure S1D**) between A+P^ki/ki^ and A+/P^wt/wt^ mice in the entorhinal cortex (**Figure 6A**) or in the hippocampus (**Figure S5A**). In line with this, soluble and insoluble APOE levels in the temporo-occipital cortex and hippocampus did not differ between the genotypes (**Figure S5B**).

**Figure 6.**
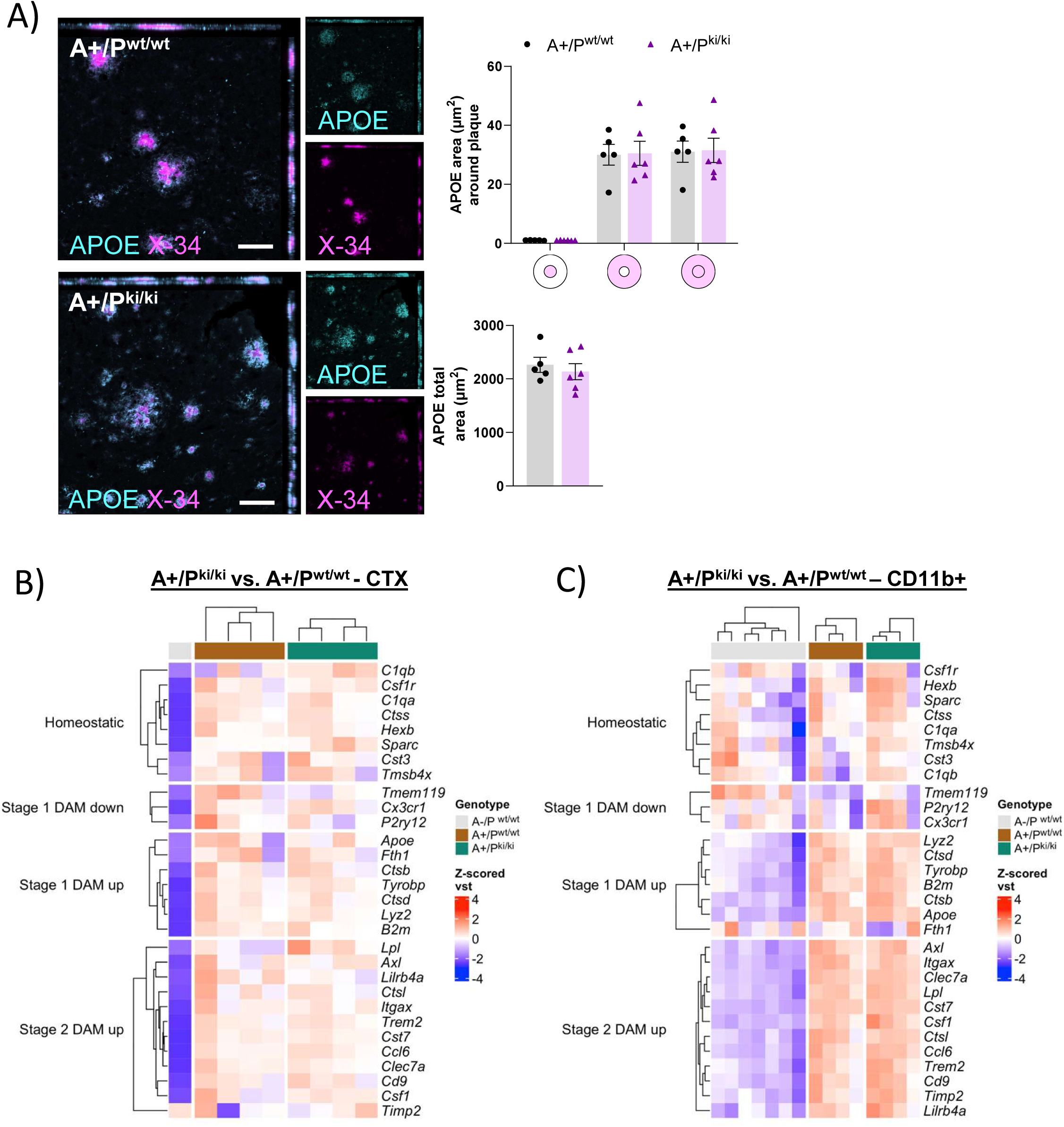
PLCγ2-P522R variant does not increase plaque-associated APOE or disease-associated microglia RNA signature in APP/PS1 mouse brain. A) Analysis of total APOE and APOE area (µm^2^) within and surrounding (within 0-10 and 10-20 µm distance from the β-amyloid plaque outline) β-amyloid plaques reveals no differences between the APP/PS1xPlγ2-P522R (A+/P^ki/ki^) and APP/PS1 (A+/P^wt/wt^) mice. n(A+/P^wt/wt^)=5, n(A+/P^ki/ki^)=6. Colored disks on the x-axis indicate the analyzed area overlapping (middle circle) and surrounding β-amyloid plaque. B) Soluble and insoluble APOE levels in the temporo-occipital cortex (CTX) and in the hippocampus (HC) are similar between the genotypes. CTX(A+/P^wt/wt^) n=4, CTX(A+/P^ki/ki^) n=4, HC(A+/P^wt/wt^) n=5, HC(A+/P^ki/ki^) n=6. C) RNA expression of *Apoe* and *Trem2* in the CTX and CD11b+ microglia isolated from the whole brain of A+/P^wt/wt^ and A+/P^ki/ki^ mice remain unaltered. CTX(A+/P^wt/wt^) n=4, CTX(A+/P^ki/ki^) n=4, CD11b+(A+/P^wt/wt^) n=4, CD11b+(A+/P^ki/ki^) n=4. D) Soluble TREM2 (sTREM2) levels do not differ between A+/P^wt/wt^ and A+/P^ki/ki^ mice either in the CTX or HC. CTX(A+/P^wt/wt^) n=4, CTX(A+/P^ki/ki^) n=4, nHC(A+/P^wt/wt^)=5, HC(A+/P^ki/ki^) n=6. The scale bar in the representative immunofluorescent images is 50 μm. Unpaired samples T-Test. All data are presented as mean ± SEM. Each datapoint represents an individual mouse.

Upregulation of *Apoe* is part of the disease-associated microglia (DAM) signature (Keren-Shaul et al., 2017). DAM activation is regulated by *Trem2* expression and it is needed for restricting β-amyloid pathology in AD (Keren-Shaul et al., 2017). Hence, to study if the protective variant exacerbates DAM signature in the brain of APP/PS1 mice, RNA sequencing was carried out in the temporo-occipital cortex and in CD11b+ microglia acutely isolated from the whole brain. The expression of both *Apoe* and *Trem2* in cortical tissue and CD11b+ microglia did not differ between the A+/P^ki/ki^and A+/P^wt/wt^ mice, suggesting that the PLCγ2-P522R variant does not further push microglia towards the DAM signature in APP/PS1 mice (**Figure 6 B and C**). In fact, no DEGs or enriched biological pathways were detected in temporo-occipital cortex (**Figure S5D**) or isolated CD11b+microglia (**Figure S5E**) when A+/P^ki/ki^ were compared to A+/P^wt/wt^ mice.

At the protein level, shedding of soluble TREM2 (sTREM2) terminates signaling of full-length TREM2 and therefore, influences microglia activation (Schlepckow et al., 2020). To assess whether PLCγ2-P522R modulates TREM2 shedding, we measured TREM2 levels in the PBS soluble faction of CTX and HC lysates. There were no differences between the genotypes in the levels of sTREM2 in these brain areas (**Figure S5C**), suggesting that the protective variant does not promote negative feed-back-loop restricting TREM2 signaling.

### PLCγ2-P522R variant promotes anxiety in APP/PS1 mice

Given that PLCγ2-P522R reduces β-amyloid deposition in the brain and associated neuronal pathologies in APP/PS1 mice, we next wanted to investigate if the protective variant influences the behavioral alterations observed in APP/PS1 mice. First, the body weight showed a APP/PS1 x PLCγ2-P522R genotype interaction (F_1,64_ = 4.4, p = 0.04), such that the APP/PS1 wildtype (-) A-/P^ki/ki^ mice had a lower weight than the A-/P^wt/wt^ littermates (p=0.024), but the same was not detected among the APP/PS1 hemizygous (+) mice (**Figure 7A**). Given that the PLCγ2-P522R variant did not influence the ambulatory distance travelled in the spontaneous activity test (**Figure 7B**), the observed weight reduction was not related to increased activity. However, the time spent in the center of the activity-monitoring box was significantly shorter in the P^ki/ki^ than P^wt/wt^ mice regardless of the APP/PS1 genotype (F_2,64_=7.2, p=0.02, **Figure 7C**), suggesting that the PLCγ2-P522R variant increases the avoidance of open places. In the light-dark box, the mice were given a free choice between a lit and a dark compartment. In this test, the ANOVA showed a main effect of both the PLCγ2-P522R (p=0.06) and APP/PS1 (p=0.26) genotypes and their interaction (p=0.035). The P^ki/ki^ mice showed an increased avoidance of the lit environment, which was further accentuated by the APP/PS1+ genotype (p=0.03, **Figure 7D**). In the passive avoidance test, performed a day after the mice had received a foot shock when entering the dark compartment, the ANOVA showed an even stronger main effect of both the PLCγ2-P522R (p<0.001) and APP/PS1 (p<0.001) genotypes and their interaction (p=0.017). On the day after the foot shock, the P^ki/ki^ mice rushed into the dark compartment, whereas the P^wt/wt^ mice hesitated to enter (p<0.0001, **Figure 7E**). This tendency was further accentuated by the APP/PS1+ genotype. Given the fact that P^ki/ki^ and P^wt/wt^ mice did not differ in their motor abilities in the rotating rod (**Figure 7F**) and that they had similar pain threshold (**Figure 7G**), this outcome was not explained by declined motor capability or impaired ability to sense pain.

**Figure 7.**
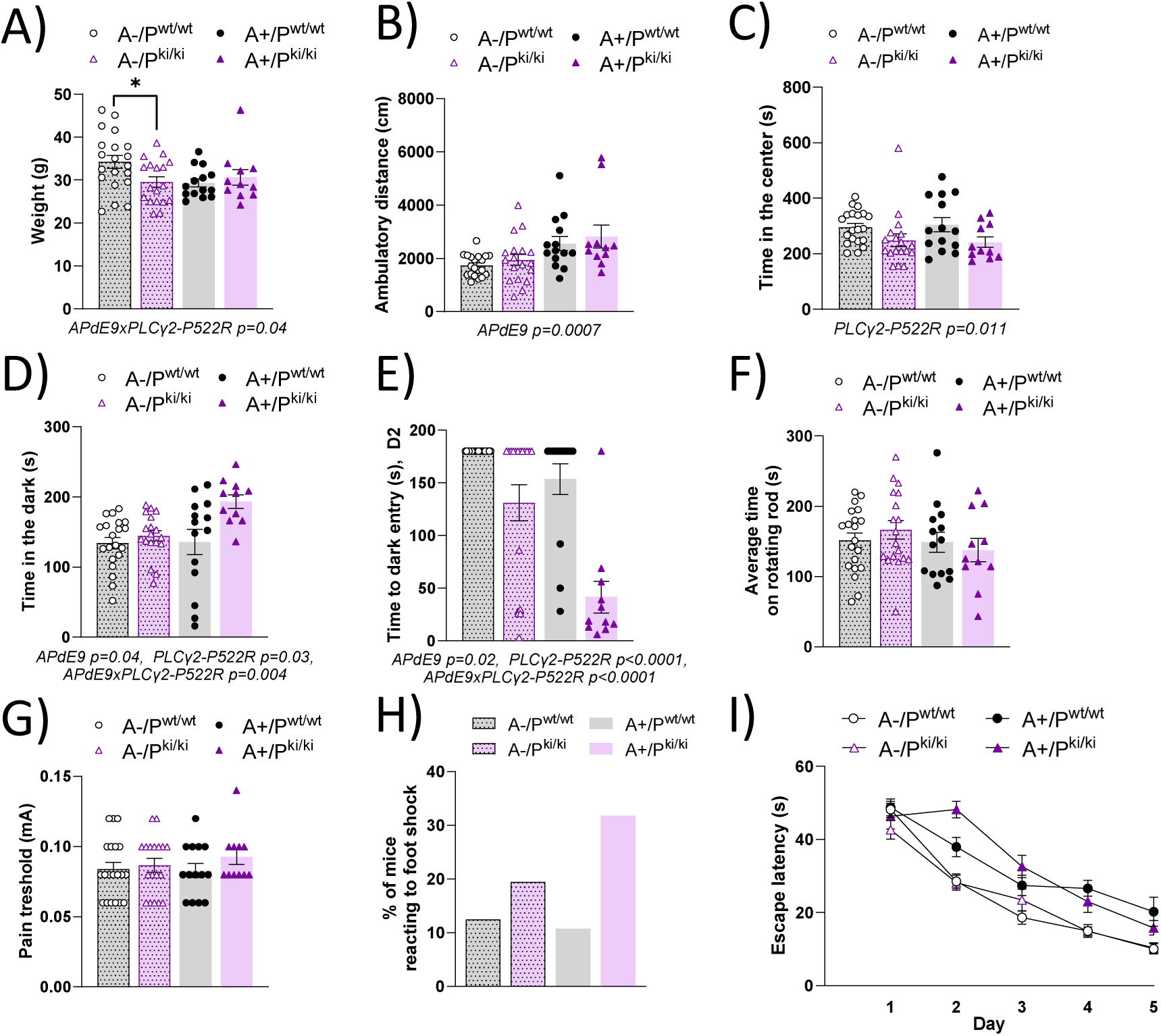
PLCγ2-P522R mice show avoidance response to open places, bright environment, and sudden unpleasant stimulus. A) Body weight of the PLCγ2-P522R (P^ki/ki^) mice is lower than their wildtype (P^wt/wt^) littermates in APP/PS1-negative (A-, *p=0.024) but not in the APP/PS1 hemizygous background (A+). B) PLCγ2-P522R does not influence ambulatory distance travelled in the spontaneous activity test. C) Time spent in the center of the novel test box is significantly shorter in the P^ki/ki^ mice regardless of the APP/PS1 genotype. D) In passive avoidance test, the P^ki/ki^ mice show an increased avoidance of the lit compartment, which is further accentuated by the APP/PS1 hemizygous genotype. E) Even after receiving a foot shock, the P^ki/ki^ mice have shorter latency to enter the dark compartment as compared to the P^wt/wt^ mice. This tendency is further accentuated by the APP/PS1 hemizygous genotype. F) No difference in motor skills is observed between the genotypes as indicated by the time spent in the rotating rod. G) P^ki/ki^ and P^wt/wt^ mice have similar threshold to pain (foot shock), but H) P^ki/ki^ mice have a stronger behavioral response (crab-like walking backwards) to the electric foot shock. I) PLCγ2-P522R does not influence spatial learning and memory in the Morris swim navigation task. APP/PS1 hemizygous genotype strongly impairs learning in terms of escape latency (F_1,64_ = 46.8, p < 0.001). n(A-/P^wt/wt^)=20, n(A-/P^ki/ki^)=18, n(A+/P^wt/wt^)=14, n(A+/P^ki/ki^)=11. Two-way ANOVA with Šídák’s multiple comparisons test. All data are presented as mean ± SEM. Each datapoint represents an individual mouse.

In fact, the P^ki/ki^ mice had a stronger behavioral response (crab-like walking backwards) to the foot shock than the P^wt/wt^ mice (**Figure 7H**). Finally, PLCγ2-P522R variant did not influence spatial learning and memory in the Morris swim navigation task either in mice with APP/PS1-or APP/PS1+ background (**Figure 7I**), whereas APP/PS1+ strongly impaired learning in terms of escape latency (F_1,64_ = 46.8, p < 0.001). This suggest that rushing into the dark is not explained by declined memory related to the foot shock in the P^ki/ki^ mice.

### PLCγ2-P522R variant improves lipid processing in primary microglia

To address cellular mechanisms affected by the PLCγ2-P522R variant, we utilized cultures of PLCγ2-P522R KI and WT mouse primary microglia. Impaired lipid metabolism potentially underlies the dysfunctional phenotype observed in TREM2 KO and PLCγ2 KO microglia (Andreone et al., 2020; Van Lengerich et al., 2023). Thus, analysis of intracellular LDs was performed to study if the protective PLCγ2-P522R has opposite effects to the KO on lipid accumulation upon LPS– and myelin –induced stress conditions. LPS and even more so myelin treatment increased the number of LD-positive microglia as compared to the untreated microglia, whereas the genotype had no effect on the percentage of LD-accumulating microglia (**Figure 8A**). However, the size of the LDs was clearly smaller in the PLCγ2-P522R KI as compared to WT microglia (p<0.0001) upon myelin treatment. A similar, but non-significant trend was also detected in LPS-treated microglia. These findings suggest that PLCγ2-P522R KI reduces the buildup of large LDs in microglia upon cellular stress conditions.

**Figure 8.**
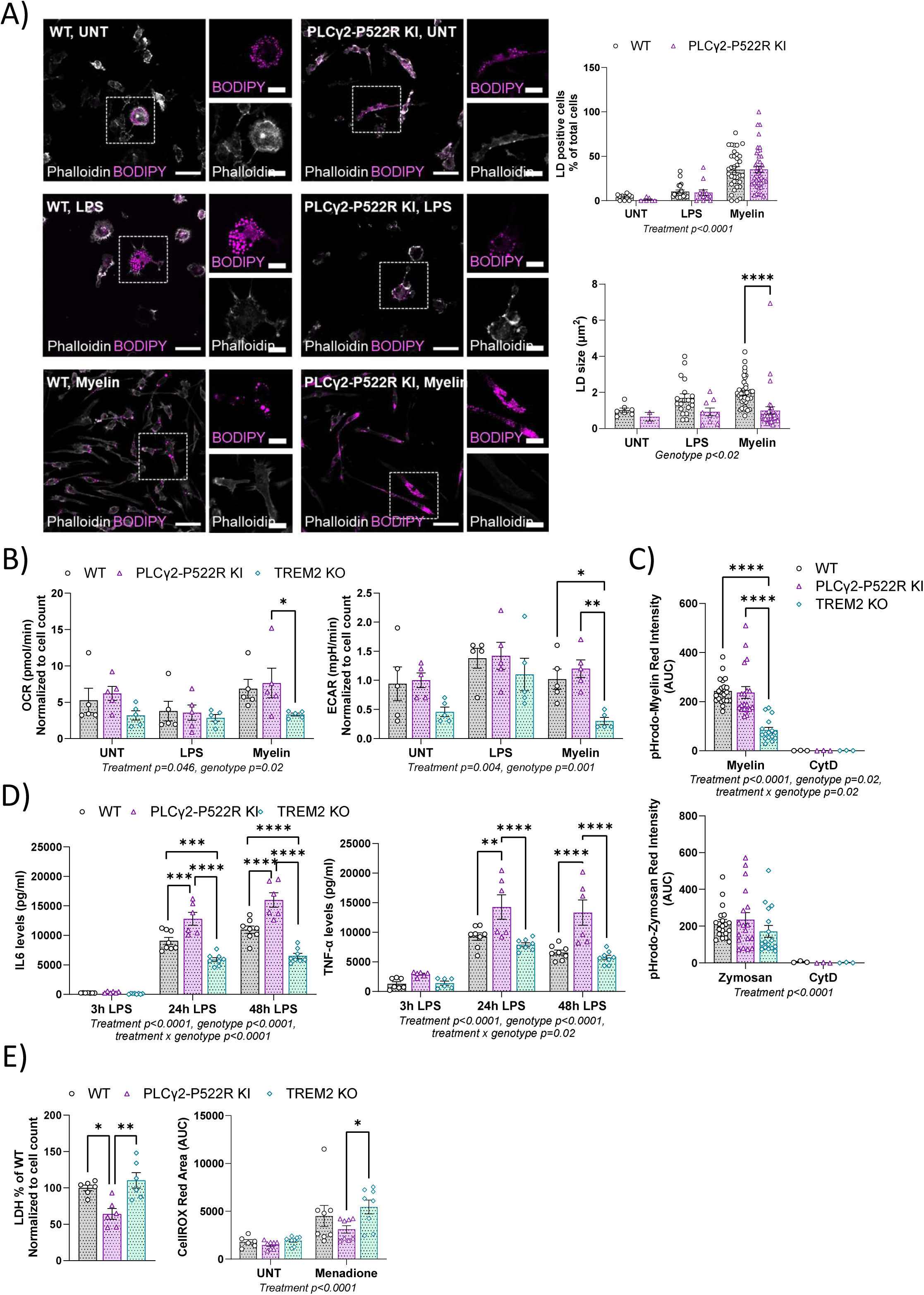
PLCγ2-P522R changes lipid accumulation in mouse primary microglia. A) The percentag of lipid droplet (LD)-positive cells is increased in mouse primary microglia upon 24-h lipopolysaccharide (LPS) and 48-h myelin treatments in comparison to untreated cells (UNT). The percentage of LD-positive cells do not differ between wildtype (WT) and PLCγ2-P522R knock-in (KI) microglia. PLCγ2-P522R KI decreases the size (µm^2^) of individual LDs in myelin-treated conditions (****p<0.0001). n(analyzed images) WT UNT=11, LPS=22, myelin=42; KI PLCγ2-P522R UNT=6, LPS=12, myelin=35. B)) Mitochondrial oxygen consumption rate (OCR, maximal respiration) and glycolytic capacity in mitochondrial and glycolysis stress tests, respectively, are improved in PLCγ2-P522R KI microglia as compared to TREM2 knock-out (KO) microglia under myelin treatment (*p=0.04 and **p=0.005, respectively). Glycolytic capacity also differs between TREM2 KO and WT microglia (*p=0.029). n=5 independent experiments, minimum of 4 technical replicates per experiment. C) Myelin phagocytosis is decreased in TREM2 KO microglia as compared to PLCγ2-P522R KI (****p<0.0001) and WT cells (****p<0.0001). PLCγ2-P522R KI group did not significantly differ from WT group. Zymosan uptake does not differ between the genotypes. Cytochalasin D (CytD) is used as a control. n=5 independent experiments, 4 technical replicates per experiment. D) PLCγ2-P522R increases IL-6 levels in the conditioned medium upon 24-h and 48-h LPS treatment as compared to WT (***p=0.0001 and ****p<0.0001, respectively) and TREM2 KO (****p<0.0001, both) microglia. TREM2 KO cells also secrete less IL-6 than WT cells at both timepoints (***p=0.0004 and ****p<0.00001). TNF-α levels in the medium of PLCγ2-P522R KI microglia are increased upon 24-h and 48-h LPS treatment as compared to WT (**p=0.001 and ****p<0.0001, respectively) and TREM2 KO cells (****p<0.0001, both). n=2 independent experiments, 3-4 technical replicates per experiment. E) Lactate dehydrogenase (LDH) –mediated cytotoxicity is decreased in PLCγ2-P522R KI microglia as compared to WT (*p=0.015) and TREM2 KO (**p=0.002) cells. PLCγ2-P522R KI microglia produce less reactive oxygen species (ROS) as compared to TREM2 KO cells upon menadione-induced oxidative stress (*p=0.02). n=2 independent experiments, 3-4 technical replicates per experiment. Two-way ANOVA with Tukey’s multiple comparisons test. All data are presented as mean ± SEM.

Next, mitochondrial respiration and glycolysis, phagocytosis, inflammatory response, and cellular stress were assessed to study the functional phenotype associated with the PLCγ2-P522R variant in mouse primary microglia. Given that TREM2 depletion impairs mitochondrial function, lipid metabolism, and viability of microglia, (Andreone et al., 2020; Keren-Shaul et al., 2017; Kunkle et al., 2019; Yuan et al., 2016), TREM2 KO primary microglia were included as controls in these assays. To study potential changes in energy production, mitochondrial oxygen consumption rate (OCR) and glycolysis were monitored upon LPS and myelin treatments. As compared to untreated condition, LPS shifted the energy metabolism towards glycolysis in all microglia, whereas myelin induced mitochondrial respiration in WT and PLCγ2-P522R KI microglia (**Figure 8B**). As expected, TREM2 KO microglia showed impaired glycolytic capacity (p=0.029) upon the myelin challenge. They also showed a similar decreasing trend of glycolytic capacity already in the untreated condition when compared to WT microglia. Similar reduction was observed in mitochondrial maximal OCR, but the difference was not statistically significant. TREM2 KO did not influence energy metabolism upon LPS treatment, suggesting that TREM2 is not associated in TLR-mediated cellular responses. As compared to TREM2 KO cells, PLCγ2-P522R KI microglia had significantly better maximal OCR (p=0.04) and glycolytic capacity (p=0.005) in myelin-treated conditions. However, PLCγ2-P522R KI microglia were not significantly different from WT cells (**Figure 8B**).

Increased accumulation of LDs impairs phagocytosis among other immune functions in microglia (Marschallinger et al., 2020). To analyze phagocytic capacity, the uptake of pHrodo-conjugated zymosan and myelin were monitored for 4 h in PLCγ2-P522R KI, TREM2 KO, and WT microglia. TREM2 KO microglia showed a significant impairment in myelin uptake as compared to WT cells (p<0.0001, **Figure 8C**). In contrast, TREM2 KO did not affect zymosan phagocytosis, again suggesting that TREM2 KO is not associated with TLR-mediated cellular functions. Myelin phagocytosis was significantly better in PLCγ2-P522R KI as compared to TREM2 KO microglia (p<0.0001). Neither myelin nor zymosan uptake were different between PLCγ2-P522R KI and WT cells.

We have previously shown that PLCγ2-P522R variant enhances acute inflammatory response in KI mouse macrophages (Takalo et al., 2020). In line with this, we detected significantly higher levels of IL-6 and TNF-α in the conditioned medium of PLCγ2-P522R KI microglia as compared to WT microglia upon 24-h (IL-6 p=0.0001, TNF-α p=0.001) and 48-h (IL-6 p<0.0001, TNF-α p<0.0001) LPS treatment (**Figure 8D**). On the contrary, TREM2 KO microglia showed significantly reduced IL-6 levels in comparison to WT and PLCγ2-P522R KI groups both at the 24-h (WT p=0.0004, PLCγ2-P522R KI p<0.0001) and 48-h (p<0.0001 both WT and PLCγ2-P522R KI) timepoints. TNF-α levels in TREM2 KO microglia medium were significantly lower than in PLCγ2-P522R group (p<0.0001 both 24 h and 48 h) but did not significantly differ from the WT group. These data show that PLCγ2-P522R KI and TREM2 KO microglia have opposite effects on acute inflammatory response.

As an indicator of cell stress, LDH-mediated cytotoxicity and production of ROS were analyzed (**Figure 8E**). LDH-mediated cytotoxicity was significantly lower in PLCγ2-P522R KI microglia as compared to WT (p=0.015) and TREM2 KO (p=0.002) microglia. In addition, under menadione-induced oxidative stress, the production of ROS was lower in PLCγ2-P522R KI as compared TREM2 KO microglia (p=0.02) and a similar, non-significant trend was observed between PLCγ2-P522R KI and WT microglia. Altogether, our findings suggest that PLCγ2-P522R decreases accumulation of large LDs, which associates with an enhanced acute inflammatory response and higher tolerance against cellular stress.

### PLCγ2-P522R KI variant upregulates calcium signaling, fatty acid metabolism, and oxidative phosphorylation-related biological pathways in the acutely isolated adult mouse microglia

To elucidate molecular pathways associated with the PLCγ2-P522R variant, global RNA sequencing and proteomics analyses were performed in CD11b+ microglia acutely isolated from 13-month-old PLCγ2-P522R KI and WT mice without the AD background. RNA sequencing revealed dramatically different RNA signatures between PLCγ2-P522R KI and WT microglia, including 3719 DEGs (FDR 0.05) (**Figure S6A**). Out of these, 1302 were downregulated and 2417 were upregulated. Gene enrichment analysis indicated several down– and upregulated biological pathways in PLCγ2-P522R KI as compared to WT microglia (**Figure 9A**). Among these, calcium signaling, FA metabolism, cholesterol homeostasis, and oxidative phosphorylation showed prominent upregulation in PLCγ2-P522R KI microglia, whereas pathways related to inflammatory response and interferon-α (IFN-α) response were downregulated. Closer examination revealed that core enrichment genes within calcium signaling pathway included *Itpr1, Itpka, Camk4,* Camk2a and *Camk2b* among many others, whereas calcium-sensitive MEF2 transcription family members *Mef2a, Mef2c,* and *Mef2d,* were downregulated in PLCγ2-P522R KI microglia (**Figure 9B, S6C**). Upregulated genes within FA metabolism pathway included e.g., genes linked to LD metabolism and FA synthesis (*Mgll, Fasn, Fads2, Fabp7, Fabp5, Fabp3,* **Figure 9B, S6C**). Interestingly, several genes encoding PI3K subunits, *Pik3cd, Pik3cg, Pik3r5, Pik3ap1, Pik3r1,* and *Pik3r4* were downregulated (**Figure S6C**). Furthermore, several mitochondrial complex I (*Ndufa1, Ndufa2, Ndufc2, Ndufb8, Ndufs6, Ndufa4, Ndufa8, Ndufa5, Ndufa3,* and *Ndufb6*), complex II (*Sdha*), complex III (*Uqcrh, Uqcrq,* and *Uqcr10*), and complex IV (*Cox5a, Cox7a2, Cox7b, Cox6a1, Cox4i1, Cox6c, Cox5b, Cox6b1,* and *Cox7c*) genes as well as genes encoding other mitochondrial enzymes involved in FA oxidation (FAO, e.g. *Eci1*) were upregulated. Finally, downregulated DEG genes in PLCγ2-P522R KI microglia related to inflammatory response, TLR-signaling and IFN-α response included e.g., *Tlr9, Tlr3, P2rx4, P2rx7,* and *Casp8* (**Figure 9B**).

**Figure 9.**
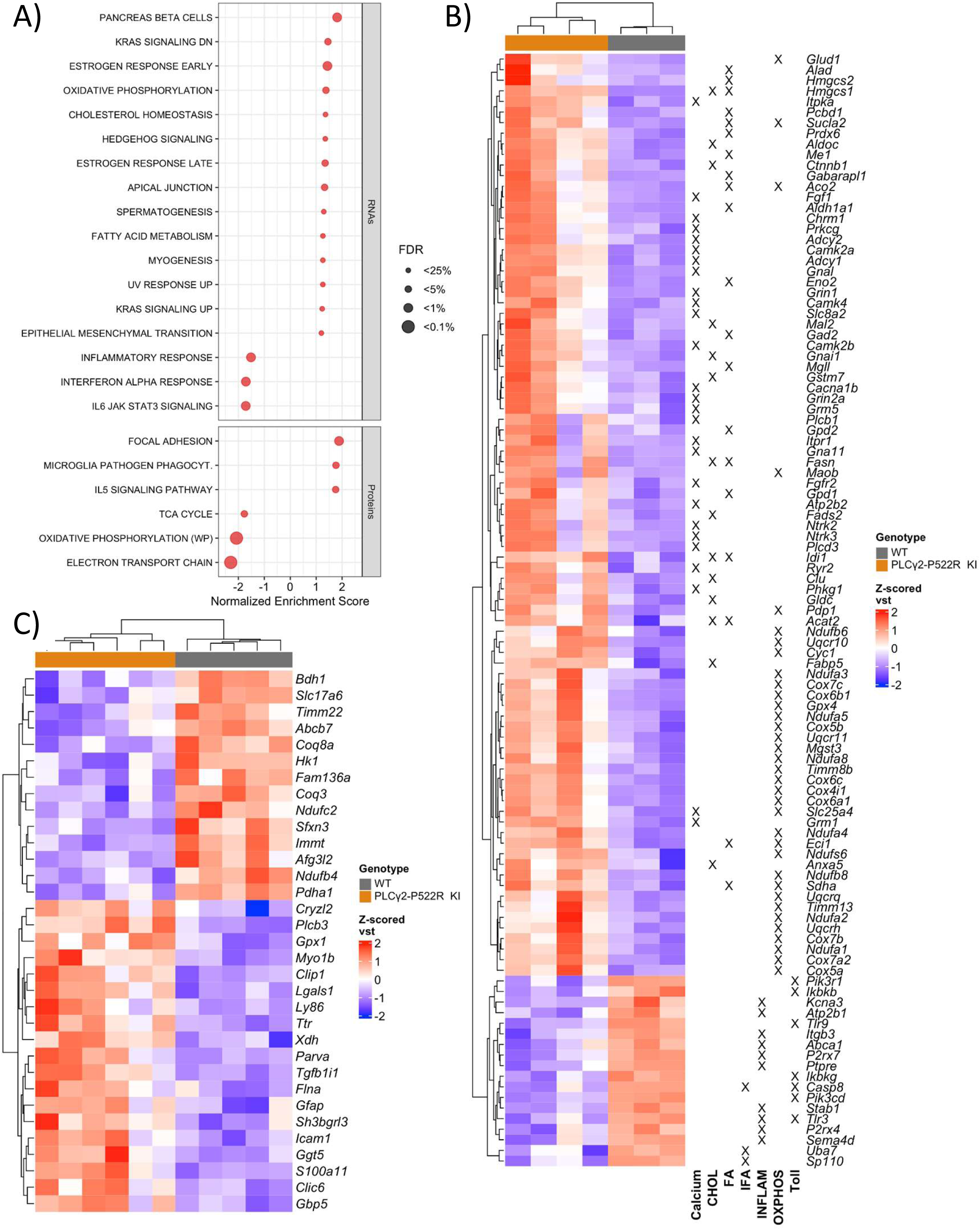
Fatty acid metabolism and mitochondrial function-related targets are upregulated in PLCγ2-P522R KI mouse microglia. A) A dot plot of normalized enrichment scores for enriched and depleted gene sets in enrichment analyses by GSEA for gene (MSigDB Hallmarks, FDR<0.25) and protein (Wikipathways, FDR<0.05) expression in CD11b+ microglia isolated from 13-month-old PLCγ2-P522R KI and wildtype (WT) mice. B) A heatmap of z-scored vst-normalized expression of up-and downregulated differentially expressed genes (DEGs) that were core enrichment genes in a GSEA analysis of gene sets for calcium signaling (Calcium), cholesterol homeostasis (CHOL), fatty acid metabolism (FA), IFN-α response (IFA), inflammatory response (INFLAM), oxidative phosphorylation (OXPHOS), and Toll-like receptor signaling (Toll). C) A heatmap of z-scored vst-normalized expression of significantly up– and downregulated differentially expressed proteins (DEPs) derived from a proteomics study. RNA n(WT)=3, n(PLCγ2-P522R KI)=4, protein n(WT)=5, n(PLCγ2-P522R KI)=6.

Proteomics analysis revealed 34 differentially expressed proteins (DEPs, FDR 0.05) out of which 14 were downregulated and 20 were upregulated in PLCγ2-P522R KI microglia as compared to WT cells (**Figure 9C, S6B**). DEPs and biological pathways only mildly correlated with changes detected in gene expression. Opposite to gene expression, core enrichment proteins related to oxidative phosphorylation were downregulated in the proteomics data set **(Figure 9C**). In contrast, proteins related to FA oxidation showed mild upregulation in PLCγ2-P522R KI microglia, which was similar to that observed at the gene expression level (**Figure S6D**). Together, RNA and proteomics analyses suggest that PLCγ2-P522R modulates pathways related to calcium signaling, inflammatory/IFN response, and mitochondrial FAO and/or FA metabolism in acutely isolated adult mouse microglia.

### PLCγ2-P522R variant improves mitochondrial function in human microglia models

To assess if PLCγ2-P522R-driven mechanisms are translated to human microglia models, we conducted RNA sequencing of the human MDMi cells derived from PLCγ2-P522R variant carriers (CG) and matched control individuals (CC) in untreated conditions and after myelin and LPS treatments. Only 11 significantly altered DEGs were identified in untreated and 10 DEGs in myelin-treated condition in CG as compared to CC MDMi cells (**Figure 10A-B**). LPS had the most robust effect on gene expression, resulting in 236 DEGs in CG as compared to CC cells (**Figure 10C**). In the LPS treated condition, gene enrichment analysis revealed upregulation of oxidative phosphorylation and MYC targets (V1 and V2), while pathways related to inflammatory response, TNF-α signaling, IFN-γ response, and KRAS signaling were most clearly downregulated in MDMis of CG carriers (**Figure 10D**). Core enrichment genes in the oxidative phosphorylation pathway included *ACADVL,* an enzyme regulating FA oxidation, mitochondrial complex I gene, *NDUFB7,* mitochondrial ATP synthase, *ATP5MC1*, and *FXN,* and a gene associated with mitochondrial iron transport and respiration (**Figure 10E**). Upregulated genes linked to MYC target pathway were involved in RNA regulation (*FARSA, MRTO4,* and *WDR74*). Several IFN (*IRF9, IRF4, STAT4*), TNF (*TNFAIP2, TNFAIP3, TNFAIP6*) and IL (*IL1-α, IL-7R*) family members, immune sensors (*CD14, TLR8, TLR2, NLRP3*), and related transcription factors (*NFKB1, NFAT5*) were associated with downregulated pathways. To examine whether respective reduction in inflammatory cytokines can be detected at the protein level, IL-6 and TNF-α levels were measured in the conditioned medium of LPS-treated MDMi cells. However, no significant changes were found in the levels of either of the cytokines (**Figure S7A**). Together with findings in mouse CD11b+ microglia, these data confirm a central role for PLCγ2-P522R in the regulation of genes associated with mitochondrial function and FA oxidation as well as inflammatory/IFN responses.

**Figure 10.**
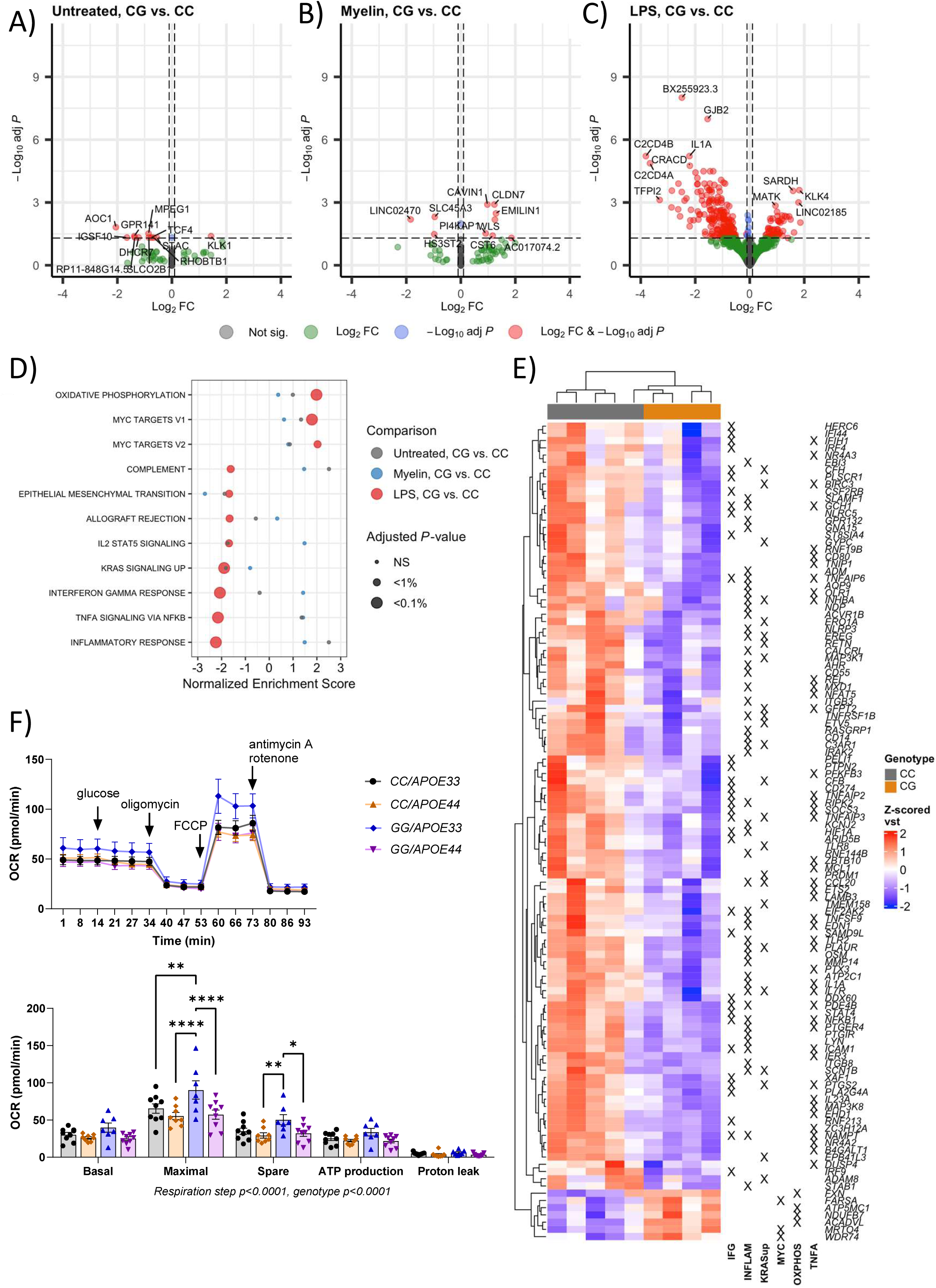
Mitochondrial function is improved in human microglia models generated from blood and skin biopsy samples of the PLCγ2-P522R variant carriers. Volcano plot of differentially expressed genes (DEGs) in blood monocyte-derived microglia-like cells (MDMi) of PLCγ2-P522R variant carriers (CG) and matched controls (CC) in A) untreated (UNT), B) lipopolysaccharide (LPS), and C) myelin-treated conditions. FDR<0.05. Horizontal dashed line: adjusted p-value 0.05; vertical dashed lines: |log2FC|=0.1. D) A dot plot of normalized enrichment scores for enriched and depleted gene sets in enrichment analyses by GSEA for gene (Hallmarks, FDR<0.25) expression in MDMi cells of CG and CC individuals upon UNT, LPS, and myelin treatments. E) A heatmap of z-scored vst-normalized expression of up– and downregulated DEGs in LPS-treated condition that were core enrichment genes in a GSEA analysis of gene sets for IFN-γ response (IFG), inflammatory response (INFLAM), Kras signaling up (KRASup), MYC-targets (MYC), oxidative phosphorylation (OXPHOS), and TNF-α signaling (TNFA). UNT n(CC)=5, n(CG)=4, Myelin n(CC)=5, n(CG)=4, LPS n(CC)=5, n(CG)=4. F) Mitochondrial oxygen consumption rate (OCR, maximal respiration and spare respiratory capacity) is increased in homozygous PLCγ2-P522R (GG) induced pluripotent stem cell-derived microglia (iMGL) as compared to isogenic controls (CC) having *APOE33,* but not *APOE44* genetic background. iMGL n=1 line per group, 7-9 technical replicates per line. Two-way ANOVA with Tukey’s multiple comparisons test. Data are presented as mean ± SEM.

To further investigate mitochondrial function in human microglia cells, we employed PLCγ2-P522R homozygous (GG) iMGLs and isogenic control lines (CC) with *APOE33* or *APOE44* backgrounds (**Figure S7B-C**). In general, mitochondrial OCR was significantly different between the genotypes (p<0.0001). Specifically, maximal respiration was higher in PLCγ2-P522R *GG*/*APOE33* iMGLs as compared to all the other groups (*CC/APOE33* p=0.001, *CC/APOE44* p<0.0001, *CG/APOE4* p<0.0001) (**Figure 10F**). In addition, spare respiratory capacity was significantly higher in *GG*/*APOE33* cells as compared to *CC/APOE44* (p=0.009) and *GG/APOE44* (p=0.027) iMGLs. PLCγ2-P522R did not, however, increase mitochondrial respiration in the *APOE44* background. Taken together, both mouse and human data in this study point towards improved lipid/FA metabolism and mitochondrial function in PLCγ2-P522R-carrying microglia.

## DISCUSSION

PLCγ2-P522R is a well-established variant that reduces the risk for LOAD and recently shown to decrease β-amyloid pathology in the 5XFAD mouse model of AD (Bellenguez, et al., 2019; Sims et al., 2017; Tsai et al., 2023). In line with these findings, we here demonstrate that PLCγ2-P522R decreases β-amyloid plaque burden also in the brain of APP/PS1 mice, a less aggressive mouse model of AD. This was associated with increased microglia activation and microglia clustering around β-amyloid plaques as well as decreased formation of β-amyloid plaque-associated neuronal dystrophies. Importantly, at the mechanistic level, PLCγ2-P522R variant upregulated pathways related to lipid/FA metabolism and mitochondrial function in several different mouse and human microglia models. Conversely, signaling pathways related to inflammatory response were downregulated by the protective PLCγ2-P522R variant.

In the 5XFAD mice, PLCγ2-P522R reduced the size of β-amyloid plaques and shifted plaque morphology from diffuse to a more compacted form (Tsai et al., 2023). In the brain tissue of 13-month-old APP/PS1 mice, PLCγ2-P522R variant reduced the number of X-34-positive compact plaques. On the contrary, the ratio of 6E10-positive diffuse plaques and X-34-positive compact plaques as well as sizes of the individual plaques remained unaffected. Furthermore, in contrast to what was reported in 5XFAD mice, PLCγ2-P522R variant did not change colocalization of 6E10 and IBA1, suggesting that phagocytosis of diffuse β-amyloid remained unaltered in these mice. Yet, we found a significant increase in microglia clustering around β-amyloid plaques, which was accompanied by the reduced number of dystrophic neurites, indicative of decreased β-amyloid-driven axonal damage (Sadleir et al., 2016). Together with increased microglia PET signal in the brain tissue of 13-month-old mice, these findings suggest that PLCγ2-P522R variant enhances microglia activation and shifts them to a more β-amyloid-responsive state. This then leads to an enhanced barrier formation around β-amyloid plaques, which exerts protection to the surrounding neuronal population. These findings suggest that the protective PLCγ2-P522R variant may exert similar beneficial effects on the functions of microglia as those previously reported with a TREM2-activating antibody in different AD disease models (Van Lengerich et al., 2023).

Although PLCγ2-P522R reduced β-amyloid deposition and associated neuronal pathologies, there was no evident improvement in learning and memory functions in the APP/PS1 mice. The most robust manifestation in the behavioral phenotype associated with the PLCγ2-P522R was observed in the passive avoidance task. The tendency of PLCγ2-P522R KI mice to rush into the dark despite having a mild foot shock a day earlier can stem from changes in four CNS systems that regulate (1) pain threshold, (2) spontaneous motor activity, (3) memory, and (4) level of anxiety. The threshold to react to an electric foot shock did not differ between the genotypes. The total distance traversed in a new test box did not differ either. The ‘gold standard’ spatial memory test, the Morris swim task, did reveal a robust impairment associated with the APP/PS1 genotype, but no effect by the PLCγ2-P522R. By excluding these issues, changes in the anxiety level remain the most plausible explanation for the passive avoidance test results. This interpretation is supported by direct evidence of increased anxiety in the PLCγ2-P522R KI mice. First, before the passive avoidance memory testing, we observed that the PLCγ2-P522R KI mice spent significantly less time in the lit compartment than other groups. Second, after receiving the foot shock, PLCγ2-P522R KI mice displayed a crab-like walking backwards that can be interpreted as an attempt to retreat from the daunting environment. Third, during the activity monitoring, the PLCγ2-P522R KI mice avoided the arena center more than the control mice did. Fear and anxiety are essential emotions that aid the survival of species like the mouse living under constant threat by predators in the wild (Silva et al., 2016). Furthermore, there is increasing evidence in that microglia activation may be a key driver of anxiety phenotype in mice (Lehmann et al., 2019). Together, these findings suggest that the PLCγ2-P522R variant promotes anxiety phenotype in APP/PS1 mice, which may promote longevity via protection against environmental threats through mechanisms associated with altered microglia functions.

We have previously shown that PLCγ2-P522R KI mouse macrophages respond more efficiently to LPS-induced acute inflammation than WT cells (Takalo et al., 2020). In line with the initial finding, PLCγ2-P522R KI mouse primary microglia secreted significantly more IL-6 and TNF-α than WT cells when treated with a high dose of LPS (1 µg/ml). In contrast, there was a general trend towards lower inflammatory cytokine levels in the temporo-occipital cortex and hippocampus of A+/P^ki/ki^ mice as compared to A+/P^wt/wt^ mice. Furthermore, RNA sequencing revealed the downregulation of inflammatory and interferon signaling pathways in CD11b+ microglia isolated from 13-month-old mice without AD-associated stress as well as in human MDMi cells after a low dose of LPS. This suggests that while the PLCγ2-P522R variant enhances an acute response to strong inflammatory stimuli (1µg/ml LPS) in mouse cells *in vitro*, it suppresses chronic pro-inflammatory activation associated with β-amyloid deposition, aging, or weaker inflammatory stimulus (200 ng/ml LPS. These findings are partly different from a recent report showing that PLCγ2-P522R increases the levels of IL-1β and IL-5 in 5XFAD mouse cortex (Tsai et al., 2023). These contradicting findings may be explained by differences between the background of the mouse lines and the severity of β-amyloid pathology at the time of the analyses. Collectively, the protective PLCγ2-P522R variant is most likely centrally involved in the regulation of inflammatory response, although the effect size and direction may be context-dependent (acute vs. chronic).

Young microglia rely on flexible use of diverse energy sources, including FAs, to efficiently perform numerous immune functions and to maintain tissue homeostasis (Fairley et al., 2021). In energy-demanding conditions, FAs are released from their main intracellular lipid storage units known as LDs, which mainly contain triglycerides and cholesterol esters. Free FAs are then transported to mitochondria where they are used to fuel mitochondrial oxidative phosphorylation in a process called FA β-oxidation (FAO) (Fairley et al., 2021). Conversely, aged microglia present deterioration of mitochondrial efficiency and downregulation of oxidative phosphorylation-related genes (Hickman et al., 2013; López-Otfn et al., 2013). Depletion of either TREM2 or PLCγ2 has been shown to increase lipid accumulation and impair mitochondrial respiration, thus specifically influencing FAO (Andreone et al., 2020; Van Lengerich et al., 2023; Wei et al., 2024). This suggests that TREM2-PLCγ2 signaling is needed for maintaining microglial energy capacity and thereby slowing down ageing-related deterioration. Accordingly, reduced accumulation of large LDs was observed in the present study in PLCγ2-P522R KI mouse microglia upon myelin and, to a lesser extent, LPS treatments. The buildup of intracellular LDs is known to drive microglia dysfunction as presented by deficient phagocytosis, increased ROS production, and chronic proinflammatory response, which are the common phenotypes increasingly detected in the ageing microglia (Marschallinger et al., 2020; Prakash et al., 2023). Simultaneous increase in the acute response to LPS treatment and decrease in cellular stress, exhibited as reduced LDH and ROS levels, supports the idea that the protective PLCγ2-P522R variant suppresses the ageing-related phenotype in mouse microglia.

Based on the end point measurement exclusively, it is impossible to conclusively determine whether PLCγ2-P522R KI influences LD biogenesis, breakdown, or other aspects of LD dynamics. Genes encoding fatty acid (FA)-binding proteins (*Fabp3, Fabp5, Fabp7*), which are involved in LD lipolysis and transporting FAs into different organelles, including mitochondria (Senga et al., 2018), were significantly upregulated in the adult PLCγ2-P522R KI mouse microglia, suggesting that the PLCγ2-P522R variant might associate with the breakdown of LDs. Furthermore, we observed increased expression of *Mgll,* a gene encoding monoacylglycerol lipase, which catalyzes the final step in the hydrolysis of triglycerides to FAs (Tardelli, 2020). Although we did not find direct functional evidence for improved mitochondrial function or glycolysis in cultured PLCγ2-P522R mouse microglia, it is not completely ruled out that the PLCγ2-P522R variant specifically increases FAO for the energy production. In fact, DEGs (e.g. *Eci1*) and biological pathways (β-oxidation and FAO) arising from RNA sequencing and proteomics data directly point towards enhanced FAO in the adult PLCγ2-P522R KI mouse microglia. Together with the findings of a study showing that PLCγ2 KO impairs FAO in human iMGLs (Van Lengerich et al., 2023), these data suggest that the PLCγ2-P522R variant reinforces microglial energy metabolism through improved lipid recycling. This notion is further supported by transcriptomic and functional evidence from the human microglia models, which show upregulation of oxidative phosphorylation, FAO, and mitochondrial respiration-related genes in the MDMi cells of the PLCγ2-P522R variant carriers and enhanced mitochondrial respiration in homozygous PLCγ2-P522R iMGL lines with *APOE3/3* background. Altogether, the transcriptomic and functional observations in the mouse and human models point towards increased mitochondrial functions, FAO, and lipid metabolism and further suggest that the beneficial effects of the protective PLCγ2-P522R variant may be mediated through an anti-ageing mechanism.

PI3K-AKT signaling is another key regulator of lipid metabolism and PI3K inhibitors have been shown to efficiently block the accumulation of large LDs in numerous studies and several study models (Haney et al., 2024; Khatchadourian et al., 2012; C. B. Ryan et al., 2024). In this context, we have previously shown that AKT activation was decreased in PLCγ2-P522R mouse brain, possibly due to an increased consumption of PIP_2_ by PLCγ2 and consequently reduced PIP_3_ resources for promoting PI3K-AKT-signalling (Takalo et al., 2020). In line with this finding, RNA sequencing revealed that several genes encoding PI3K subunits were downregulated in the adult PLCγ2-P522R KI mouse microglia as compared to WT microglia. Although further studies are needed to confirm this, it is possible that the reduced accumulation of LDs observed in the present study is at least partially mediated via the inhibition of PI3K-AKT-signaling, resulting from the increased PIP_2_ consumption and/or reduced expression of PI3K subunit components.

### Conclusions

Our present study shows that the AD protective PLCγ2-P522R variant reduces the β-amyloid deposition and associated neuronal pathologies in the APP/PS1 model mice via enhancing microglia activation and response towards β-amyloid deposits, while suppressing their proinflammatory phenotype. These changes are mechanistically linked to improved lipid/FA metabolism and mitochondrial functions, which were observed in different human and mouse microglia models. Although further studies are warranted, our findings suggest that the protective effects of the PLCγ2-P522R variant upon AD-associated stress conditions are consequences of slower metabolic deterioration related to microglial ageing.

## DECLARATIONS

### Ethics approval and consent to participate

Animals were raised and handled at the Laboratory Animal Center of the University of Eastern Finland, Kuopio, Finland. All mouse studies were carried out in accordance with the guidelines of the European Community Council Directives 86/609/EEC and approved by the Regional State Administrative Agency and Project Authorisation Board (ESAVI/7315/2024, EKS-004-2019).

All study protocols concerning human samples were approved by Medical Research Ethics Committee of Wellbeing Services County of North Savo (formerly Medical Research Ethics Committee of North Savo Hospital District, 833/2023, 123/2016). Participant recontacting was approved by the Scientific Steering Committee of Auria Biobank (BB_2020-0062).

### Consent for publication

All authors have approved the final version of the manuscript and give their consent for publication.

### Availability of data and materials

Datasets generated and/or analyzed in this manuscript are included within the article and its additional files or are available from the corresponding author on reasonable request.

### Authors’ contributions

Conceptualization and study design MT, MH; Methodology MT, HJ, TR, MHe, IK, SH, KJ, PP, SM, EMK, RMW, DH, HJä, TK, TN, SK, MP, HM, AH, SL, HT, CH, MH; Investigation: MT, HJ, TR, IK, MHe, HKoi, KJ, SM, PM, SPJ, RS, DH; Formal analysis: MT, HJ, TR, MHe, SH, KJ, SM, DH, HT; Visualization: MT, SH; Resources: MT, EMK, PP, KF, AH, VL, AR, SL, HT, CH, JK, MH; Funding acquisition: MT, SL, CH, JK, MH; Project administration: MT; Supervision: MT, MH; Writing – original draft: MT; Writing – review & editing: All authors.

## Acknowledgements

The authors would like to thank University of Eastern Finland Cell and Tissue Imaging Unit (University of Eastern Finland, Kuopio, Finland) for providing the infrastructure and help for microscopy as well as Biomedical Imaging Unit (University of Eastern Finland, Kuopio, Finland) for carrying out *in vivo* PET imaging. We thank Mrs. Maarit Pulkkinen for her expert technical assistance with PET imaging. We acknowledge the Biocenter Finland and Biocenter Kuopio Phenotyping Center (University of Eastern Finland, Kuopio, Finland) for the service in behavioral testing and analysis and UEF Bioinformatics Center (University of Eastern Finland, Kuopio, Finland) for providing servers for computational analyses. The authors also want to thank the Auria Biobank and Dr. Merja Perälä for recontacting biobank sample donors.

## References

1. Abud, E. M., Ramirez, R. N., Martinez, E. S., Healy, L. M., Nguyen, C. H. H., Newman, S. A., Yeromin, A. V., Scarfone, V. M., Marsh, S. E., Fimbres, C., Caraway, C. A., Fote, G. M., Madany, A. M., Agrawal, A., Kayed, R., Gylys, K. H., Cahalan, M. D., Cummings, B. J., Antel, J. P., … Blurton-Jones, M. (2017). iPSC-Derived Human Microglia-like Cells to Study Neurological Diseases. Neuron, 94(2), 278–293.e9. 10.1016/j.neuron.2017.03.042

2. Andreone, B. J., Przybyla, L., Llapashtica, C., Rana, A., Davis, S. S., Van Lengerich, B., Lin, K., Shi, J., Mei, Y., Astarita, G., Di Paolo, G., Sandmann, T., Monroe, K. M., & Lewcock, J. W. (2020). Alzheimer’s-associated PLCγ2 is a signaling node required for both TREM2 function and the inflammatory response in human microglia. Nature Neuroscience, 23(8), 927–938. 10.1038/s41593-020-0650-6

3. ARUK Consortium, GERAD/PERADES, CHARGE, ADGC, EADI, Sims, R., Van Der Lee, S. J., Naj, A. C., Bellenguez, C., Badarinarayan, N., Jakobsdottir, J., Kunkle, B. W., Boland, A., Raybould, R., Bis, J. C., Martin, E. R., Grenier-Boley, B., Heilmann-Heimbach, S., Chouraki, V., Kuzma, A. B., Sleegers, K., Vronskaya, M., … Schellenberg, G. D. (2017). Rare coding variants in PLCG2, ABI3, and TREM2 implicate microglial-mediated innate immunity in Alzheimer’s disease. Nature Genetics, 49(9), 1373–1384. 10.1038/ng.3916

4. Bill, C. A., & Vines, C. M. (2020). Phospholipase C. In Md. S. Islam (Ed.), Calcium Signaling (Vol. 1131, pp. 215–242). Springer International Publishing. 10.1007/978-3-030-12457-1_9

5. Bolger, A. M., Lohse, M., & Usadel, B. (2014). Trimmomatic: A flexible trimmer for Illumina sequence data. Bioinformatics, 30(15), 2114–2120. 10.1093/bioinformatics/btu170

6. Coulon, A., Rabiller, F., Takalo, M., Roy, A., Martiskainen, H., Siedlecki-Wullich, D., Mendes, T., Lemeu, C., Carvalho, L.-I., Ehrardt, A., Melo De Farias, A. R., Hulsman, M., Najdek, C., Lannette-Weimann, N., Freire-Regatillo, A., Amouyel, P., Charbonnier, C., Dols-Icardo, O., Jeskanen, H., … Lambert, J.-C. (2024). Neuronal downregulation of PLCG2 impairs synaptic function and elicits Alzheimer disease hallmarks. 10.1101/2024.04.29.591575

7. Cox, J., Hein, M. Y., Luber, C. A., Paron, I., Nagaraj, N., & Mann, M. (2014). Accurate Proteome-wide Label-free Quantification by Delayed Normalization and Maximal Peptide Ratio Extraction, Termed MaxLFQ. Molecular & Cellular Proteomics, 13(9), 2513–2526. 10.1074/mcp.M113.031591

8. Demichev, V., Messner, C. B., Vernardis, S. I., Lilley, K. S., & Ralser, M. (2020). DIA-NN: Neural networks and interference correction enable deep proteome coverage in high throughput. Nature Methods, 17(1), 41–44. 10.1038/s41592-019-0638-x

9. Dobin, A., Davis, C. A., Schlesinger, F., Drenkow, J., Zaleski, C., Jha, S., Batut, P., Chaisson, M., & Gingeras, T. R. (2013). STAR: Ultrafast universal RNA-seq aligner. Bioinformatics, 29(1), 15–21. 10.1093/bioinformatics/bts635

10. Fairley, L. H., Wong, J. H., & Barron, A. M. (2021). Mitochondrial Regulation of Microglial Immunometabolism in Alzheimer’s Disease. Frontiers in Immunology, 12, 624538. 10.3389/fimmu.2021.624538

11. Haney, M. S., Pálovics, R., Munson, C. N., Long, C., Johansson, P. K., Yip, O., Dong, W., Rawat, E., West, E., Schlachetzki, J. C. M., Tsai, A., Guldner, I. H., Lamichhane, B. S., Smith, A., Schaum, N., Calcuttawala, K., Shin, A., Wang, Y.-H., Wang, C., … Wyss-Coray, T. (2024). APOE4/4 is linked to damaging lipid droplets in Alzheimer’s disease microglia. Nature, 628(8006), 154–161. 10.1038/s41586-024-07185-7

12. Hansen, D. V., Hanson, J. E., & Sheng, M. (2018). Microglia in Alzheimer’s disease. Journal of Cell Biology, 217(2), 459–472. 10.1083/jcb.201709069

13. Hickman, S. E., Kingery, N. D., Ohsumi, T. K., Borowsky, M. L., Wang, L., Means, T. K., & El Khoury, J. (2013). The microglial sensome revealed by direct RNA sequencing. Nature Neuroscience, 16(12), 1896–1905. 10.1038/nn.3554

14. Jankowsky, J. L., Fadale, D. J., Anderson, J., Xu, G. M., Gonzales, V., Jenkins, N. A., Copeland, N. G., Lee, M. K., Younkin, L. H., Wagner, S. L., Younkin, S. G., & Borchelt, D. R. (2004). Mutant presenilins specifically elevate the levels of the 42 residue β-amyloid peptide in vivo: Evidence for augmentation of a 42-specific γ secretase. Human Molecular Genetics, 13(2), 159–170. 10.1093/hmg/ddh019

15. Keren-Shaul, H., Spinrad, A., Weiner, A., Matcovitch-Natan, O., Dvir-Szternfeld, R., Ulland, T. K., David, E., Baruch, K., Lara-Astaiso, D., Toth, B., Itzkovitz, S., Colonna, M., Schwartz, M., & Amit, I. (2017). A Unique Microglia Type Associated with Restricting Development of Alzheimer’s Disease. Cell, 169(7), 1276–1290.e17. 10.1016/j.cell.2017.05.018

16. Khatchadourian, A., Bourque, S. D., Richard, V. R., Titorenko, V. I., & Maysinger, D. (2012). Dynamics and regulation of lipid droplet formation in lipopolysaccharide (LPS)-stimulated microglia. Biochimica et Biophysica Acta (BBA) – Molecular and Cell Biology of Lipids, 1821(4), 607–617. 10.1016/j.bbalip.2012.01.007

17. Kleineidam, L., Chouraki, V., Próchnicki, T., van der Lee, S. J., Madrid-Márquez, L., Wagner-Thelen, H., Karaca, I., Weinhold, L., Wolfsgruber, S., Boland, A., Martino Adami, P. V., Lewczuk, P., Popp, J., Brosseron, F., Jansen, I. E., Hulsman, M., Kornhuber, J., Peters, O., Berr, C., … Ramirez, A. (2020). PLCG2 protective variant p.P522R modulates tau pathology and disease progression in patients with mild cognitive impairment. Acta Neuropathologica, 139(6), 1025–1044. 10.1007/s00401-020-02138-6

18. Koskuvi, M., Pörsti, E., Hewitt, T., Räsänen, N., Wu, Y.-C., Trontti, K., McQuade, A., Kalyanaraman, S., Ojansuu, I., Vaurio, O., Cannon, T. D., Lönnqvist, J., Therman, S., Suvisaari, J., Kaprio, J., Blurton-Jones, M., Hovatta, I., Lähteenvuo, M., Rolova, T., … Koistinaho, J. (2024). Genetic contribution to microglial activation in schizophrenia. Molecular Psychiatry. 10.1038/s41380-024-02529-1

19. Kunkle, B. W., Alzheimer Disease Genetics Consortium (ADGC), The European Alzheimer’s Disease Initiative (EADI), Cohorts for Heart and Aging Research in Genomic Epidemiology Consortium (CHARGE), Genetic and Environmental Risk in AD/Defining Genetic, Polygenic and Environmental Risk for Alzheimer’s Disease Consortium (GERAD/PERADES), Grenier-Boley, B., Sims, R., Bis, J. C., Damotte, V., Naj, A. C., Boland, A., Vronskaya, M., Van Der Lee, S. J., Amlie-Wolf, A., Bellenguez, C., Frizatti, A., Chouraki, V., Martin, E. R., Sleegers, K., … Pericak-Vance, M. A. (2019). Genetic meta-analysis of diagnosed Alzheimer’s disease identifies new risk loci and implicates Aβ, tau, immunity and lipid processing. Nature Genetics, 51(3), 414–430. 10.1038/s41588-019-0358-2

20. Kurki, M. I., Karjalainen, J., Palta, P., Sipilä, T. P., Kristiansson, K., Donner, K. M., Reeve, M. P., Laivuori, H., Aavikko, M., Kaunisto, M. A., Loukola, A., Lahtela, E., Mattsson, H., Laiho, P., Della Briotta Parolo, P., Lehisto, A. A., Kanai, M., Mars, N., Rämö, J., … Palotie, A. (2023). FinnGen provides genetic insights from a well-phenotyped isolated population. Nature, 613(7944), 508–518. 10.1038/s41586-022-05473-8

21. Larocca, J. N., & Norton, W. T. (2006). Isolation of Myelin. Current Protocols in Cell Biology, 33(1). 10.1002/0471143030.cb0325s33

22. Lehmann, M. L., Weigel, T. K., Poffenberger, C. N., & Herkenham, M. (2019). The Behavioral Sequelae of Social Defeat Require Microglia and Are Driven by Oxidative Stress in Mice. The Journal of Neuroscience, 39(28), 5594–5605. 10.1523/JNEUROSCI.0184-19.2019

23. Liao, Y., Smyth, G. K., & Shi, W. (2019). The R package Rsubread is easier, faster, cheaper and better for alignment and quantification of RNA sequencing reads. Nucleic Acids Research, 47(8), e47–e47. 10.1093/nar/gkz114

24. Liberzon, A., Birger, C., Thorvaldsdóttir, H., Ghandi, M., Mesirov, J. P., & Tamayo, P. (2015). The Molecular Signatures Database Hallmark Gene Set Collection. Cell Systems, 1(6), 417–425. 10.1016/j.cels.2015.12.004

25. Liddelow, S. A., Guttenplan, K. A., Clarke, L. E., Bennett, F. C., Bohlen, C. J., Schirmer, L., Bennett, M. L., Münch, A. E., Chung, W.-S., Peterson, T. C., Wilton, D. K., Frouin, A., Napier, B. A., Panicker, N., Kumar, M., Buckwalter, M. S., Rowitch, D. H., Dawson, V. L., Dawson, T. M., … Barres, B. A. (2017). Neurotoxic reactive astrocytes are induced by activated microglia. Nature, 541(7638), 481–487. 10.1038/nature21029

26. López-Otfn, C., Blasco, M. A., Partridge, L., Serrano, M., & Kroemer, G. (2013). The Hallmarks of Aging. Cell, 153(6), 1194–1217. 10.1016/j.cell.2013.05.039

27. Love, M. I., Huber, W., & Anders, S. (2014). Moderated estimation of fold change and dispersion for RNA-seq data with DESeq2. Genome Biology, 15(12), 550. 10.1186/s13059-014-0550-8

28. Magno, L., Lessard, C. B., Martins, M., Lang, V., Cruz, P., Asi, Y., Katan, M., Bilsland, J., Lashley, T., Chakrabarty, P., Golde, T. E., & Whiting, P. J. (2019). Alzheimer’s disease phospholipase C-gamma-2 (PLCG2) protective variant is a functional hypermorph. Alzheimer’s Research & Therapy, 11(1), 16. 10.1186/s13195-019-0469-0

29. Marschallinger, J., Iram, T., Zardeneta, M., Lee, S. E., Lehallier, B., Haney, M. S., Pluvinage, J. V., Mathur, V., Hahn, O., Morgens, D. W., Kim, J., Tevini, J., Felder, T. K., Wolinski, H., Bertozzi, C. R., Bassik, M. C., Aigner, L., & Wyss-Coray, T. (2020). Lipid-droplet-accumulating microglia represent a dysfunctional and proinflammatory state in the aging brain. Nature Neuroscience, 23(2), 194–208. 10.1038/s41593-019-0566-1

30. Martiskainen, H., Willman, R.-M., Heikkinen, S., Müller, S. A., Sinisalo, R., Takalo, M., Mäkinen, P., Kuulasmaa, T., Pekkala, V., Galván Del Rey, A., Harju, P., Juopperi, S.-P., Jeskanen, H., Kervinen, I., Saastamoinen, K., FinnGen, Niiranen, M., Heikkinen, S. V., Kurki, M. I., … Hiltunen, M. (2024). Monoallelic TYROBP deletion is a novel risk factor for Alzheimer’s disease. 10.1101/2024.05.09.24307099

31. McQuade, A., Coburn, M., Tu, C. H., Hasselmann, J., Davtyan, H., & Blurton-Jones, M. (2018). Development and validation of a simplified method to generate human microglia from pluripotent stem cells. Molecular Neurodegeneration, 13(1), 67. 10.1186/s13024-018-0297-x

32. Parhizkar, S., Arzberger, T., Brendel, M., Kleinberger, G., Deussing, M., Focke, C., Nuscher, B., Xiong, M., Ghasemigharagoz, A., Katzmarski, N., Krasemann, S., Lichtenthaler, S. F., Müller, S. A., Colombo, A., Monasor, L. S., Tahirovic, S., Herms, J., Willem, M., Pettkus, N., … Haass, C. (2019). Loss of TREM2 function increases amyloid seeding but reduces plaque-associated ApoE. Nature Neuroscience, 22(2), 191–204. 10.1038/s41593-018-0296-9

33. Peitz, M., Bechler, T., Thiele, C. C., Veltel, M., Bloschies, M., Fliessbach, K., Ramirez, A., & Brüstle, O. (2018). Blood-derived integration-free iPS cell line UKBi011-A from a diagnosed male Alzheimer’s disease patient with APOE ɛ4/ɛ4 genotype. Stem Cell Research, 29, 250–253. 10.1016/j.scr.2018.04.011

34. Prakash, P., Manchanda, P., Paouri, E., Bisht, K., Sharma, K., Wijewardhane, P. R., Randolph, C. E., Clark, M. G., Fine, J., Thayer, E. A., Crockett, A., Gasmi, N., Stanko, S., Prayson, R. A., Zhang, C., Davalos, D., & Chopra, G. (2023). Amyloid β Induces Lipid Droplet-Mediated Microglial Dysfunction in Alzheimer’s Disease. 10.1101/2023.06.04.543525

35. Ryan, C. B., Choi, J. S., Kang, B., Herr, S., Pereira, C., Moraes, C. T., Al-Ali, H., & Lee, J. K. (2024). PI3K signaling promotes formation of lipid-laden foamy macrophages at the spinal cord injury site. Neurobiology of Disease, 190, 106370. 10.1016/j.nbd.2023.106370

36. Ryan, K. J., White, C. C., Patel, K., Xu, J., Olah, M., Replogle, J. M., Frangieh, M., Cimpean, M., Winn, P., McHenry, A., Kaskow, B. J., Chan, G., Cuerdon, N., Bennett, D. A., Boyd, J. D., Imitola, J., Elyaman, W., De Jager, P. L., & Bradshaw, E. M. (2017). A human microglia-like cellular model for assessing the effects of neurodegenerative disease gene variants. Science Translational Medicine, 9(421), eaai7635. 10.1126/scitranslmed.aai7635

37. Sadleir, K. R., Kandalepas, P. C., Buggia-Prévot, V., Nicholson, D. A., Thinakaran, G., & Vassar, R. (2016). Presynaptic dystrophic neurites surrounding amyloid plaques are sites of microtubule disruption, BACE1 elevation, and increased Aβ generation in Alzheimer’s disease. Acta Neuropathologica, 132(2), 235–256. 10.1007/s00401-016-1558-9

38. Safaiyan, S., Kannaiyan, N., Snaidero, N., Brioschi, S., Biber, K., Yona, S., Edinger, A. L., Jung, S., Rossner, M. J., & Simons, M. (2016). Age-related myelin degradation burdens the clearance function of microglia during aging. Nature Neuroscience, 19(8), 995–998. 10.1038/nn.4325

39. Schlepckow, K., Monroe, K. M., Kleinberger, G., Cantuti-Castelvetri, L., Parhizkar, S., Xia, D., Willem, M., Werner, G., Pettkus, N., Brunner, B., Sülzen, A., Nuscher, B., Hampel, H., Xiang, X., Feederle, R., Tahirovic, S., Park, J. I., Prorok, R., Mahon, C., … Haass, C. (2020). Enhancing protective microglial activities with a dual function TREM 2 antibody to the stalk region. EMBO Molecular Medicine, 12(4), e11227. 10.15252/emmm.201911227

40. Schmid, B., Holst, B., Clausen, C., Bahnassawy, L., Reinhardt, P., Bakker, M. H. M., Díaz-Guerra, E., Vicario, C., Martino-Adami, P. V., Thoenes, M., Ramirez, A., Fliessbach, K., Grezella, C., Brüstle, O., Peitz, M., Ebneth, A., & Cabrera-Socorro, A. (2021). Generation of a set of isogenic iPSC lines carrying all APOE genetic variants (Ɛ2/Ɛ3/Ɛ4) and knock-out for the study of APOE biology in health and disease. Stem Cell Research, 52, 102180. 10.1016/j.scr.2021.102180

41. Senga, S., Kobayashi, N., Kawaguchi, K., Ando, A., & Fujii, H. (2018). Fatty acid-binding protein 5 (FABP5) promotes lipolysis of lipid droplets, de novo fatty acid (FA) synthesis and activation of nuclear factor-kappa B (NF-κB) signaling in cancer cells. Biochimica et Biophysica Acta (BBA) – Molecular and Cell Biology of Lipids, 1863(9), 1057–1067. 10.1016/j.bbalip.2018.06.010

42. Silva, B. A., Gross, C. T., & Gräff, J. (2016). The neural circuits of innate fear: Detection, integration, action, and memorization. Learning & Memory, 23(10), 544–555. 10.1101/lm.042812.116

43. Styren, S. D., Hamilton, R. L., Styren, G. C., & Klunk, W. E. (2000). X-34, A Fluorescent Derivative of Congo Red: A Novel Histochemical Stain for Alzheimer’s Disease Pathology. Journal of Histochemistry & Cytochemistry, 48(9), 1223–1232. 10.1177/002215540004800906

44. Subramanian, A., Tamayo, P., Mootha, V. K., Mukherjee, S., Ebert, B. L., Gillette, M. A., Paulovich, A., Pomeroy, S. L., Golub, T. R., Lander, E. S., & Mesirov, J. P. (2005). Gene set enrichment analysis: A knowledge-based approach for interpreting genome-wide expression profiles. Proceedings of the National Academy of Sciences, 102(43), 15545–15550. 10.1073/pnas.0506580102

45. Takalo, M., Wittrahm, R., Wefers, B., Parhizkar, S., Jokivarsi, K., Kuulasmaa, T., Mäkinen, P., Martiskainen, H., Wurst, W., Xiang, X., Marttinen, M., Poutiainen, P., Haapasalo, A., Hiltunen, M., & Haass, C. (2020). The Alzheimer’s disease-associated protective Plcγ2-P522R variant promotes immune functions. Molecular Neurodegeneration, 15(1), 52. 10.1186/s13024-020-00402-7

46. Tardelli, M. (2020). Monoacylglycerol lipase reprograms lipid precursors signaling in liver disease. World Journal of Gastroenterology, 26(25), 3577–3585. 10.3748/wjg.v26.i25.3577

47. Tsai, A. P., Dong, C., Lin, P. B.-C., Oblak, A. L., Viana Di Prisco, G., Wang, N., Hajicek, N., Carr, A. J., Lendy, E. K., Hahn, O., Atkins, M., Foltz, A. G., Patel, J., Xu, G., Moutinho, M., Sondek, J., Zhang, Q., Mesecar, A. D., Liu, Y., … Landreth, G. E. (2023). Genetic variants of phospholipase C-γ2 alter the phenotype and function of microglia and confer differential risk for Alzheimer’s disease. Immunity, 56(9), 2121–2136.e6. 10.1016/j.immuni.2023.08.008

48. Ulrich, J. D., Ulland, T. K., Mahan, T. E., Nyström, S., Nilsson, K. P., Song, W. M., Zhou, Y., Reinartz, M., Choi, S., Jiang, H., Stewart, F. R., Anderson, E., Wang, Y., Colonna, M., & Holtzman, D. M. (2018). ApoE facilitates the microglial response to amyloid plaque pathology. Journal of Experimental Medicine, 215(4), 1047–1058. 10.1084/jem.20171265

49. Van Der Lee, S. J., DESGESCO (Dementia Genetics Spanish Consortium), EADB (Alzheimer Disease European DNA biobank), EADB (Alzheimer Disease European DNA biobank), IFGC (International FTD-Genomics Consortium), IPDGC (The International Parkinson Disease Genomics Consortium), IPDGC (The International Parkinson Disease Genomics Consortium), RiMod-FTD (Risk and Modifying factors in Fronto-Temporal Dementia), Netherlands Brain Bank (NBB), The GIFT (Genetic Investigation in Frontotemporal Dementia and Alzheimer’s Disease) Study Group, Conway, O. J., Jansen, I., Carrasquillo, M. M., Kleineidam, L., Van Den Akker, E., Hernández, I., Van Eijk, K. R., Stringa, N., Chen, J. A., Zettergren, A., Andlauer, T. F. M., … Holstege, H. (2019). A nonsynonymous mutation in PLCG2 reduces the risk of Alzheimer’s disease, dementia with Lewy bodies and frontotemporal dementia, and increases the likelihood of longevity. Acta Neuropathologica, 138(2), 237–250. 10.1007/s00401-019-02026-8

50. Van Lengerich, B., Zhan, L., Xia, D., Chan, D., Joy, D., Park, J. I., Tatarakis, D., Calvert, M., Hummel, S., Lianoglou, S., Pizzo, M. E., Prorok, R., Thomsen, E., Bartos, L. M., Beumers, P., Capell, A., Davis, S. S., De Weerd, L., Dugas, J. C., … Monroe, K. M. (2023). A TREM2-activating antibody with a blood–brain barrier transport vehicle enhances microglial metabolism in Alzheimer’s disease models. Nature Neuroscience. 10.1038/s41593-022-01240-0

51. Wei, W., Zhang, L., Xin, W., Pan, Y., Tatenhorst, L., Hao, Z., Gerner, S. T., Huber, S., Juenemann, M., Butz, M., Huttner, H. B., Bähr, M., Fitzner, D., Jia, F., & Doeppner, T. R. (2024). TREM2 regulates microglial lipid droplet formation and represses post-ischemic brain injury. Biomedicine & Pharmacotherapy, 170, 115962. 10.1016/j.biopha.2023.115962

52. Wilson, A. A., Garcia, A., Parkes, J., McCormick, P., Stephenson, K. A., Houle, S., & Vasdev, N. (2008). Radiosynthesis and initial evaluation of [18F]-FEPPA for PET imaging of peripheral benzodiazepine receptors. Nuclear Medicine and Biology, 35(3), 305–314. 10.1016/j.nucmedbio.2007.12.009

53. Yuan, P., Condello, C., Keene, C. D., Wang, Y., Bird, T. D., Paul, S. M., Luo, W., Colonna, M., Baddeley, D., & Grutzendler, J. (2016). TREM2 Haplodeficiency in Mice and Humans Impairs the Microglia Barrier Function Leading to Decreased Amyloid Compaction and Severe Axonal Dystrophy. Neuron, 92(1), 252–264. 10.1016/j.neuron.2016.09.016

54. Zhu, A., Ibrahim, J. G., & Love, M. I. (2019). Heavy-tailed prior distributions for sequence count data: Removing the noise and preserving large differences. Bioinformatics, 35(12), 2084–2092. 10.1093/bioinformatics/bty895

