## Supplemental files for "Protective variant in PLCγ2 mitigates Alzheimer’s disease-associated pathologies via enhancing beneficial microglia functions"

**IL-6 and TNF-α measurement in conditioned MDMi medium**

For analyzing inflammatory cytokines in the conditioned medium, medium was collected, centrifuged at 10000 x g, for 10 min at +4$^{o}$C, and supernatant was stored at -80$^{o}$C until analysis (below). IL-6 and TNF-α levels in the medium were analyzed with IL6 and TNF alpha Human Uncoated ELISA Kit (#88-7346-22 and #88-7066-22, respectively Thermo Fisher).

**Immunocytochemistry and qPCR-based characterization of microglia markers in iMGL cells**

iMGL cells grown on coverslips coated with Matrigel (1:100) were fixed with 4% formaldehyde for 20 min at RT and stained overnight with rabbit anti-P2Y12 (1:1000, #HPA014518, Merc) and goat anti-TREM2 (1:100 #AF1828, R&D Systems) primary antibodies diluted in PBS containing 1% BSA and 0.05% Tween20 at +4 °C. After 4 x 5min washing in PBS + 0.05% Tween20, the cells were incubated for 1 h with chicken anti-rabbit AlexaFluor 647 (#A21443, ThermoFisher Scientific) and chicken anti-goat AlexaFluor488 (#A21467, ThermoFisher Scientific) secondary antibodies (both diluted 1:500 in PBS containing 1% BSA and 0.05% Tween20) at RT. During the washing step, the cells were further incubated for 5 min with DAPI at 1 µg/ml in PBS to stain the nuclei. The coverslips were mounted on glass slides using Fluoromount-G mounting medium (#0100-01, Southern Biotech), and imaged using Zeiss AxioImager M1 microscope.

iMGL cells for qPCR-based characterization were grown and maturated for 4-5 days on 12-well plates coated with Matrigel (2:100). RNA was isolated by using Qiagen RNeasy Mini Kit (#74106, Qiagen) according to manufacturer's instructions. cDNA was synthetized using Maxima reverse transcriptase enzyme (#EP0742, Thermo Fisher Scientific) according to manufacturer's instructions. RT-qPCR was used to analyze the expression levels of microglia genes, *P2RY12, TREM2, TLRM2, CLEC7A, APOE,* and *PLCG2* by using Maxima Probe/ROX qPCR Master Mix (#11813923, Thermo Fisher Scientific) and the Taqman primers listed in **Supplementary table 1** on Bio-Rad CFX96 Real-Time System (Biorad).

The relative mRNA expression results were normalized to GAPDH using Q-gene program (DOI: 10.1093/bioinformatics/btg157 ).

**Table 1:** Primers used for characterization of microglia markers in iMGL cells.

| Primers | Producer | Catalog number |
|---|---|---|
| P2RY12 | Thermo Fisher Scientific | Hs00375457_m1 |
| TREM2 | Thermo Fisher Scientific | Hs00219132_m1 |
| TLR2 | Thermo Fisher Scientific | Hs00152932_m1 |
| CLEC7A | Thermo Fisher Scientific | Hs00224028_m1 |
| APOE | Thermo Fisher Scientific | Hs00171168_m1 |
| PLCG2 | Thermo Fisher Scientific | Hs00182192_m1 |
| GAPDH | Thermo Fisher Scientific | Hs99999905_m1 |

**Supplementary figure 1. Representative ROIs used for analyzing IBA1, GFAP, 22C11, and APOE within and surrounding β-amyloid plaques.** The inner circle was calculated based on the plaque size and width of the outer circle(s) were kept constant within each analysis. Analysis of IBA1 and GFAP within 0-20 and 20-40 µm (A-B), 22C11 within 0-14 µm (C), and APOE within 0-20 µm (D) from the plaque outline was conducted.

**Supplementary figure 2. PLCγ2-P522R variant does not change β-amyloid load in the hippocampus of APP/PS1 mice.** A) β-amyloid plaque count, coverage (total plaque area, % of whole analyzed area, $\mu m^2$), and size ($\mu m^2$) of the individual plaques in the hippocampus of the APP/PS1xPlγ2-P522R (A+/$P^{ki/ki}$) and APP/PS1 (A+/$P^{wt/wt}$) mice. n(A+/$P^{wt/wt}$)=4, n(A+/$P^{ki/ki}$)=6. B) Insoluble, but not soluble Aβ40 and -42 levels are slightly, but not significantly lower in the hippocampus of the A+/$P^{ki/ki}$ mice as compared to the A+/$P^{wt/wt}$ mice. Aβ40 and -42 levels are normalized to the total protein concentration in the same sample. n(A+/$P^{wt/wt}$)=5, n(A+/$P^{ki/ki}$)=6. C) Representative Western blots and corresponding quantification show no differences in the levels of full-length APP (APPtot, normalized to GAPDH), APP C-terminal fragments (C99 and C83, normalized to APPtot), or soluble APPα and –β (sAPPα, sAPPβ, normalized to APPtot) species in the hippocampus of the $A+/P^{ki/ki}$ and the $A+/P^{wt/wt}$ mice. n($A+/P^{wt/wt}$)=4, n($A+/P^{ki/ki}$)=4. Scale bars in the representative immunofluorescent images are 657 µm for the whole area and 164 µm for the zoomed view. Unpaired T-test. All data are presented as mean ± SEM. Each datapoint represents an individual mouse.

**Supplementary figure 3. PLCγ2-P522R variant increases microglia clustering around β-amyloid plaques in the hippocampus of APP/PS1 mice.** A) Area ($\mu m^2$) of IBA1-positive microglia is increased within 0-20 µm (*p=0.023) and 0-40 µm (*p=0.037) from the plaque outline in the hippocampus of the $A+/P^{ki/ki}$ mice as compared to the $A+/P^{wt/wt}$ mice. Simultaneously, total IBA1 area is decreased (*p=0.043). n($A+/P^{wt/wt}$)=4, n($A+/P^{ki/ki}$)=6. B) GFAP-positive astrocyte area ($\mu m^2$) around plaques (within 0-20 and 20-40 µm from the plaque outline) and total GFAP area remain unaltered between the genotypes. n($A+/P^{wt/wt}$)=5, n($A+/P^{ki/ki}$)=6. The scale bar in the representative immunofluorescent images is 50 µm. Unpaired samples T-Test. All data are presented as mean ± SEM. Each datapoint represents an individual mouse.

**Supplementary figure 4. PLCγ2-P522R variant decreases dystrophic neurites around β-amyloid plaques in the hippocampus of APP/PS1 mice.** A) Total area ($\mu m^2$) of diffuse Aβ (6E10) and composition β-amyloid plaques, as indicated by percentage of compact amyloid (X-34) of all β-amyloid (X-34+6E10), remain unaltered in the hippocampus of the APP/PS1xPlγ2-P522R ($A+/P^{ki/ki}$) mice as compared to the APP/PS1 ($A+/P^{wt/wt}$) mice. n($A+/P^{wt/wt}$)=5, n($A+/P^{ki/ki}$)=6. B) A 3D-reconstruction of 6E10 and IBA1 (microglia) signal and their co-localization and quantification showing 6E10 within IBA1 as % of all 6E10 in the hippocampus of $A+/P^{ki/ki}$ mice as compared to the

A+/$P^{wt/wt}$ mice. C) Area (µm$^2$) of 22C11-labeled dystrophic neurites around β-amyloid plaques strongly correlates with plaque size (r=0.569, p<0.0001). 22C11 area is decreased within 0-7 µm (*p=0.01) and 0-14 µm (*p=0.01) distance from the β-amyloid plaque outline in the hippocampus of the A+/$P^{ki/ki}$ mice as compared to the A+/$P^{wt/wt}$ mice when normalized to the plaque size. n(A+/$P^{wt/wt}$)=4, n(A+/$P^{ki/ki}$)=6. The scale bar in the representative immunofluorescent images is 50 µm. Unpaired T-test and Pearson correlation. All data are presented as mean ± SEM. Each datapoint represents an individual mouse.

**Supplementary figure 5. PLCγ2-P522R variant does not change plaque-associated APOE in the hippocampus of APP/PS1 mice.** A) Analysis of total APOE or APOE area (µm$^2$) on top and surrounding (within 0-10 and 10-20 µm from the β-amyloid plaque outline) β-amyloid plaques reveals no differences between the APP/PS1xPlγ2-P522R (A+/$P^{ki/ki}$) and APP/PS1 (A+/$P^{wt/wt}$) mice. n(A+/$P^{wt/wt}$)=5, n(A+/$P^{ki/ki}$)=6. The scale bar in the representative immunofluorescent images is 50 µm. Unpaired samples T-Test. All data are presented as mean ± SEM. Each datapoint represents an individual mouse. B) Volcano blot showing differentially expressed genes (DEGs) in parieto-occipital cortex and C) CD11b+ microglia isolated from A+/$P^{wt/wt}$ and A+/$P^{ki/ki}$ mouse brain.

**Supplementary figure 6. Differentially expressed genes and proteins in CD11b+ microglia of WT and PLCγ2-P522R KI mice.** Volcano plot of A) differentially expressed genes (DEGs) and B) differentially expressed proteins (DEPs) in CD11b+ microglia isolated from 13-month-old PLCγ2-P522R KI and WT mice. FDR<0.05. Horizontal dashed line: adjusted p-value 0.05; vertical dashed lines: |log2FC|=0.1. C) PLCγ2-P522R downregulates calcium-sensitive transcription family members, *Mef2a* (****pAdj<0.0001), *Mef2c* (****pAdj<0.0001), and *Mef2d* (*pAdj=0.015) as compared to WT microglia. Expression of genes encoding fatty acid binding proteins *Fabp7* (****pAdj<0.0001), *Fabp5* (pAdj=0.003), and *Fabp3* (*pAdj=0.049) is increased in PLCγ2-P522R KI microglia. Expression of several genes encoding phosphatidylinositol 3-kinase (PI3K) subunits, *Pik3cd* (****pAdj<0.0001), *Pik3cg* (***pAdj=0.0003), *Pik3r5* (**pAdj=0.002), *Pik3ap1* (**pAdj=0.003), *Pik3r1* (*pAdj=0.012), *Pik3r4* (*pAdj=0.033), is downregulated in PLCγ2-P522R KI as compared to WT microglia. D) A dot plot of normalized enrichment scores for enriched and depleted protein sets in enrichment analyses by GSEA for protein (Wikipathways) expression in PLCγ2-P522R KI and WT mice. RNA n(WT)=3, n(PLCγ2-P522R KI)=4, protein n(WT)=5, n(PLCγ2-P522R KI)=6

**Supplementary figure 7. Characterization of human microglia-like cell models.** A) IL-6 and TNF-α levels in the conditioned medium of blood monocyte-derived microglia (MDMi) of the PLCγ2-P522R variant carriers (CG) and matched controls (CC) after treatment with lipopolysaccharide for 24 h. IL-6 and TNF-α levels are normalized to the total protein concentration in the respective lysate. n(CC)=3, n(CG)=4. Each datapoint represents data from one individual. Colored circles indicate data obtained from females and hollow circles data obtained from males. Microglial markers in induced pluripotent stem cell-derived microglia (iMGL) detected by immunocytochemistry (B) and RT-qPCR (C). In (B) scale bars are 50µm. Data are from PLCγ2 control line (CC) and isogenic PLCγ2-P522R homozygous line (GG) with either *APOE33* or *APOE44* background. In (C) there were two batches of cells, two technical replicates each. All data are presented as mean ± SEM.

**Supplementary figure 8.** Uncut Western blot images of APP and its metabolites in A) parieto-occipital cortex and B) hippocampus of the APP/PS1xPlγ2-P522R (A+/$P^{ki/ki}$) and APP/PS1 (A+/$P^{wt/wt}$) mice. Red X indicates a sample unrelated to the project.

Supplementary figure 1

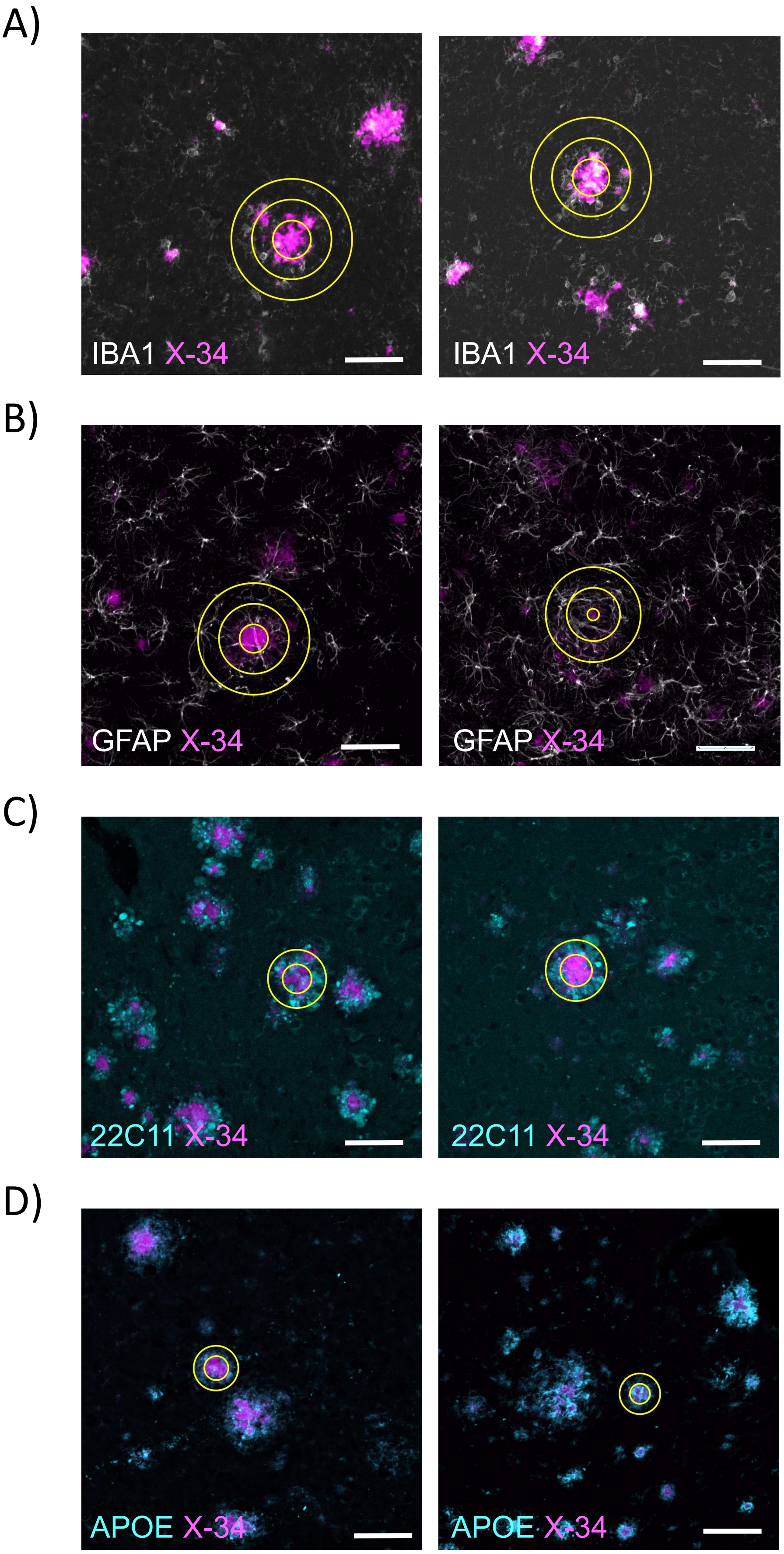

Supplementary figure 2

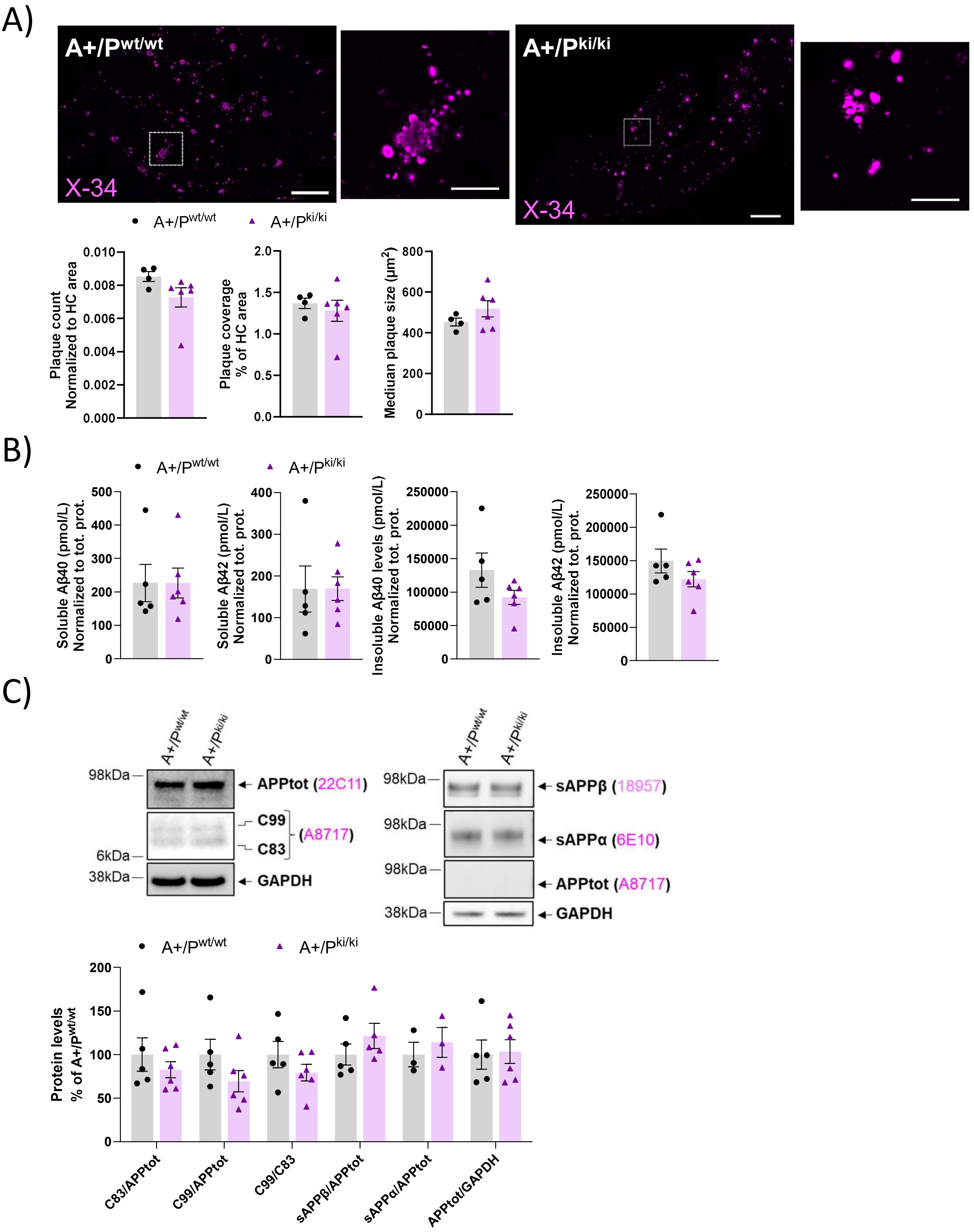

### Supplementary figure 3

A)

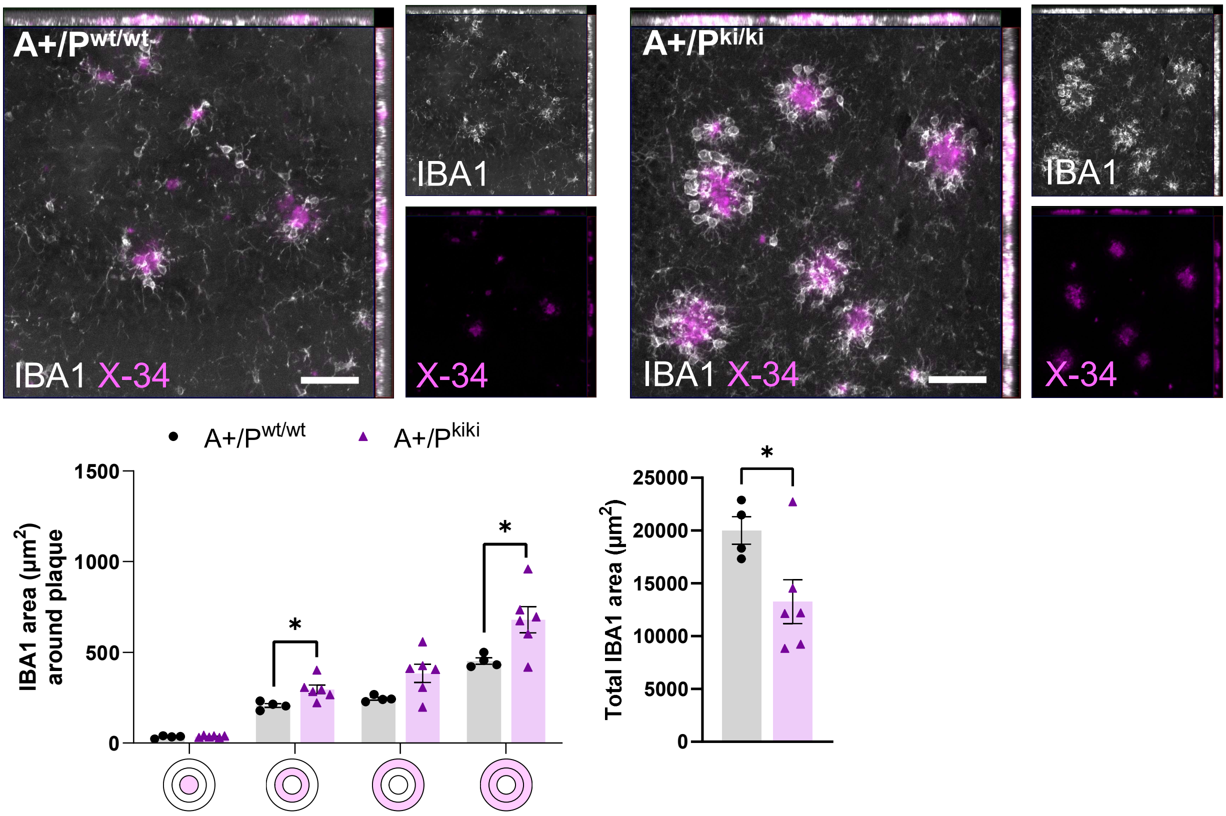


B)

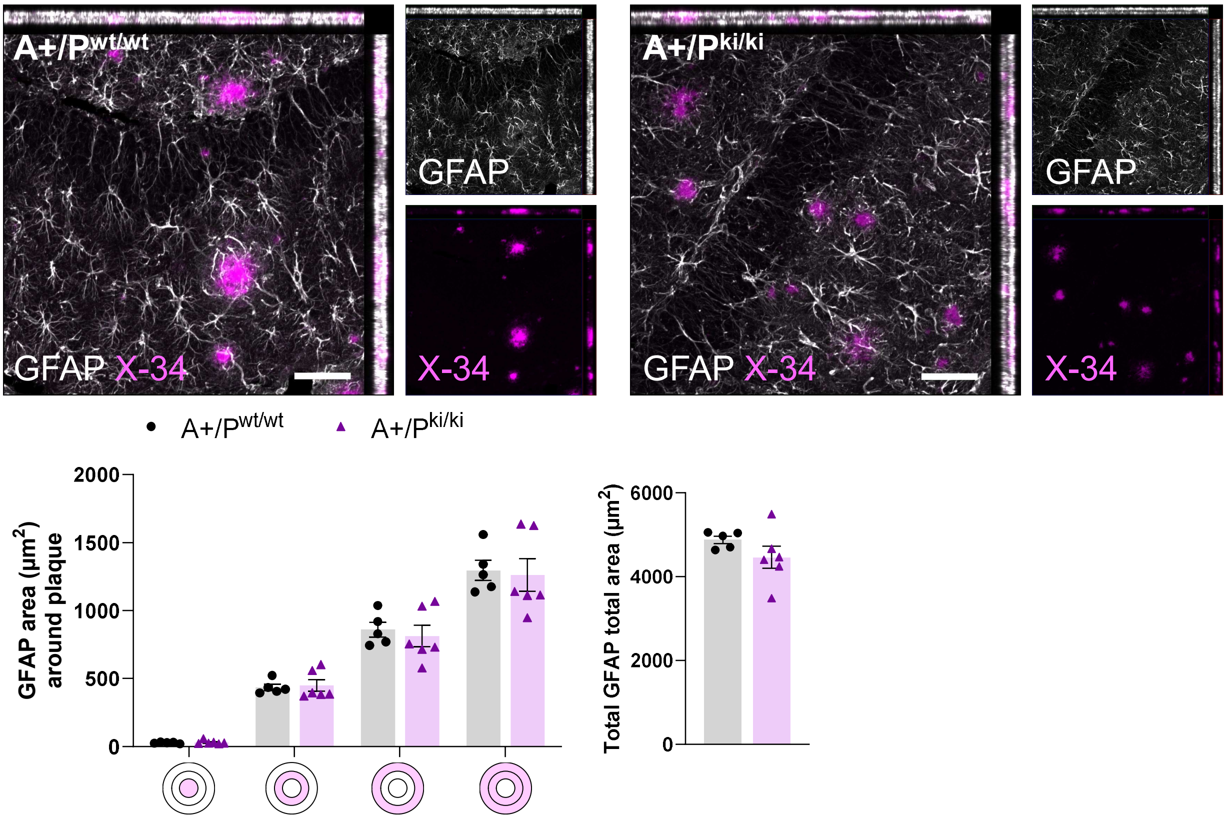

### Supplementary figure 4

A)

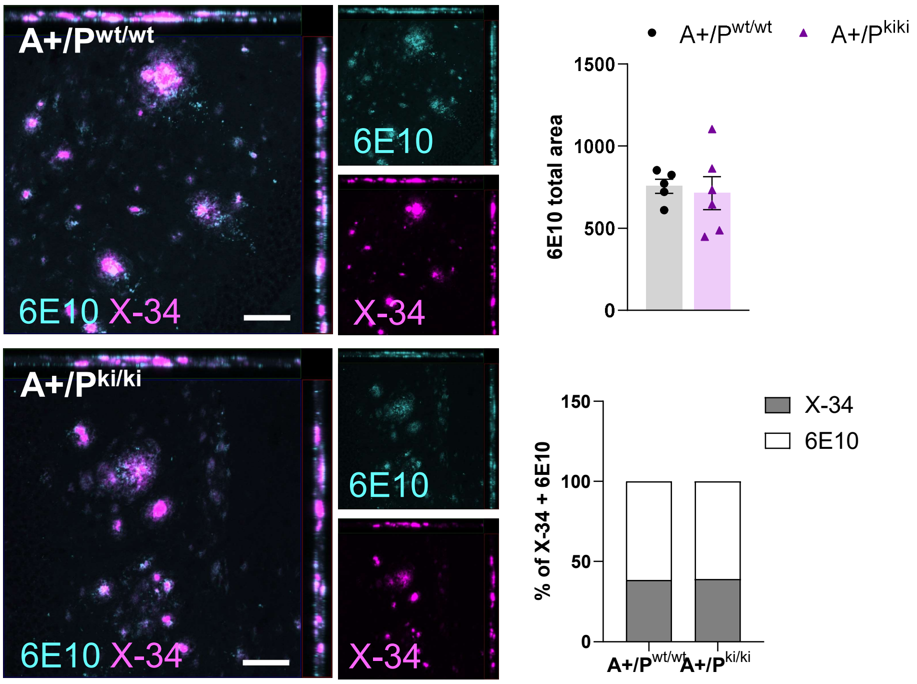


B)

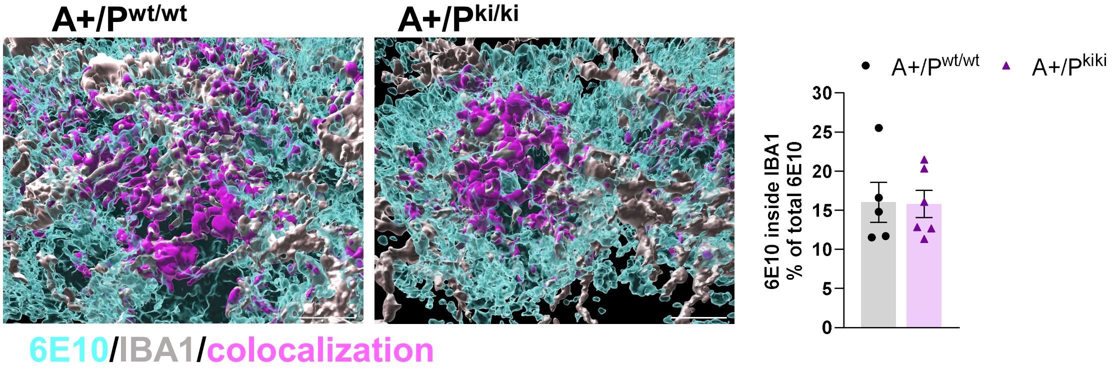


C)

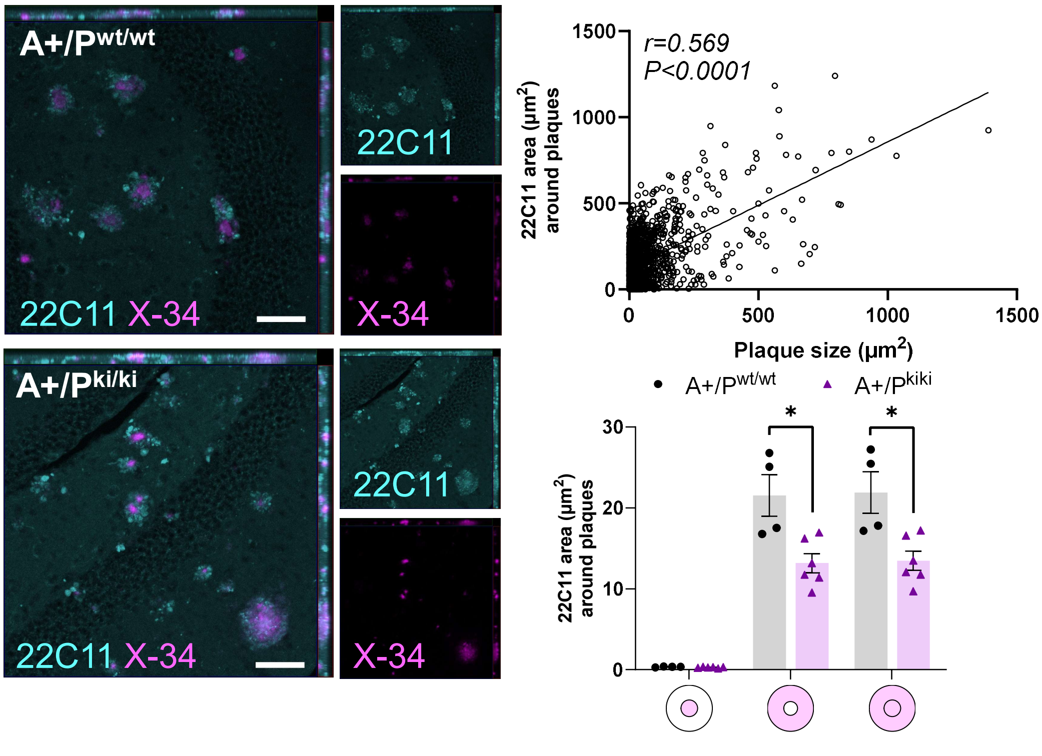

### Supplementary figure 5

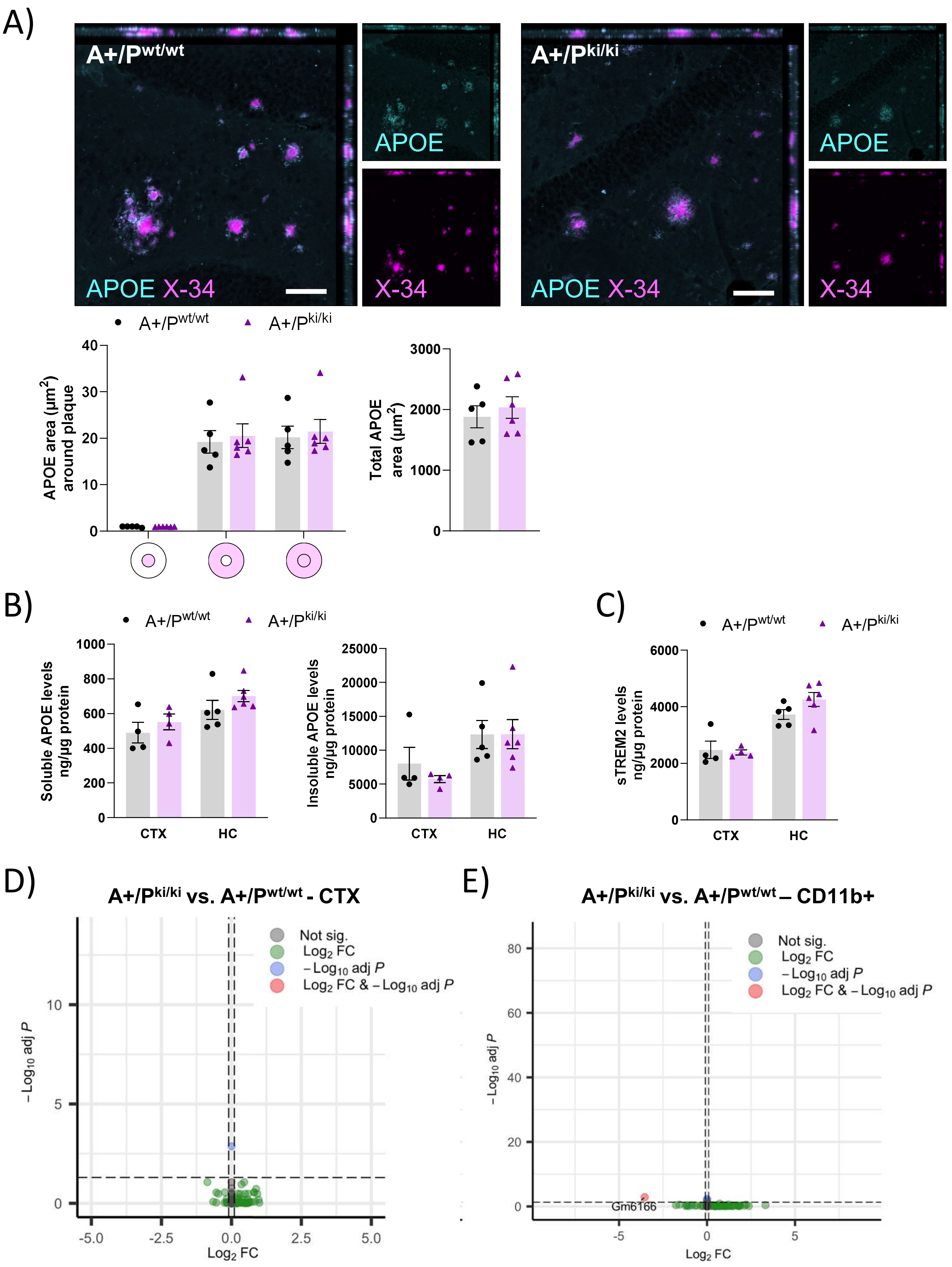

### Supplementary figure 6

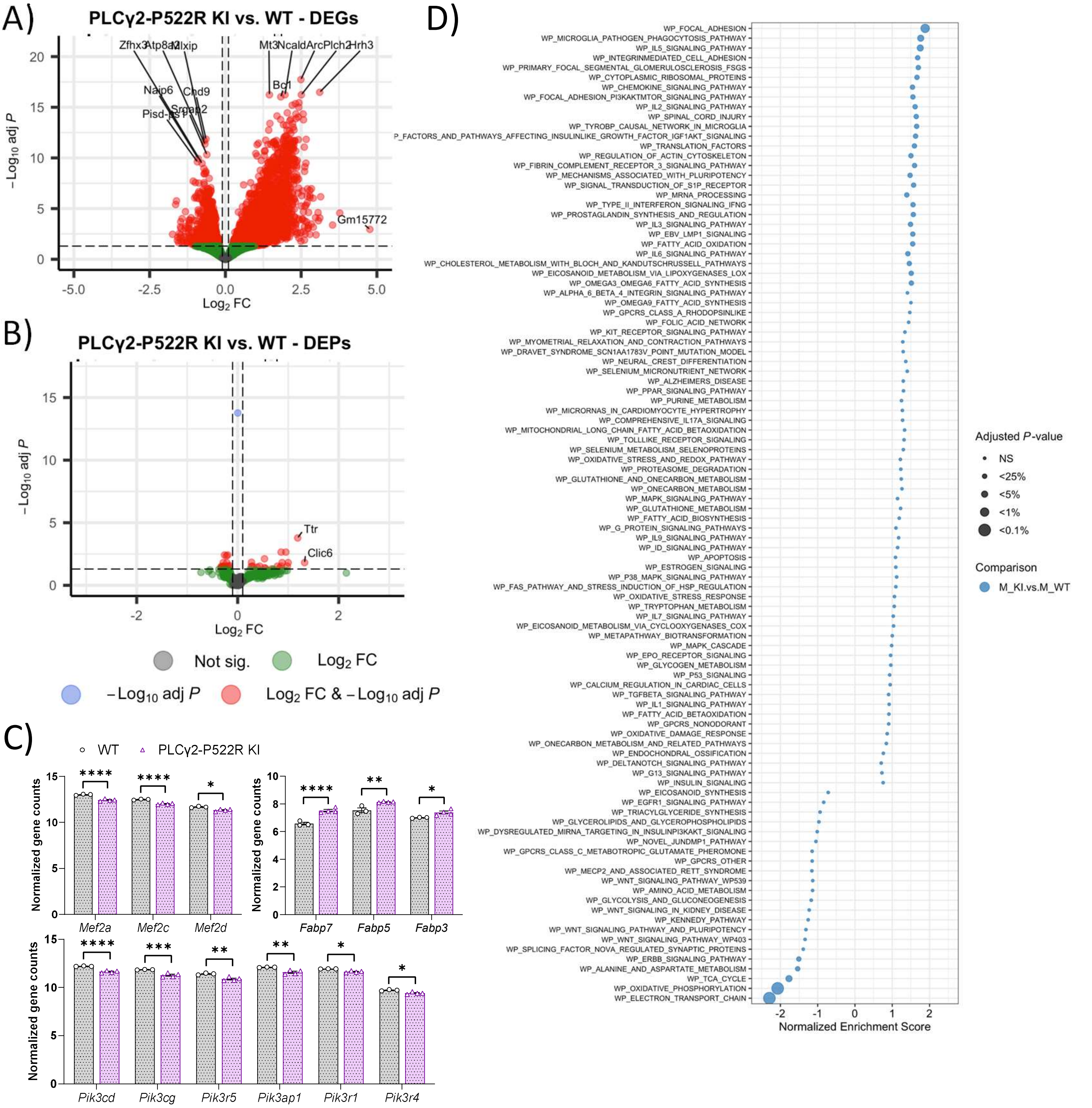

### Supplementary figure 7

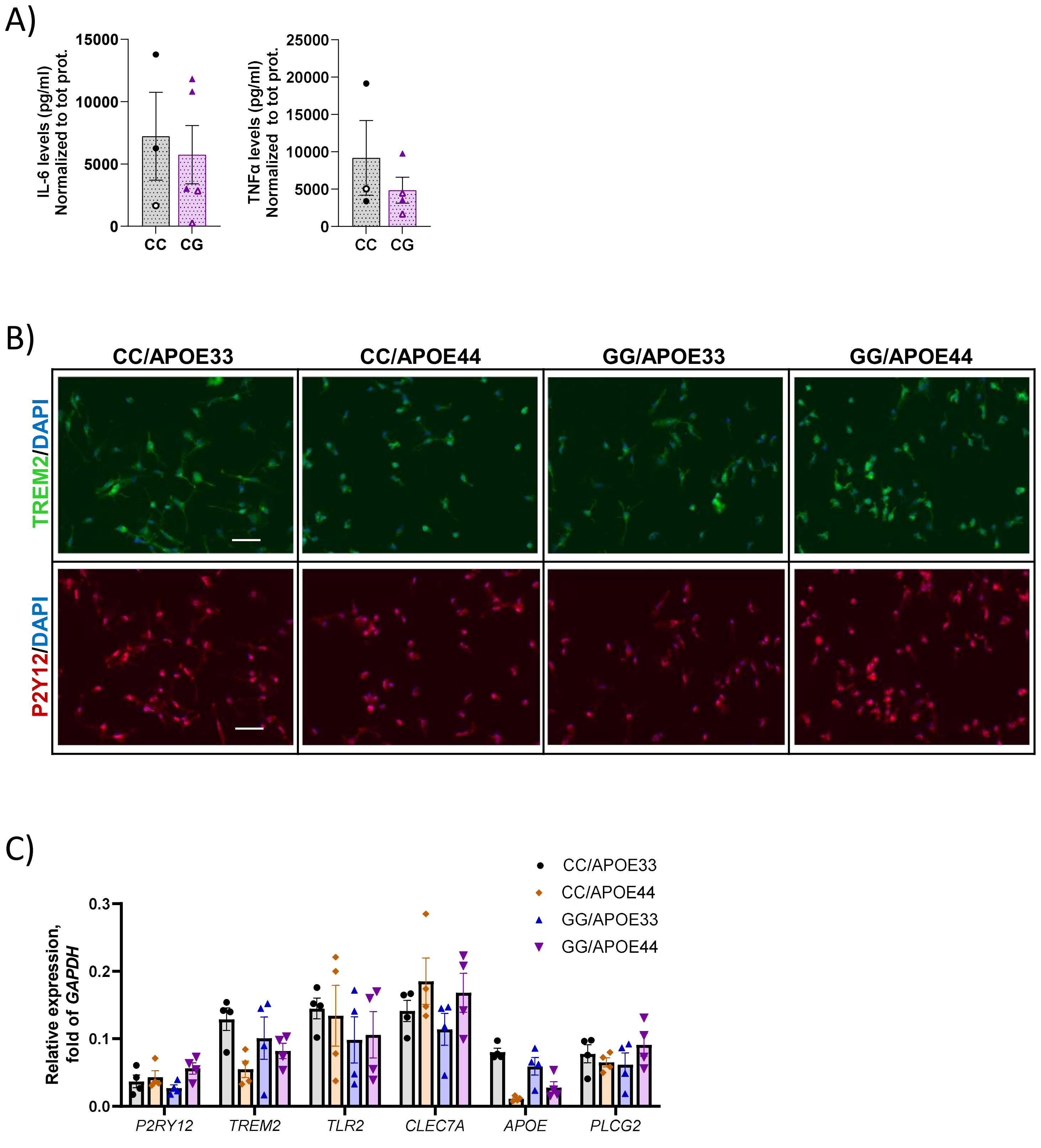
